# Scaling pattern of the carnivoran forelimb: Locomotor types, differential scaling and thoughts on a dying similarity

**DOI:** 10.1101/2022.06.29.498091

**Authors:** Eloy Gálvez-López, Adrià Casinos

## Abstract

The scaling pattern of the forelimb in Carnivora was determined using a sample of 30 variables measured on the scapula, humerus, radius, ulna, and third metacarpal, of 429 specimens belonging to 137 species of Carnivora. Standardized major axis regressions on body mass were calculated for all variables, using both traditional regression methods and phylogenetically independent contrasts (PIC). In agreement with previous studies on the scaling of the appendicular skeleton, conformity to either the geometric similarity hypothesis or the elastic similarity hypothesis was low. The scaling pattern of several phyletic lines and locomotor types within Carnivora was also determined, and significant deviations from the scaling pattern of the order were found in some of these subsamples. Furthermore, significant evidence for differential scaling was found for several variables, both in the whole sample and in various phylogenetic and locomotor subsamples. Contrary to previous studies, significant differences were found between the allometric exponents obtained with traditional and PIC regression methods, emphasizing the need to take into account phylogenetic relatedness in scaling studies. In light of these and previous results, we conclude that similarity hypotheses are too simplistic to describe scaling patterns in the carnivoran appendicular skeleton, and thus we propose that scaling hypotheses should be built from similarities in the scaling patterns of phylogenetically narrow samples of species with similar locomotor requirements. The present work is a first step in the study of those samples.

## Introduction

Size is one of the most important factors affecting the shape and function of the elements of the musculoskeletal system of animals, as well as the parameters defining their locomotor dynamics (e.g. duty factor) (Schmidt-Nielsen, 1984; Alexander, 2002; Biewener, 2003). Thus, several hypotheses have been proposed to predict how these musculoskeletal elements and locomotor parameters would be affected by variations in body size (i.e., scaling). The most widespread of these similarity hypotheses are the geometric similarity hypothesis (already supported by Hill, 1950) and the elastic similarity hypothesis (proposed by McMahon, 1973). The former states that all linear measurements of an organism are proportional to its body mass^0.33^, while according to the latter, lengths scale to body mass^0.25^ and diameters to body mass^0.375^.

In the case of skeletal measurements, early studies suggested that geometric similarity (GS) explained their scaling in mammals (e.g. Alexander et al., 1979), while elastic similarity (ES) was only found in Bovidae (McMahon, 1975*a*; Alexander, 1977). However, as the number of studies in this subject increased, empirical evidence showed that conformity to either hypotheses was low in mammals (Bou et al., 1987; Bertram & Biewener, 1990; Christiansen, 1999*a,b*; Carrano, 2001; Llorens et al., 2001; Lilje et al., 2003; Casinos et al., 2012). Furthermore, in some cases it has been found that the same skeletal measurement scaled geometrically in small species and elastically in large species (Economos, 1983; Bertram & Biewener, 1990; Silva, 1998; Christiansen, 1999*a,b*; Carrano, 2001). This differential scaling (or complex allometry) suggests that general allometric calculations would thus not be applicable to a large range of variations in body size.

Despite the large number of studies on the scaling of the mammalian appendicular skeleton, little to no consideration has been given to the scaling of skeletal measurements other than the length and diameters of the humerus, radius/ulna, femur and tibia. In fact, only the study of Lilje et al. (2003) on Ruminantia and that of Schmidt & Fischer (2009) on Mammalia have paid any attention to the scaling of the scapula, which has been shown to be the main propulsive element of the forelimb (Lilje & Fischer, 2001; Fischer et al., 2002). Furthermore, although several studies have dealt with the scaling of particular orders within Mammalia, their sample sizes are usually too small to perform interfamilial comparisons. Finally, no work so far has studied how locomotor specializations affect the scaling pattern of the appendicular skeleton in a comparative framework. It has been suggested that similarity hypotheses imply adaptive neutrality, or at least independence of the locomotor type of the species that are compared (Bou et al., 1987). Therefore, samples including extreme locomotor patterns should deviate markedly from the predictions of similarity hypotheses. The order Carnivora is one of the few groups of mammals that allows an allometric study of the appendicular skeleton in such a multifaceted approach, since: 1) carnivorans span a size range of four orders of magnitude (from less than 0.1 kg in the least weasel (*Mustela nivalis*) to well over two tonnes in elephant seals (*Mirounga sp*.)), which enables not only classic allometric studies but also to test for differential scaling; 2) they constitute a monophyletic group with several well-represented families, granting interfamilial scaling comparisons; and 3) they present one of the widest locomotor diversities among mammals, which allows to study the effect of locomotor specializations in the scaling of the limb bones (Van Valkenburgh, 1987; Bertram & Biewener, 1990; Wilson & Mittermeier, 2009; Nyakatura & Bininda-Emonds, 2012).

Thus, the first aim of this study was to determine the scaling pattern of the carnivoran appendicular skeleton, with emphasis on the scapula and several morphofunctional dimensions of the appendicular skeleton, and to assess whether differential scaling could be found in this pattern. Previous studies on the scaling of the appendicular skeleton in Carnivora have shown low conformity to either similarity hypothesis when long bone lengths are regressed against diameters (Bertram & Biewener, 1990). However, when regressed against body mass, bone lengths tend to scale geometrically and least circumference elastically (Christiansen, 1999*a*). More recently, two studies on the scaling of relative segment lengths in Mammalia have also presented separate results for the carnivoran species in their sample. However, while first Schmidt (2008) suggested that limb proportions are size-independent in Carnivora, significant size-related variation in those variables was later found by the same author (Schmidt & Fischer, 2009). Finally, regarding differential scaling, Bertram & Biewener (1990) found evidence for complex allometry in the length and diameters of the carnivoran humerus, radius, femur and tibia.

Once this scaling pattern for the whole order was determined, the second aim of this study was to analyze whether the main phyletic lines (families) within Carnivora deviated from it, and if so, then how. Few scaling studies have been carried out on the appendicular skeleton of any particular family within Carnivora. When regressing long bone lengths and diameters to femur length in Canidae, Wayne (1986) found significant deviations from isometric scaling, which suggested low conformity with either GS or ES in the appendicular skeleton of canids. However, in a study with over sixty dog breeds, Casinos et al. (1986) found that the scaling of humerus, radius and tibia conformed to GS but not that of the femur, which could explain the lack of conformity in Wayne’s study. Heinrich & Biknevicius (1998) showed that, in Martinae (Mustelidae), long bone dimensions tended to scale elastically, but conformity was also low. Recent studies suggest geometric scaling with no differential scaling in Felidae (Day & Jayne, 2007; Gálvez-López & Casinos, 2012). Finally, evidence for scaling differences between Felidae and Canidae was presented by Meachen-Samuels & Van Valkenburgh (2009).

The last objective of the present study was to test whether particular locomotor habits within Carnivora cause deviations from the general scaling pattern for the order. To our knowledge, only the study of Bou et al. (1987) has pursued a similar approach, but then in rodents and other small mammals. In the case of Carnivora, this lack of studies could be related to the general belief that their appendicular skeleton is highly conservative in terms of bone morphology and locomotor style (Flynn et al., 1988; Bertram & Biewener, 1990; Day & Jayne, 2007; but see Heinrich & Biknevicious, 1998; Gálvez-López, 2021).

## Material and Methods

The sample consisted of 429 specimens from 137 species of Carnivora (Table 1), representing about 48% of extant species (Wozencraft, 2005). Table 2 describes the locomotor categories used in this study, which represent the locomotor specialization of each species (i.e., the main locomotor habit of each species). For each specimen, measurements were taken on the scapula, humerus, radius, ulna, and third metacarpal. The variables analyzed in this study have already been described in the Supplementary Information of Gálvez-López (2021) but are repeated here in Table 3 for simplicity. The 30 studied variables included 19 linear measurements, one projected distance (**T**), 8 ratios, and 2 angles (θ, α), and are summarized in Figure 1.

**Table 1.**
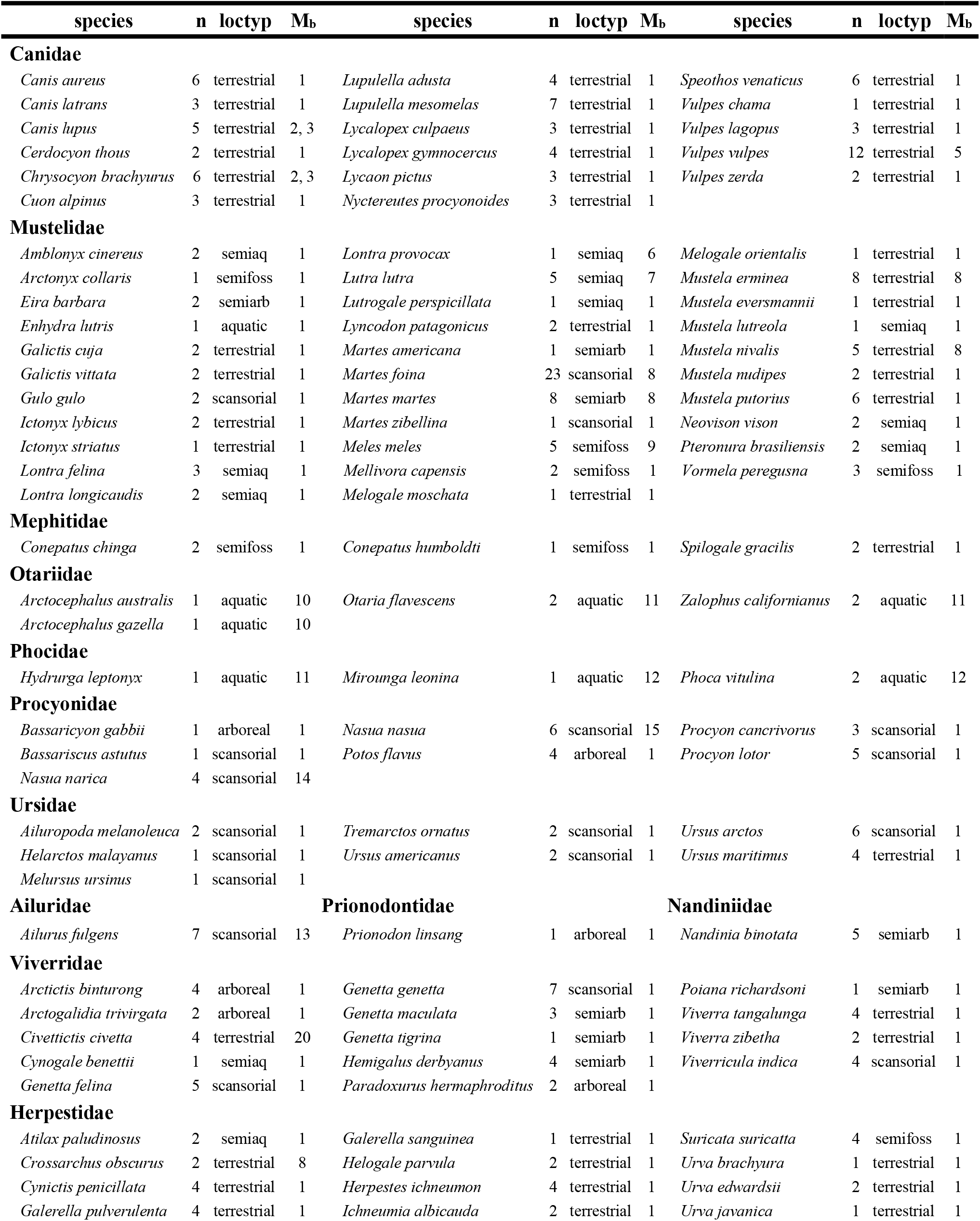
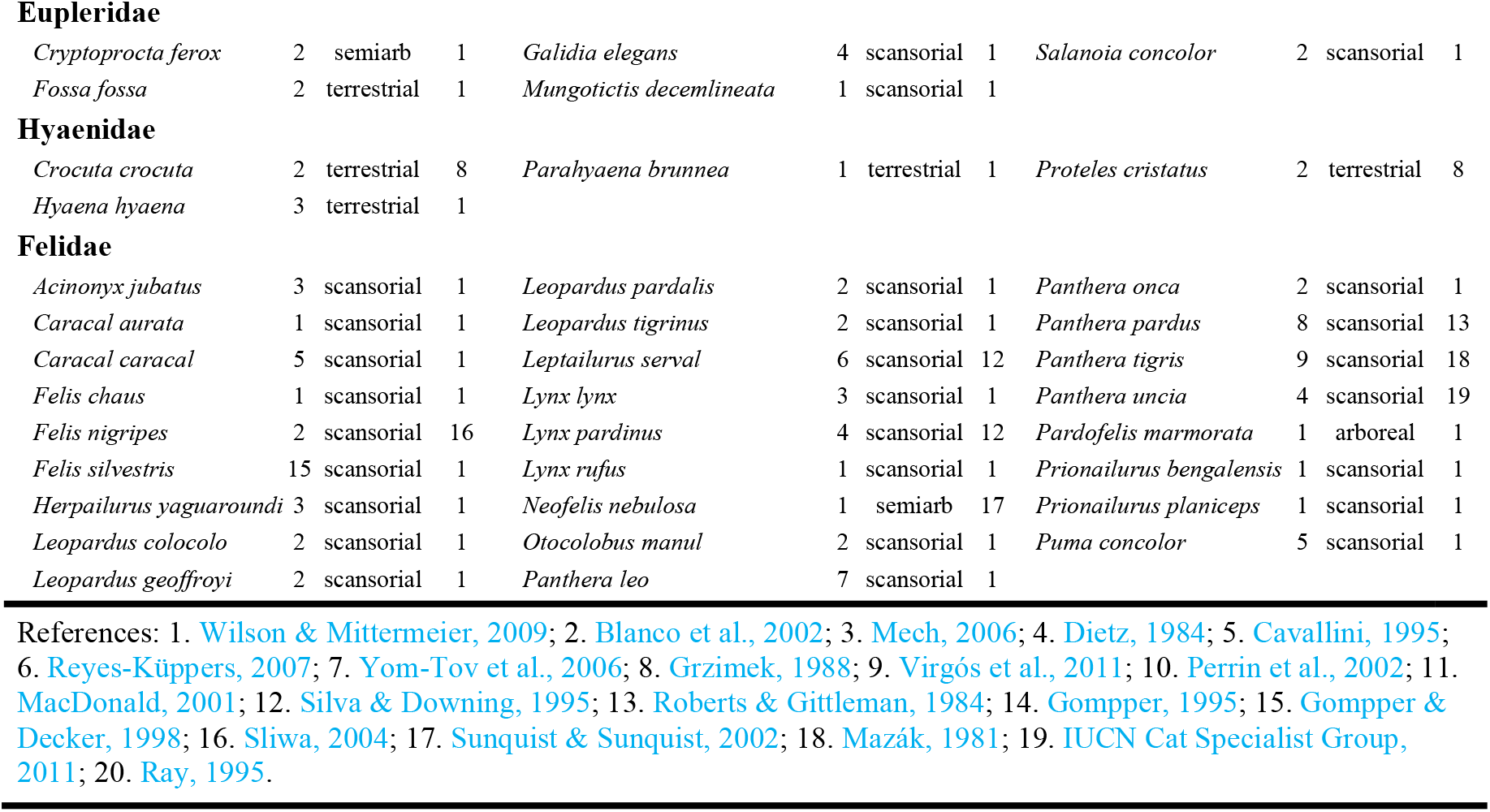
Species measured. For each species, the table shows the number of measured specimens (n), the assigned category for locomotor type (loctyp), and the references from which the mean body mass value for that species was taken (**M**_**b**_). See Table 2 for a description of locomotor type categories. Abbreviations: semiaq, semiaquatic; semiarb, semiarboreal; semifoss, semifossorial.

**Table 2.** Locomotor type categories. Locomotor type categories were adapted from previous works on the relatioship between locomotor behavior and forelimb morphology (Eisenberg, 1981; Van Valkenburgh, 1985, 1987).

| Locomotor type | Description |
| --- | --- |
| arboreal | species that spend most of their life in trees (over 75%), rarely descending to the ground |
| semiarboreal | species that spend a large amount of their time in the trees (between 50% and 75%), both foraging and resting, but also on the surface of the ground |
| scansorial | species that, although mostly terrestrial (over half their time is spent on the ground), can climb well and will readily do so to chase arboreal prey or escape |
| terrestrial | species that rarely or never climb or swim, and that may dig to modify a burrow but not regularly for food |
| semifossorial | species that dig regularly for both food and shelter, but that still show considerable ability to move about on the surface |
| semiaquatic | species that forage regularly underwater and usually plunge into the water to escape, but must spend time ashore to groom,... |
| aquatic | species that carry out most of their life cycle in water, although some part of it can be confined to land (parturition, mating, rearing the young) |

**Table 3.** Variable names and abbreviations. Two subsamples can be defined within the studied variables: linear measurements (dark grey) and ratios and angles (light grey). For each variable, it is also indicated which table in the Supplementary Materials shows the regression results.

| Name | Abbr. | Table |
| --- | --- | --- |
| Body mass | M<sub>b</sub> |  |
| Scapular length | L<sub>s</sub> | SR1, 31 |
| Maximum width of supraspinous fossa | S | SR2, 32 |
| Maximum width of infraspinous fossa | I | SR3, 33 |
| Maximum scapular width | A | SR4, 34 |
| Scapular spine height | H<sub>s</sub> | SR5, 35 |
| Humerus functional length | L<sub>h</sub> | SR6, 36 |
| Humerus sagittal diameter | d<sub>sh</sub> | SR7, 37 |
| Humerus transverse diameter | d<sub>th</sub> | SR8, 38 |
| Projected height of greater tubercle | T | SR9 |
| Humerus robusticity | HR | SR10, 39 |
| Radius functional length | L<sub>r</sub> | SR11, 40 |
| Radius sagittal diameter | d<sub>sr</sub> | SR12, 41 |
| Radius transverse diameter | d<sub>tr</sub> | SR13, 42 |
| Styloid process length | P | SR14, 43 |
| Radius robusticity | RR | SR15, 44 |
| Ulna functional length | L<sub>u</sub> | SR16, 45 |
| Ulna sagittal diameter | d<sub>su</sub> | SR17, 46 |
| Ulna transverse diameter | d<sub>tu</sub> | SR18, 47 |
| Olecranon process length | O | SR19, 48 |
| Olecranon angle | α | SR20, 49 |
| Olecranon abduction angle | θ | SR21, 50 |
| Ulna robusticity | UR | SR22, 51 |
| Indicator of Fossorial Ability | IFA | SR23, 52 |
| Third metacarpal functional length | L<sub>m</sub> | SR24, 53 |
| Third metacarpal sagittal diameter | d<sub>sm</sub> | SR25, 54 |
| Third metacarpal transverse diameter | d<sub>tm</sub> | SR26, 55 |
| Third metacarpal robusticity | MR | SR27, 56 |
| Relative length of the proximal segment of the forelimb | %<sub>prox</sub> | SR28, 57 |
| Relative length of the middle segment of the forelimb | %<sub>mid</sub> | SR29, 58 |
| Relative length of the distal segment of the forelimb | %<sub>dist</sub> | SR30, 59 |

**Figure 1.**
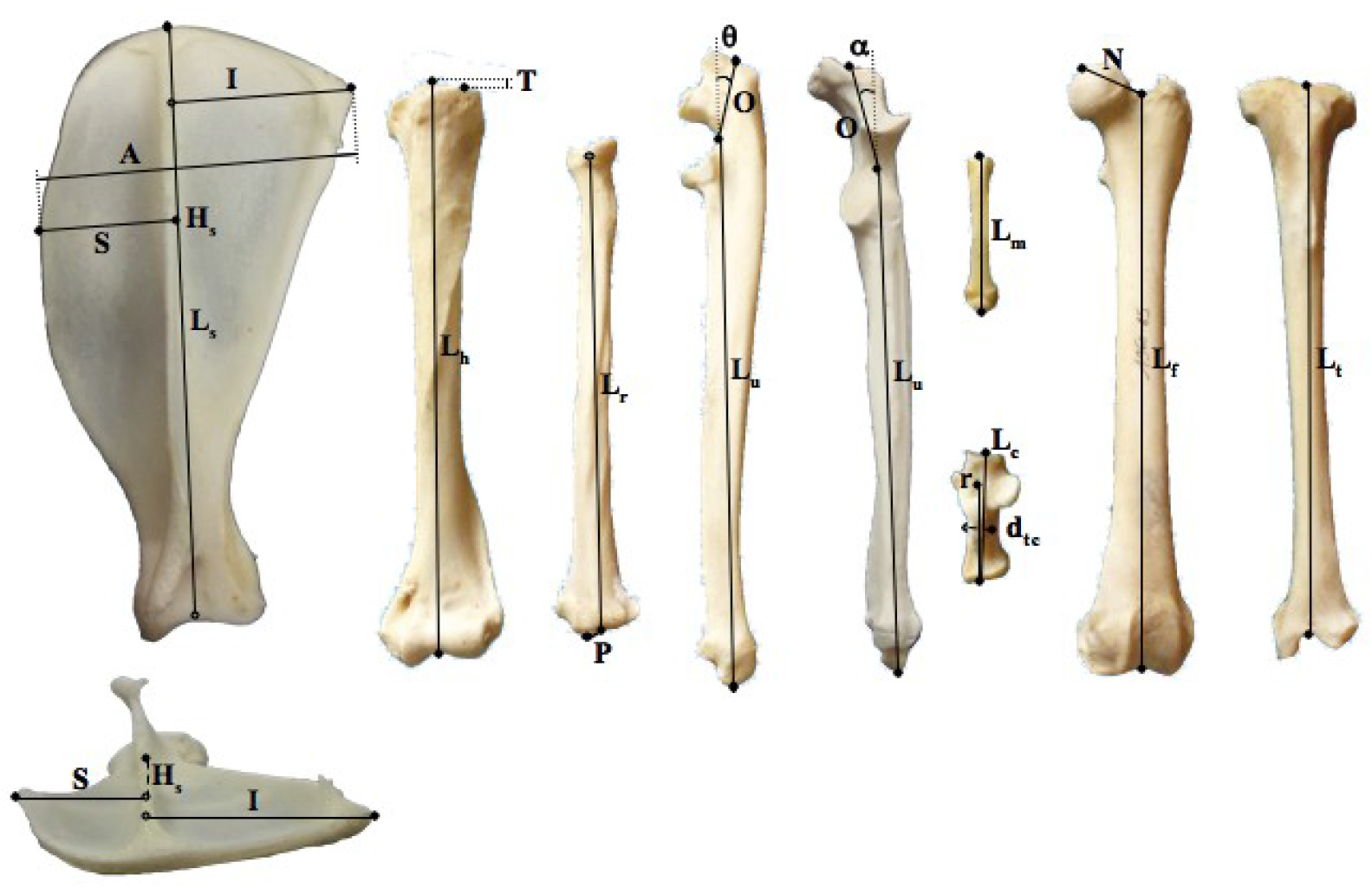
Bone measurements. All other variables were calculated from these measurements. See Gálvez-López (2021) for a more detailed definition of each variable, and an assessment of measurement error. Variable names are listed in Table 3.

Regression methods were used to relate each variable to body mass (**M**_**b**_). All regressions were calculated with the standardised major axis method (SMA), since regression slopes were the primary interest of this study, and ordinary least squares regression methods (OLS) tend to understimate the slope of the line-of-best-fit because its calculation involves fitting the predicted y-values as closely as possible to the observed y-values (Warton et al., 2006). The power equation (***y*** = *a* · ***x***^*b*^ ; Eq. 1) was assumed for all variables but **T** and θ, for which the linear model of regression was used (***y*** = *a* + *b* · ***x*** ; Eq. 2), and 95% confidence intervals were calculated for both the coefficient (*a*) and the allometric exponent (*b*_*trad*_). All regressions were calculated using PAST (Hammer et al., 2001). In order to compare the present results with those previously published using OLS regressions, SMA slopes were calculated for those studies prior to the comparison by dividing their OLS slopes by the corresponding correlation coefficient (Sokal & Rohlf, 1995).

Additionally, all the SMA regression slopes were also calculated using phylogenetically independent contrasts (PIC; Felsenstein, 1985). This methodology takes into account the phylogenetic signal inherent to interspecific data and thus accounts for the potential correlation of the error terms that could arise due to the lack of independence among species, since they can be arranged in a hierarchical sequence (i.e., a phylogenetic tree; Felsenstein, 1985; Grafen, 1989; Harvey & Pagel, 1991; Christiansen, 2002*a, b*). PIC regression slopes (*b*_*PIC*_) were calculated using the PDAP: PDTREE module of Mesquite (Maddison & Maddison, 2010; Midford et al., 2010). The structure of the phylogenetic tree used in this study is discussed and detailed in the Supplementary Information of Gálvez-López (2021), but is reproduced here in Figure 2. When necessary, branch lengths were transformed in order to obtain a low and non-significant correlation between the standardized value of the PIC contrasts and their corresponding standard deviation. This process has proven to be a good solution against possible violations of the assumptions implied by PIC methodology (Felsenstein, 1985; Grafen, 1989; Díaz-Uriarte & Garland, 1996, 1998).

**Figure 2.**
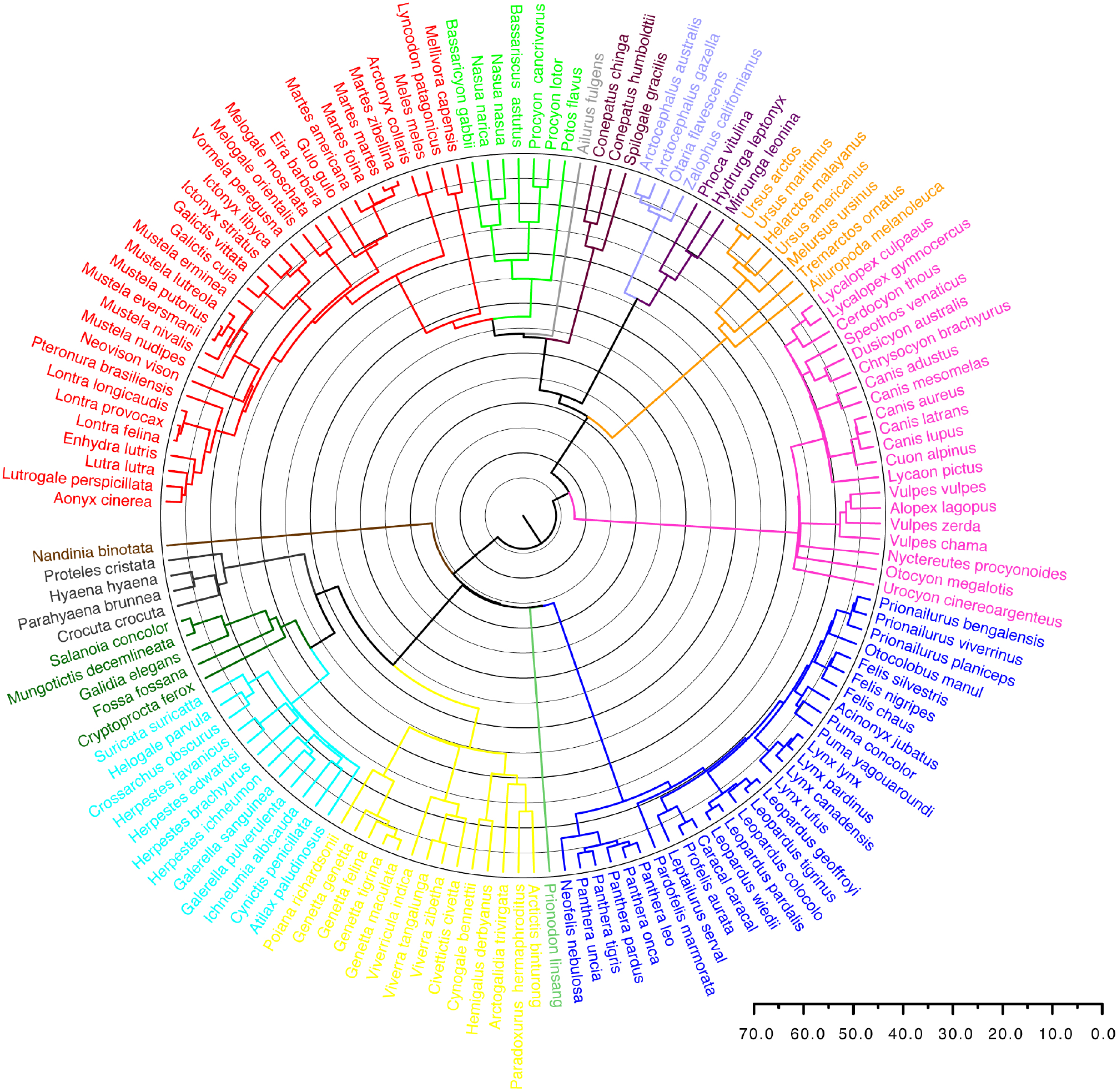
Phylogenetic relationships among the species of Carnivora used in this study. The timescale represents divergence times in millions of years. The phylogeny shown was modified after Nyakatura & Bininda-Emonds (2012), as described in Gálvez-López (2021, Supplementary Information).

For each variable and methodology (traditional and PIC), separate regressions were calculated for the whole sample, for a subsample excluding Pinnipedia (i.e., a “fissiped” subsample, since pinnipeds showed atypical values for their body mass in most of the scatter plots), and for several subsamples by family and by locomotor type. Regressions were not calculated for any subsample with a sample size lower than 5, which was the case for Hyaenidae, Mephitidae, Phocidae, Otariidae, Prionodontidae, and the monotypic families (Ailuridae, Nandiniidae), and also for Eupleridae when using PIC regression.

Allometric exponents were considered to deviate significantly from the predictions of either similarity hypothesis when their 95%CI did not include the corresponding theoretical value. As stated in the introduction, according to GS, all linear dimensions should be proportional to **M**_**b**_^0.33^. Thus, all ratios, including relative lengths and bone robusticities, should present an allometric exponent not different from 0. On the other hand, ES proposes that lengths are proportional to **M**_**b**_^0.25^ and diameters to **M**_**b**_^0.375^, which derives into bone robusticities scaling with a theoretical exponent of 0.125 while ratios other than bone robusticities should present an allometric exponent not different from 0. Finally, angles, when measured in radians, can be considered lengths, and thus they should scale to **M**_**b**_^0.33^ or **M**_**b**_^0.25^, according to GS or ES, respectively.

For each variable, allometric exponents were then compared between the whole sample and the fissiped subsample, and between the different subsamples by family and locomotor type. Furthermore, the PIC slopes (*b*_*PIC*_) were compared to those obtained by traditional regression analysis (*b*_*trad*_) with an F-test (p < 0.05) to assess whether the phylogenetic signal had any effect on the results.

Finally, also for each variable and each subsample, the presence of differential scaling was also evaluated using the model proposed by Jolicoeur (1989):

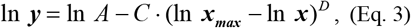

where *A* is a constant (corresponding to *a* in Eq. 1), *C* is the coefficient of allometry, **x**_**max**_ is the maximum observed value of the independent variable (i.e., body mass, **M**_**b**_), and *D* is the exponent of complex allometry, a time-scale factor. In our case, *D* > 1 indicated faster relative growth in small carnivorans, and *D* < 1 indicated that relative growth increased with size. The complex allometry hypothesis was thus accepted when *D* was significantly different from 1 (p < 0.05). Equation 3 was fitted with SPSS for Windows (release 15.0.1 2006; SPSS Inc., Chicago, IL, USA), and 95% confidence intervals were calculated for all parameters.

## Results

Supplementary Tables SR1 through SR30 show the regression results for each variable. As observed in previous studies comparing traditional and PIC regressions (Christiansen, 2002*a,b*; Gálvez-López & Casinos, 2012), the correlation coefficients (*R*) from the PIC analyses were lower than those from traditional regressions in most cases, which sometimes resulted in regressions no longer being significant (e.g. Table SR30). Some authors have attributed this phenomenon to a higher risk of type I errors (i.e., indicating a significant correlation between two variables when there was none) when the effect of phylogeny is neglected in correlation analyses (Grafen, 1989; Christiansen, 2002*a*). In some cases, however, *R* actually increased after taking into account the effect of phylogeny, which could result in regressions becoming significant (e.g. Table SR28). Branch lengths had to be transformed in most cases before performing the PIC regressions (Table S1).

### Whole sample vs. Fissiped subsample

No significant relation with body mass was found for the olecranon abduction angle (θ), or the robusticity of the ulna (**UR**) or the third metacarpal (**MR**). Neither was significant the regression of radial robusticity (**RR**) in the whole sample (*b*_*trad*_), nor those of **IFA** (*b*_*trad*_), **T** (*b*_*PIC*_), **%**_**prox**_ (*b*_*trad*_), and **%**_**dist**_ (both), after removing Pinnipedia (i.e., in the fissiped subsample).

Overall, removal of Pinnipedia from the sample caused a generalized increase of the allometric exponents when using traditional regression methods, although this increase was only significant for **L**_**h**_, **L**_**r**_, **L**_**u**_, and **%**_**mid**_. The exception to this general trend were **S, A, d**_**th**_, **HR**, and **d**_**tr**_, for which a reduction in the allometric exponent was observed (although it was only significant for **S**; Table SR2). These differences were not recovered by the PIC regressions, which produced fairly similar allometric exponents for the whole sample and the fissiped subsample. In fact, only for **d**_**tu**_ was the allometric exponent of the fissiped subsample significantly different from that obtained for the whole sample.

Contrary to previous studies comparing traditional and PIC regression methods (Christiansen, 2002*b*; Christiansen & Adolfssen, 2005; Gálvez-López & Casinos, 2012), significant differences between the allometric exponents obtained with each method were observed in the present study. In the case of **S, I, A**, and **d**_**tr**_ for both samples, and **L**_**s**_, **L**_**h**_, **d**_**th**_, **L**_**r**_, and **L**_**u**_ for the fissiped subsample, the allometric exponents obtained using traditional regression methods were significantly higher than PIC slopes (Tables SR1–SR4, SR6, SR8, SR11, SR13, SR16). On the other hand, in **HR** and α for both samples, **d**_**tm**_ for the whole sample, and **RR** and **d**_**tu**_ for the fissiped subsample, the PIC slopes were significantly higher than those obtained with traditional regression methods (Tables SR10, SR15, SR18, SR21, SR26).

Regarding conformity with the similarity hypotheses, Table 4 presents the percentage of linear measurements that conform to each similarity hypothesis in both the whole sample and the fissiped subsample, and also using either traditional regression methods or PIC. As indicated by the low percentages, the scaling pattern of the forelimb in Carnivora conformed poorly to either similarity hypothesis, no matter whether Pinnipedia was included in the sample. The decrease of most allometric exponents after taking into account phylogenetic relatedness resulted in about half the variables including 0.33 in their 95% CI_b_, improving thus conformity to the geometric similarity (see Table 4, PIC results). Again, results were the same with or without Pinnipedia.

**Table 4.** Conformity to the similarity hypotheses summary. For each subsample, the number of linear measurements conforming to geometric (G) or elastic similarity (E) is given, as is the percentage of the significant regressions for that subsample that they represent. Values in grey indicate that the number of variables conforming to a particular similarity hypothesis is either less than half the number of variables, or over 20% lower than the number of variables conforming to the other similarity hypothesis.

|  |  | traditional | PIC |  |  | traditional | PIC |
| --- | --- | --- | --- | --- | --- | --- | --- |
| whole sample | G | 7/19 (36.8%) | 9/19 (47.4%) | Viverridae | G | 16/19 (84.2%) | 17/17 (100%) |
|  | E | 4/19 (21.1%) | 5/19 (26.3%) |  | E | 15/19 (78.9%) | 14/17 (82.4%) |
| fissipeds | G | 2/19 (10.5%) | 9/19 (47.4%) | arboreal | G | 18/18 (100%) | 13/17 (76.5%) |
|  | E | 4/19 (21.1%) | 5/19 (26.3%) |  | E | 15/18 (83.3%) | 6/17 (35.3%) |
| Canidae | G | 11/19 (57.9%) | 10/19 (52.6%) | semiarboreal | G | 13/19 (68.4%) | 16/19 (84.2%) |
|  | E | 7/19 (36.8%) | 7/19 (36.8%) |  | E | 14/19 (73.7%) | 17/19 (89.5%) |
| Mustelidae | G | 14/19 (73.7%) | 17/19 (89.5%) | scansorial | G | 9/19 (47.4%) | 13/19 (68.4%) |
|  | E | 6/19 (31.6%) | 8/19 (42.1%) |  | E | 5/19 (26.3%) | 7/19 (36.8%) |
| Procyonidae | G | 18/18 (100%) | 7/7 (100%) | terrestrial | G | 5/19 (26.3%) | 18/19 (94.7%) |
|  | E | 17/18 (94.4%) | 7/7 (100%) |  | E | 6/19 (31.6%) | 7/19 (36.8%) |
| Ursidae | G | 8/18 (44.4%) | 6/8 (75.0%) | semifossorial | G | 18/18 (100%) | 18/18 (100%) |
|  | E | 14/18 (77.8%) | 8/8 (100%) |  | E | 17/18 (94.4%) | 17/18 (94.4%) |
| Felidae | G | 9/19 (47.4%) | 14/19 (73.7%) | semiaquatic | G | 18/19 (94.7%) | 17/17 (100%) |
|  | E | 9/19 (47.4%) | 7/19 (36.8%) |  | E | 14/19 (73.7%) | 13/17 (76.5%) |
| Herpestidae | G | 18/19 (94.7%) | 18/19 (94.7%) | aquatic | G | 12/17 (70.6%) | 6/11 (54.5%) |
|  | E | 9/19 (47.4%) | 11/19 (57.9%) |  | E | 11/17 (64.7%) | 7/11 (63.6%) |
| Eupleridae | G | 18/19 (94.7%) | – | freshwater | G | 18/19 (94.7%) | 18/19 (94.7%) |
|  | E | 16/19 (84.2%) | – |  | E | 14/19 (73.7%) | 14/19 (73.7%) |

Although **IFA** and the relative segment lengths were supposed to be independent of body mass according to both similarity hypotheses, this was not the case (Tables SR23, SR28–SR30). In the case of **T** a significant but minimal allometric effect was detected (Tables SR9). The olecranon angle (α) scaled with an exponent not significantly different from 0.33 in most cases (Tables SR21). Finally, regarding bone robusticities, regressions were only significant for **HR** and **RR**. Traditional regression provided conflicting results between the whole sample and the fissiped subsample in each bone robusticity. On the other hand, using PIC regression both bone robusticities in both subsamples scaled with positive allometry to body mass, no matter which similarity hypotheses was used (Tables SR10, SR15).

### Family subsamples

No significant differences were found between the allometric exponents obtained with each method (Tables SR1–SR30), which agrees with previous studies comparing traditional and PIC regression methods (Christiansen, 2002*b*; Christiansen & Adolfssen, 2005; Gálvez-López & Casinos, 2012).

Whereas the scaling pattern of some families conformed clearly better to the geometric similarity hypothesis (Mustelidae, Herpestidae) or the elastic similarity hypothesis (Ursidae), for others the 95% CI_b_ were wide enough to include the theoretic value for both hypotheses in most of the variables and no similarity hypothesis could be ruled out (Procyonidae, Eupleridae, Viverridae) (Table 4). In Canidae, conformity to the geometric similarity hypothesis was low (under 60%), but clearly better than to elastic similarity (under 40%, just diameters conformed to elastic similarity). In the case of Felidae, conformity to either similarity hypotheses was low when considering traditional regression results, since many of the narrow 95% CI_b_ excluded the theoretical values proposed by both hypotheses. Considering the PIC regression results, however, the felid scaling pattern clearly conformed to the geometric similarity hypothesis (Table 4).

As observed for the whole sample and the fissiped subsample, when significant, **IFA** scaled positively to body mass (except for Eupleridae; Table SR23), and **T** presented a significant but minimal allometric exponent (except for Mustelidae; Table SR9). In the case of relative segment lengths (Tables SR28–SR30), regressions were significant only in a few cases, but **%**_**prox**_ always increased with body mass (*b* > 0), while **%**_**mid**_ always decreased with increasing body mass (*b* < 0). Regarding the angles, regressions for θ were only significant for Herpestidae (*b*_*trad*_) and Canidae (*b*_*PIC*_), in both cases presenting allometric exponents very close to zero (Table SR20). On the other hand, the 95% CI_b_ for α included both 0.25 and 0.33 in all significant traditional regressions. However, after correcting for phylogeny, only the regression for Felidae remained significant (and scaled geometrically; Table SR21). Finally, regressions of bone robusticities on body mass were not significant in most cases, but when they were significant, their allometric exponents conformed better to the predictions of the hypothesis of elastic similarity, since they were in every case different from 0 (Tables SR10, SR15, SR22, SR27).

Figure 3 shows comparisons of the allometric exponents between different families for each variable, which are summarized in Table 5. No significant differences between families were found for **HR**, θ, α, **UR, IFA, MR, %**_**prox**_, **%**_**mid**_, or **%**_**dist**_. Overall, Canidae scaled faster than all other families in each case where significant differences between allometric exponents were found (especially when considering PIC regression results), while the relationships among the rest of the families varied among the variables studied.

**Figure 3.**
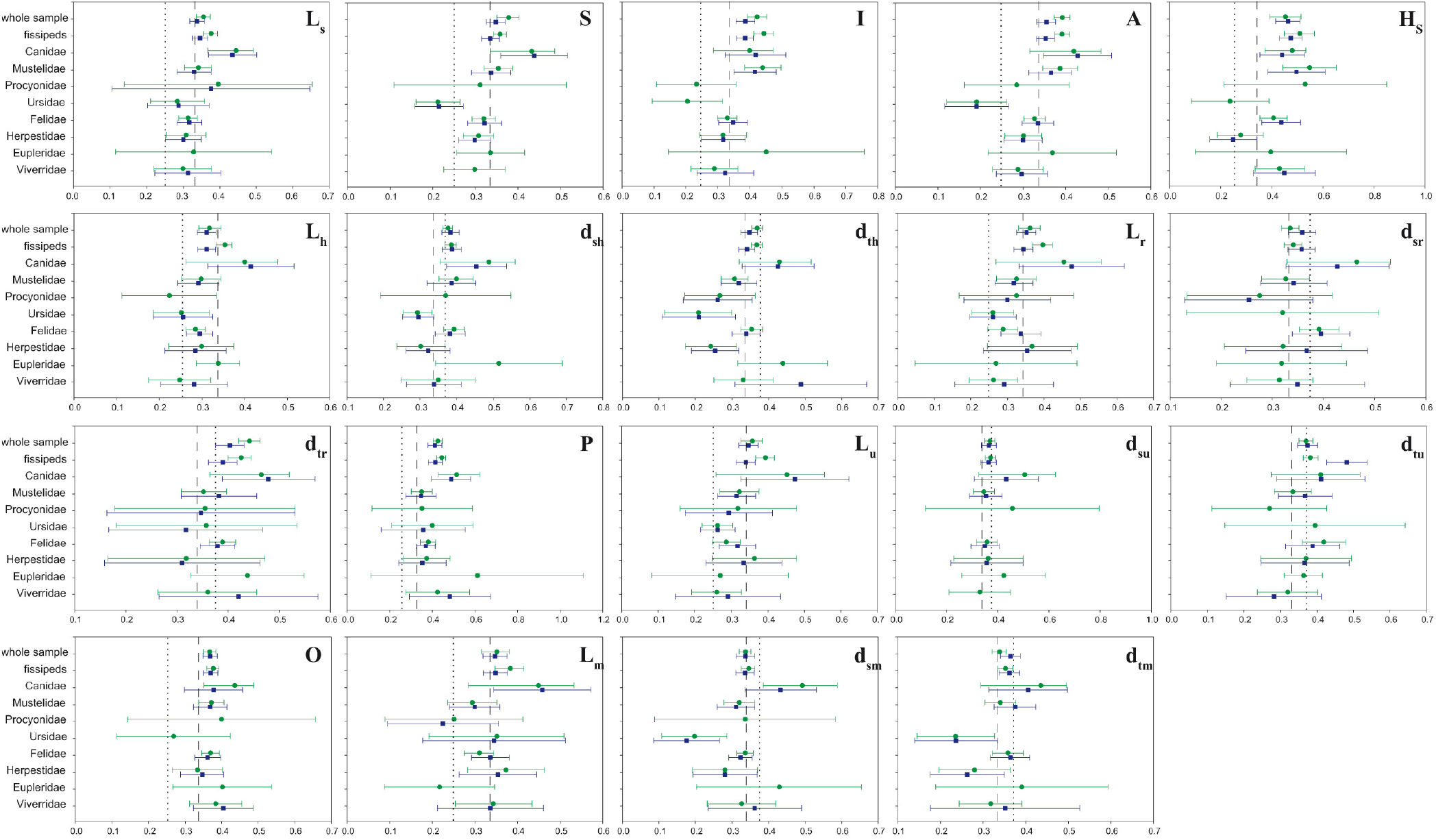
Allometric exponents by family. For each subsample, the allometric exponents obtained using traditional regression methods (green) and phylogenetically independent contrasts (blue), as well as their 95% confidence intervals, are shown. Only the results of significant regressions are presented. The allometric exponents obtained for the whole sample and the fissiped subsample are included as a reference. The dashed line represents the theoretical value proposed by the geometric similarity hypothesis, while the dotted line corresponds to that proposed by the elastic similarity hypothesis. Variable names are listed in Table 3.

**Table 5.**
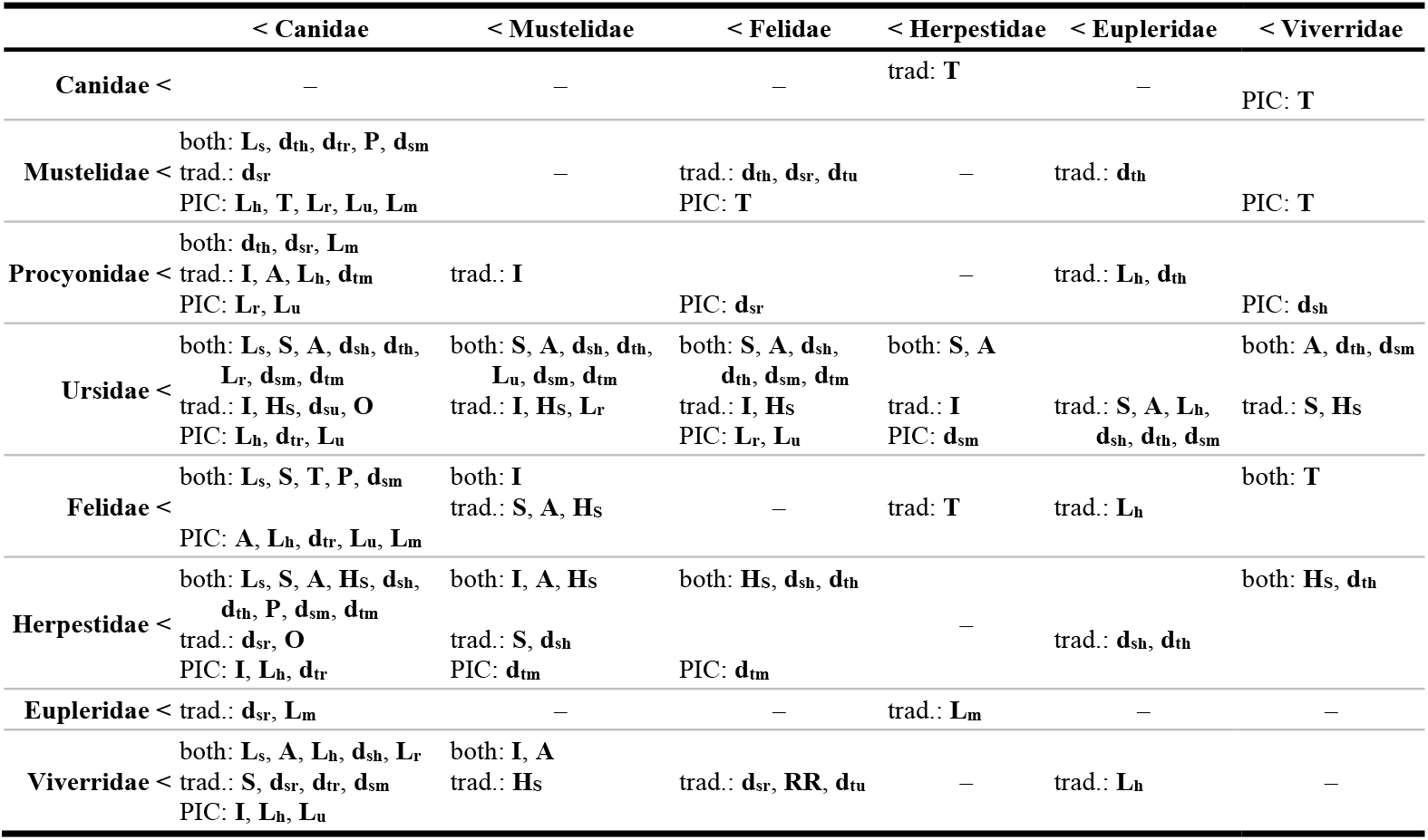
Differences in allometric exponents between families. Rows list families with an allometric exponent (*b*) significantly lower than the families listed in columns. That can happen when comparing allometric exponents from traditional regression (trad.), phylogenetically independent contrasts (PIC), or when using both methodologies (both). Variable names are listed in Table 3.

### Locomotor type subsamples

Contrary to previous studies comparing traditional and PIC regression methods (Christiansen, 2002*b*; Christiansen & Adolfssen, 2005; Gálvez-López & Casinos, 2012), significant differences between the allometric exponents obtained with each method were observed for some locomotor type categories. Most of these significant differences occurred in terrestrial carnivorans, where PIC slopes were generally lower than those obtained using traditional regression methods (**L**_**s**_, **I, A, L**_**h**_, **L**_**r**_, **d**_**tr**_, **P, L**_**u**_, **L**_**m**_; Tables SR1, SR3, SR4, SR6, SR11, SR13, SR14, SR16, SR24, respectively). However, significantly lower PIC slopes were also found for scansorial carnivorans (**d**_**th**_; Table SR8). Finally, PIC slopes were significantly higher for **%**_**mid**_ in terrestrial carnivorans (Table SR29).

The scaling pattern of scansorial and semiaquatic carnivorans conformed better to the geometric similarity hypothesis (Table 4). In the case of semiarboreal, semifossorial, and aquatic carnivorans, however, the 95% CI_b_ were wide enough to include the theoretic value for both hypotheses in most of the variables and thus no similarity hypothesis could be ruled out. In the case of arboreal carnivorans, conformity to both similarity hypotheses was high when considering traditional regression results. On the other hand, for terrestrial carnivorans, the scaling pattern obtained using traditional regression methods did not conform to any similarity hypothesis. Considering the PIC regression results, however, the scaling pattern of both locomotor types clearly conformed to the geometric similarity hypothesis (Table 4).

Regarding ratios and angles, the results were similar to those obtained for the whole sample, the fissiped subsamples and the family subsamples. First, when significant, **IFA** scaled positively to body mass (except for arboreal and terrestrial carnivorans, *b*_*PIC*_ and *b*_*trad*_ respectively; Table SR23), and **T** presented a significant but minimal allometric exponent (Table SR9). And second, in the case of relative segment lengths (Tables SR28–SR30), **%**_**prox**_ always increased with body mass (*b* > 0), while **%**_**mid**_ generally decreased with increasing body mass (*b* < 0; except for arboreal and semiaquatic carnivorans, *b*_*PIC*_ both). On the other hand, **%**_**dist**_ either increased (terrestrial, aquatic) or decreased (semiarboreal, semiaquatic) with body mass. Regarding the angles, again regressions for θ were only significant in two cases, in both cases presenting allometric exponents very close to zero: semifossorial (*b*_*trad*_, *b* > 0) and arboreal (*b*_*PIC*_, *b* < 0) (Table SR20). The scaling of the olecranon angle (α) conformed either to elastic similarity (scansorial, *b*_*trad*_), to geometric similarity (scansorial, *b*_*PIC*_), or to both (terrestrial, *b*_*trad*_) (Table SR21). Finally, although the allometric exponents for bone robusticities were positive and conforming to the elastic similarity hypothesis for most locomotor types (Tables SR10, SR22, SR27), contrary to the results for the previous subsamples, the allometric exponents were negative in the radius, ulna, and third metacarpal, of terrestrial carnivorans (*b*_*trad*_ in all cases), indicating that bone robusticity decreased with increasing body mass values (Tables SR15, SR22, SR27).

Figure 4 shows comparisons of the allometric exponents between different locomotor types for each variable, which are summarized in Table 6.

**Figure 4.**
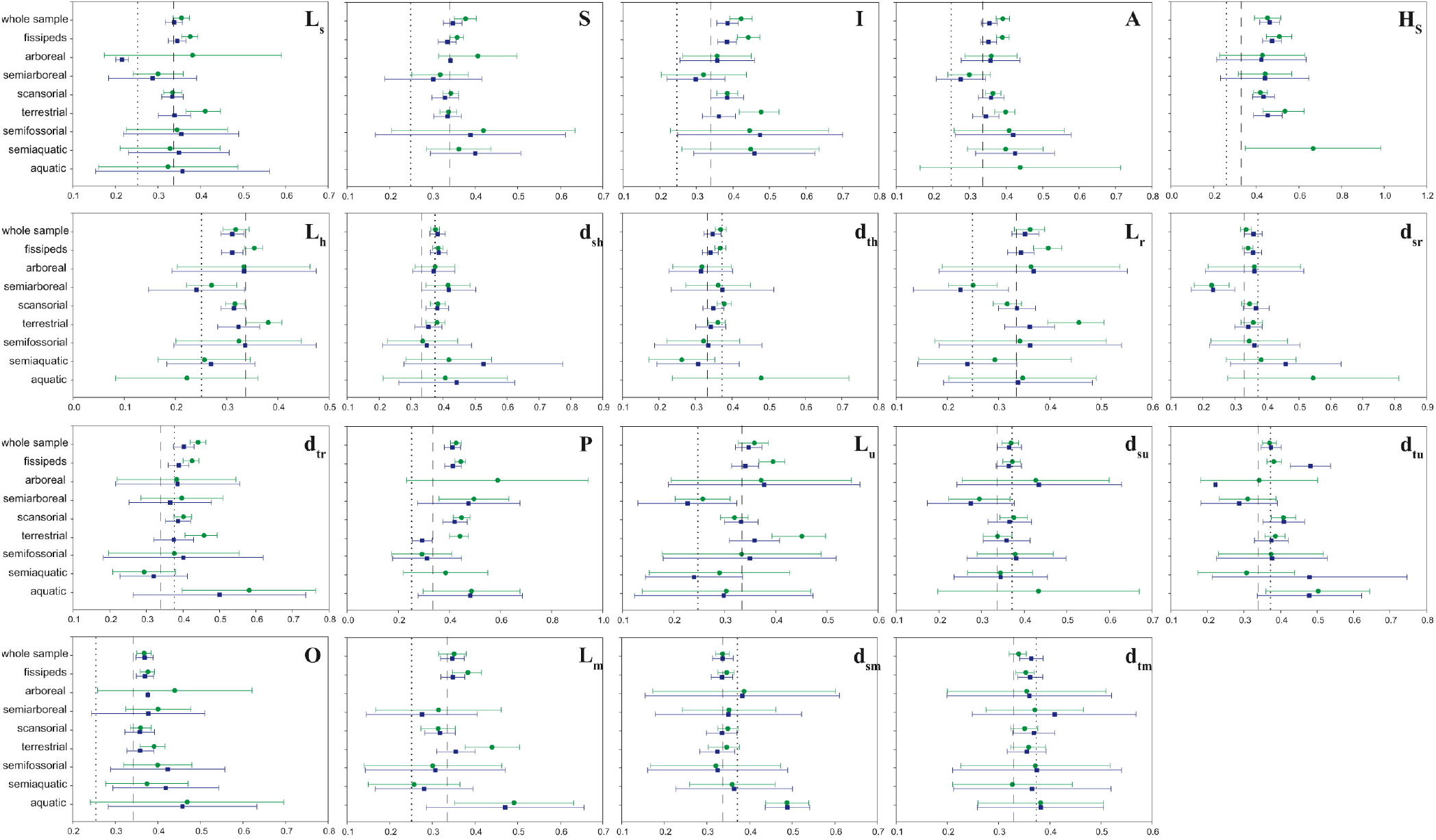
Allometric exponents by locomotor type. For each subsample, the allometric exponents obtained using traditional regression methods (green) and phylogenetically independent contrasts (blue), as well as their 95% confidence intervals, are shown. Only the results of significant regressions are presented. The allometric exponents obtained for the whole sample and the fissiped subsample are included as a reference. The dashed line represents the theoretical value proposed by the geometric similarity hypothesis, while the dotted line corresponds to that proposed by the elastic similarity hypothesis. See Table 2 for a description of locomotor type categories. Variable names are listed in Table 3.

**Table 6.**
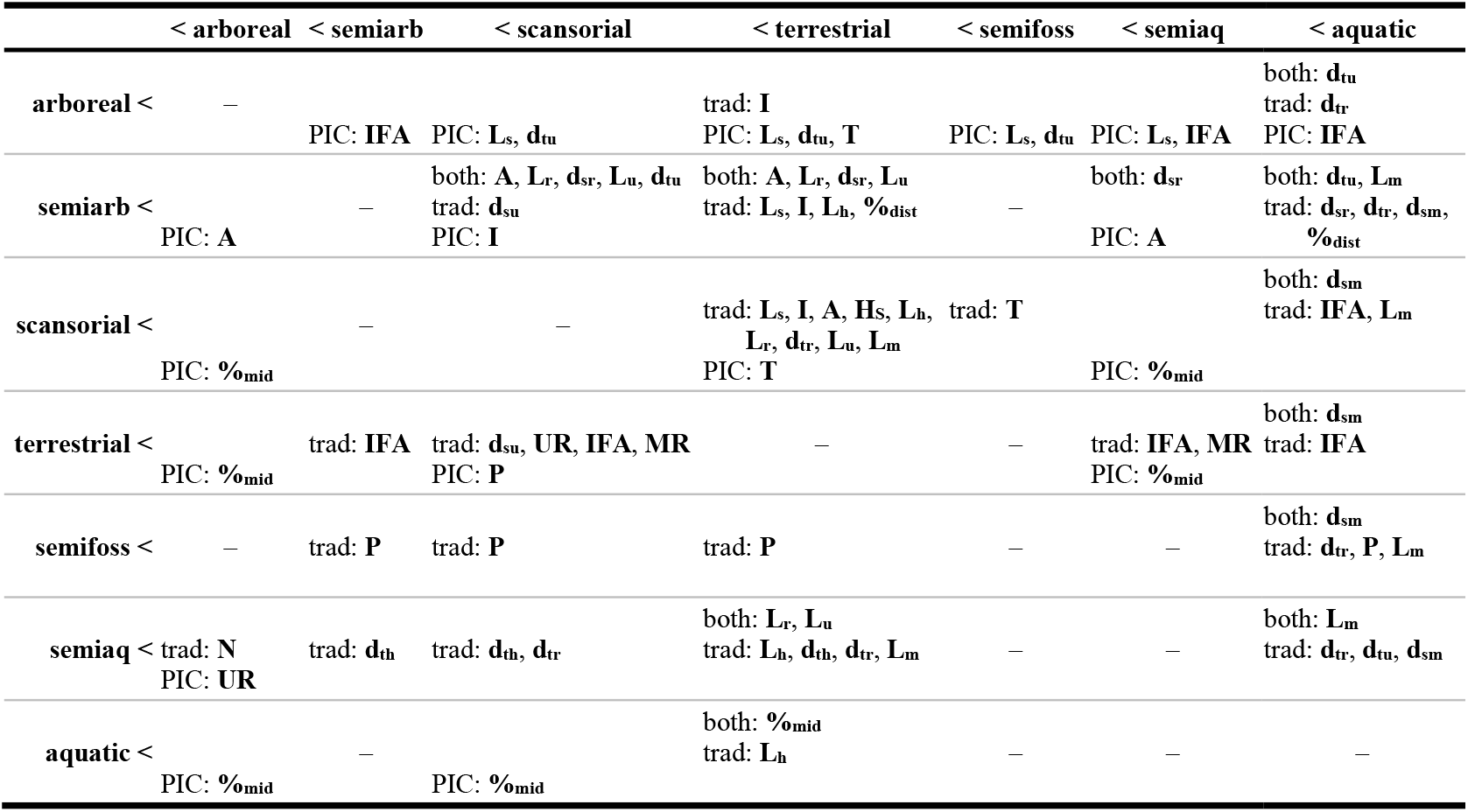
Differences in allometric exponents between locomotor types. Rows list categories with an allometric exponent (*b*) significantly lower than the categories listed in columns. That can happen when comparing allometric exponents from traditional regression (trad.), phylogenetically independent contrasts (PIC), or when using both methodologies (both). Variable names are listed in Table 3.

### Complex allometry

Results for the test for complex allometry are shown in Tables SR31 through SR59. Since **T** presented negative values, Equation 3 could not be fit, which made impossible testing for complex allometry with this method.

In the whole sample, evidence for complex allometry was found in almost half of the variables. In the case of **L**_**s**_, **I, H**_**S**_, **L**_**h**_, **L**_**r**_, **L**_**u**_, **O, L**_**m**_, and **%**_**dist**_, *D* was significantly higher than 1, indicating that these variables scale faster in small species; while in **HR, IFA, %**_**prox**_, and **%**_**mid**_, *D* was significantly lower than 1, suggesting that these variables scale faster in large species. However, in all cases where *D*<1, the 95% CI_*D*_ included 0, which would result in (ln ***x***_***max***_ − ln ***x***)^*D*^ = 1, and hence ln ***y*** = ln *A* − *C*, which indicates independence from the dependent variable ***x*** (here body mass).

After removing Pinnipedia from the sample (i.e. in the fissiped subsample), evidence for complex allometry was not recovered in most cases. Only for **H**_**S**_, **O, L**_**m**_, and **%**_**dist**_, was *D* still significantly different from 1 (*D* > 1 in all cases). Furthermore, significant evidence for complex allometry was also found for **d**_**tu**_, which presented *D* < 1.

Overall, significant evidence for complex allometry was scarce in the family subsamples. In Procyonidae, Ursidae and Felidae no variable presented complex allometry, while in Canidae and Eupleridae only one variable presented complex allometry in each subsample (respectively, **P** and **d**_**sh**_; *D*>1 in both cases). On the other hand, some variables presented significant evidence for complex allometry in Mustelidae (**HR, d**_**tr**_, **%**_**prox**_, **%**_**mid**_), Herpestidae (**L**_**s**_, **H**_**S**_, **P**), and Viverridae (**I, L**_**h**_, **L**_**r**_, **d**_**sr**_, **d**_**tr**_, **L**_**u**_), with *D*<1 in all cases. However, as observed for the whole sample when *D*<1, in some cases the 95% CI_*D*_ also included 0, indicating independence from body mass. This was the case for **HR, %**_**prox**_ and **%**_**mid**_ in Mustelidae, **H**_**S**_ and **P** in Herpestidae, and **L**_**h**_ in Viverridae.

Finally, in the locomotor type subsamples, significant evidence for complex allometry was even less frequent than in the family subsamples. Thus, evidence for complex allometry was only found for **A, L**_**h**_, **d**_**tr**_ and **L**_**u**_ in semiarboreal carnivorans, for **H**_**S**_, **L**_**h**_, **P, O, L**_**m**_ and **d**_**sm**_ in scansorial carnivorans, and for **d**_**su**_ in terrestrial carnivorans. In terrestrial and semiarboreal carnivorans, when complex allometry was detected, it indicated that large carnivorans scaled faster than small species (i.e. *D* < 1), while the opposite was true for scansorial carnivorans (i.e. *D* > 1). No 95% CI_*D*_ included 0.

## Discussion

### Considerations on the scaling pattern of the carnivoran forelimb

The present study is currently the largest and most thorough work on skeletal allometry in Carnivora, regarding both the number of species sampled and the skeletal elements considered. In fact, even when considering all previous allometric studies on Mammalia, only that of Christiansen (1999*a*) on long-bone allometry and that of Silva (1998) on the scaling of body length include a larger amount of species.

Regarding the scaling of the appendicular skeleton in Carnivora, similarly to previous studies on the subject in this and other groups (Bou et al., 1987; Bertram & Biewener, 1990; Christiansen, 1999*a,b*; Carrano, 2001; Llorens et al., 2001; Lilje et al., 2003; Casinos et al., 2012), conformity to either the geometric similarity hypothesis or the elastic similarity hypothesis was low. It could be argued that geometric similarity provided a better explanation than elastic similarity (Table 4), but that was only because no length scaled elastically. Also in agreement with previous studies (Economos, 1983; Bertram & Biewener, 1990; Silva, 1998; Christiansen, 1999*a,b*; Carrano, 2001), significant evidence for complex allometry was found in several of the studied variables. Finally, contrary to previous studies comparing traditional regression methods and phylogenetically independent contrasts (PIC) (Christiansen, 2002*b*; Christiansen & Adolfssen, 2005; Gálvez-López & Casinos, 2012), significant differences between the allometric exponents of both methodologies were found, especially in the fissiped subsample. Thus, in order to avoid any possible artefacts caused by the phylogenetic relatedness of the species in our sample, only the PIC results will be further discussed.

One of the predicted consequences of increasing size is enduring higher peak stresses (especially during locomotion), which could lead to mechanical failure (Alexander, 2002). Thus, as mammals get larger, they must either develop more robust bones to resist these higher stresses or change their limb posture to reduce the magnitude of these stresses (Biewener, 2003; Carrano, 2001). Based on previous results, it has been proposed that limb posture changes might be the preferred strategy to cope with the size-related increase of peak stresses, but that at body masses over 200kg more robust bones must be developed, since limbs cannot be further straightened (Christiansen, 1999*a*; Carrano, 2001). The change in bone scaling required to develop more robust bones in large mammals has commonly been considered the cause of differential scaling in bone dimensions (Biewener, 1990; Christiansen, 1999*b*). In Carnivora, only a handful of non-aquatic species attain such large body sizes, suggesting that peak stresses should be reduced in this group by limb straightening, not by changing limb bone scaling. In the present study, two arguments were found against this assumption. First, significant evidence for differential scaling was found in several variables, indicating that the scaling of the forelimb does change with size in Carnivora. However, since the amount of variables showing complex allometry severely decreased after removing Pinnipedia, most of these scaling changes are probably related to their specialized biology and ecomorphology and not to reducing peak stresses. Second, limb bones seemed to scale elastically in Ursidae, which includes most of the largest non-aquatic carnivorans. Since the elastic scaling of limb bones in Bovidae (which includes most of the largest non-aquatic mammals) was one of the main arguments supporting that large mammals develop more robust bones to cope with increased peak stresses (Economos, 1983; Christiansen, 1999*a*), the present results for Ursidae would point to a similar conclusion. However, the elastic scaling of Ursidae could be an artefact caused by the combination of their overall lower allometric exponents than other families (i.e., both in lengths and diameters) and their wide 95%CI_b_ (Fig. 3). Furthermore, although the regressions for bone robusticities are not significant in Ursidae, their allometric exponents are not higher than those of other carnivoran families. In fact, they were lower than in most other families, especially for the humerus (**HR**; Table SR10). Thus, the results of the present study support that, in large non-aquatic carnivorans, mechanical failure is mainly avoided by limb posture changes instead of by modifying limb bone scaling. Further evidences for this conclusion are the lack of differential scaling in the “large” families (Canidae, Felidae, Ursidae; Tables SR31–SR59) and the significant increase with size of the olecranon angle (α; Table SR21), especially in the fissiped subsample. This angle determines the position in which the triceps muscle has the greatest leverage, being a flexed elbow when α is small (straight or cranially bent olecranon) or an extended limb when it is large (caudally bent olecranon) (Van Valkenburgh, 1987). Thus, an allometric increase of α suggests that large carnivorans have increasingly straighter forelimbs (but see Day & Jayne, 2007).

Several authors have suggested that proximal limb segments are more conservative in lengthening with increasing body mass than distal ones (McMahon, 1975*a*; Lilje et al., 2003; Schmidt & Fischer, 2009). According to this, when regressing bone length to body mass, proximal bones should produce higher correlation coefficients, and, when comparing allometric exponents, significant differences between subsamples should be scarce for proximal segments. While this might be the case for Artiodactyla (McMahon, 1975*a*; Lilje et al., 2003), the results of the present study suggest that, while it might also apply for Carnivora as a whole, the more conservative nature of proximal limb segments is not evident in several carnivoran subsamples. For instance, the highest correlation coefficients correspond to the radius and ulna in Procyonidae and Ursidae, and to the third metacarpal in aquatic carnivorans. Furthermore, when comparing the allometric exponents obtained for bone lengths, significant differences were found for all forelimb bones in all subsample sets (i.e., by family and by locomotor type).

Previous studies had reported differences in the scaling of the various forelimb bones (Wayne, 1986; Bertram & Biewener, 1990; Christiansen, 1999*a*; Lilje et al., 2003). In those studies, the lengths of the middle segment (i.e., humerus) tended to scale slower than the rest of the forelimb segments. The scaling of the proximal element (i.e., scapula) was seldom described, but it presented intermediate values between the humerus and the distal elements in Canidae (Wayne, 1986) and the fastest scaling in Ruminantia (Lilje et al., 2003). In the present study the humerus presented the lowest allometric exponent in almost all subsamples, but no significant differences were found among the other forelimb bones. Only in arboreal, semiarboreal and semifossorial carnivorans the humerus scaled faster than other segments consistently (scapula, radius/ulna and third metacarpal, respectively). Together with previous results, this suggests that the slow scaling of the humerus relative to the other forelimb segments could be a common trend in Mammalia, with groups with particular locomotor adaptations (such as climbing or digging) deviating from this pattern.

Furthermore, the slow scaling of the humerus relative to other forelimb segments would explain the negative allometry found for its relative length (**%**_**mid**_) both here and in the study of Schmidt & Fischer (2009). Regarding bone diameters, few studies have obtained confidence intervals narrow enough to describe differences in the scaling of different bones: Cubo & Casinos (1998) reported a faster scaling of the transverse diameter of the radius (**d**_**tr**_) relative to the sagittal diameter of the radius and both humerus diameters in Mammalia. On the other hand, while comparing the same bones, Heinrich & Biknevicius (1998) and Llorens et al. (2001) found higher allometric exponents for the sagittal diameter of the humerus (**d**_**sh**_) than for other bone diameters in Martinae and Platyrrhina, respectively. The results of the present study in Carnivora showed that the sagittal diameter of the third metacarpal (**d**_**sm**_) scaled significantly slower than most other bone diameters, and the transverse diameters of both radius and ulna (**d**_**tr**_, **d**_**tu**_) and the sagittal diameter of the humerus (**d**_**sh**_) scaled significantly faster than most other bone diameters. In the case of **d**_**sh**_, our results suggest that the conflicting results found in previous studies could be related to whether the deltoid tuberosity was included in its measurement, since it was included within **d**_**sh**_ in the present study, and only in Viverridae, whose species do not present a particularly developed deltoid tuberosity, scaled **d**_**sh**_ significantly slower than **d**_**th**_ (Tables SR7, SR8). Finally, regarding the fast scaling of **d**_**tr**_ and **d**_**tu**_, it could be related to a greater development of the muscles originating in the shaft on the radius and ulna (pronators and supinators of the hand, some wrist flexors and extensors). These increased forearm muscles would provide a stronger grip to large climbing species (e.g. bears) and also to species relying in the forelimb for prey capture (e.g. felids), but would also cause larger mediolateral stresses on those bones, hence the need of increased transverse diameters. In agreement to this, significant evidence for differential scaling was found for **d**_**tr**_ in fissipeds and in semiarboreal carnivorans, in both cases with larger species scaling faster than small species. Aiello (1981) stated that the use of ratios is only correct when both variables comprising it scale isometrically between them. In agreement with this, due to differences in scaling among bone lengths, the allometric exponents found for the relative length of the proximal and middle segment (**%**_**prox**_, **%**_**mid**_; Tables SR28, SR29) and the indicator of fossorial ability (**IFA**; Table SR23) were significantly different from zero, the value predicted by both similarity hypotheses. Furthermore, the present results on the scaling of relative segment lengths of the forelimb in Carnivora mirrored those obtained previously for Schmidt & Fischer (2009) in both Carnivora and Artiodactyla: relative humerus length scales negatively to body mass, while the relative scapula length does it positively. Finally, it has been proposed that group-specific differences in limb kinematics are characteristic of large mammals, since small mammals are relatively similar in limb kinematics regardless of locomotor habit and phylogenetic position (Fischer et al., 2002; Schmidt & Fischer, 2009). Furthermore, small mammals present crouched limbs and large mammals extended limbs, each requiring different sets of limb-segment proportions for self-stability (Seyfarth et al., 2001). Thus, since Carnivora includes both small and large species, differential scaling would be expected for their relative segment lengths, as it has been found in the present study (Tables SR57–SR59).

### Phylogenetic deviations to the scaling of the carnivoran forelimb

Overall, the scaling patterns found in the different carnivoran families for the forelimb were similar to the pattern found in the whole order. However, several families deviated significantly from it (Fig. 3). In the case of Canidae, scapula and humerus length (**L**_**s**_, **L**_**h**_), as well as the maximum width of the supraspinous fossa (**S**), scaled faster than in the rest of Carnivora. Furthermore, when comparing the allometric exponents obtained for each variable between families, Canidae scaled faster than all other families in each case. This agrees with the expectations of Wayne (1986), who suggested that size selection is likely one of the most predominant forces in canid evolution because size differences help mitigate interspecific competition. On the other hand, several variables scaled significantly slower in Ursidae and in Herpertidae than in the whole sample (Fig. 3). Finally, it should be noted that the wide confidence intervals (95%CI_b_) obtained for some families could be obscuring further significant deviations from the ordinal scaling pattern (e.g. Procyonidae, Eupleridae, Viverridae).

The lack of significant differences between the allometric exponents calculated using traditional and PIC regression methods agrees with a previous study stating that most morphological variability of the appendicular skeleton in Carnivora occurs at the family level (Gálvez-López, 2021).

Regarding conformity to the similarity hypotheses, the present results agree with those of Bertram & Biewener (1990) in that 1) Ursidae tended to conform better to the elastic similarity hypothesis; 2) mustelids scaled geometrically; and 3) conformity to either similarity hypotheses was low in Canidae, but slightly better to geometric similarity. However, contrary to the results of Bertram & Biewener (1990) but in agreement with those of Day & Jayne (2007) and Gálvez-López & Casinos (2012), felids conformed well to the geometric similarity hypothesis. The wide 95%CI_b_ obtained for Procyonidae in both studies made both similarity hypotheses equally (un)likely. According to Wayne (1986), bone diameters in Canidae were expected not to conform to the elastic similarity hypothesis, which was not the case in the present study. The conflicting results of the present study could be caused by the lower sample size (17 spp here vs. 27 in Wayne, 1986) or, more probably, by the different independent variables used, since in the present study all variables were regressed to body mass, and Wayne used femur length.

Finally, an interesting pattern was found among the families of Caniformia: for most linear measurements, the allometric exponents consistently increased from Ursidae to Procyonidae, to Mustelidae, and then to Canidae (Fig. 3). Neither body mass nor phylogenetic relatedness could explain this pattern, since Canidae and Ursidae represent both the largest caniforms, and the first phyletic lines to diverge from the caniform stem, and are placed in opposite extremes of this pattern. A possible explanation to this pattern could be an increasing degree of adaptation to overground locomotion, or a decrease in arboreal activity. Of all bears studied, only the polar bear (*Ursus maritimus*) is not an adept climber, since young brown bears (*Ursus arctos*) do climb (Gambaryan, 1974; Wilson & Mittermeier, 2009). Procyonids stand in a similar position, which could explain why they present lower allometric exponents than bears for some variables. Several mustelid lineages have diverged from the scansorial lifestyle (e.g. Lutrinae, Mustelinae), and thus Mustelidae presents intermediate values between ursids/procyonids and Canidae, which are fully adapted to a completely terrestrial lifestyle (understanding here the word “terrestrial” as defined in Table 2, i.e., with no specific climbing, digging, or swimming capabilities). In agreement with this, the four studied families within Feliformia, all of which but Herpestidae included species with a varied degree of climbing skills, presented similar allometric exponents in most variables (Fig. 3). In fact, only the terrestrial Herpestidae presented, in a few cases, allometric exponents significantly different from the rest of feliform families (Table 5). Another possible explanation could be a different degree of size selection within each caniform family. Both the present study and that of Wayne (1986) suggest size selection as a major force in canid evolution. However, nothing is known on the importance of size selection in the rest of caniform families.

### Locomotor habit and the scaling pattern of the carnivoran forelimb

Lilje et al. (2003) suggested that the scaling of limb bone lengths is more heavily influenced by phylogenetic relatedness than by habitat preference, at least in Artiodactyla. The present results suggest that this might also be the case for Carnivora, since the comparison of allometric exponents for bone lengths obtained using traditional regression methods produced more significant differences than the comparison of PIC slopes for the same variables among locomotor types.

Regarding the particular deviations associated to each locomotor type, in arboreal carnivorans scapular length (**L**_**s**_) and ulna transverse diameter (**d**_**tu**_) increased with body mass with significantly lower exponents than those obtained for Carnivora as a whole and the fissiped subsample (Fig. 4). However, the narrow 95%CI_b_ and high *R* for these regressions were unexpected given the low sample size of the arboreal subsample, suggesting that these results should be regarded cautiously (Tables SR1, SR18). Thus, the deviations observed for semiarboreal carnivorans probably represent a more accurate description of the scaling pattern associated to species spending most of their time in the canopy. In this subsample, significantly lower allometric exponents than those obtained for Carnivora were obtained for the functional length of the radius and the ulna (**L**_**r**_, **L**_**u**_), the sagittal diameter of the radius (**d**_**sr**_), and most scapular widths (**A, I**) (Fig. 4; Table 6). Similar deviations were found for the other functional bone lengths (**L**_**s**_, **L**_**h**_, **L**_**m**_) and the width of the supraspinous fossa (**S**), although they were not significant (Fig. 4). Furthermore, in all these cases, the allometric exponents for semiarboreal carnivorans were lower than those for scansorial and terrestrial species (Fig. 4), often significantly (Table 6). Thus, with increasing size, semiarboreal carnivorans will present shorter limbs and narrower scapulae than similar-sized scansorial and terrestrial species. According to Cartmill (1985), the first would be a strategy to increase stability during arboreal locomotion for claw-climbing mammals, like carnivorans, since relatively shorter limbs enable to maintain their center of mass close to the support, and thus reduce lateral oscillations of the center of mass. Carnivorans less adapted to moving in arboreal settings, such as scansorial species, should then resort to postural changes and other strategies in order to gain in stability when navigating arboreal supports, as demonstrated for the domestic cat by Gálvez-López et al. (2011). Continuing with adaptations to arboreality, in a study on forelimb morphology in North American carnivorans, Iwaniuk et al. (1999) found that the degree of arboreality was positively correlated with long-bone robusticities (calculated as **L**_**x**_/**d**_**sx**_). Thus, they stated that, with increasing arboreality, forelimb bones became wider, more robust, to better withstand the multidimensional loads resulting from arboreal locomotion. However, from the definition of their ratios, their results seemed to indicate just the opposite, that is, that arboreal carnivorans presented less robust forelimb bones (i.e., relatively longer or more slender bones). In the present study, the regressions of bone robusticities onto body mass tended to produce higher allometric exponents in the subsamples with the most arboreal species (e.g. **HR**: allometric exponents for semiarboreal carnivorans were higher than for scansorial and terrestrial carnivorans; Table SR10). Since in the present study bone robusticity was the inverse of the definition of Iwaniuk et al. (1999) (i.e., **d**_**sx**_/**L**_**x**_), these higher allometric exponents did indeed suggest that forelimb bones become sturdier (i.e., relatively wider or shorter) with increasing arboreality in Carnivora. Finally, regarding the pattern of increasing allometric exponents with decreasing arboreality found in Caniformia, it was not recovered in most cases in the locomotor type subsamples (Fig. 4), which could be explained by feliform species making up around 70% of the arboreal, semiarboreal and scansorial subsamples.

Although all mammals run (i.e., present gaits, either symmetrical or asymmetrical, in which their limbs spend less than half a cycle on the ground; Alexander, 2002; Biewener, 2003), some of them have developed certain morphological adaptations to increase step length (and thus speed) and to minimize energy costs while running (e.g. Gambaryan, 1974; Hildebrand, 1985). These mammals better adapted to running are often referred to as “cursorial mammals” (Smith & Savage, 56; Gambaryan, 1974; Hildebrand, 1985). However, as pointed out by Stein & Casinos (1997), the works of Jenkins and other authors (Jenkins, 1971; Jenkins & Camazine, 1977; Alexander & Jayes, 1983) introduced ambiguity into the concept of “cursorial” so it no longer meant “specialized runner”. Thus, the term “cursorial” will not be used in the present work, and instead “efficient runner” will be used to designate those mammals that have developed morphological adaptations to run efficiently. It has been described that presenting long limbs is an adaptation to effective running, since it allows for longer steps and thus higher speeds (Lull, 1904; Gambaryan, 1974; Hildebrand, 1985; Van Valkenburgh, 1987). However, limb elongation is mainly effected through the distal segments (Hildebrand, 1985; Van Valkenburgh, 1987), and thus, the radius, ulna and metacarpals of running species should scale faster than the humerus. In the present study, there was not a specific subsample grouping “efficient runners”, but two subsamples included a fair amount of those species: Canidae, and terrestrial carnivorans. Thus, bone lengths were expected to scale faster in these subsamples than in other subsamples. Additionally, **L**_**r**_, **L**_**u**_ and **L**_**m**_ were expected to scale faster than **L**_**h**_. Both assumptions were supported by the results of the present study (Figs. 3–4; Tables 5–6). Another adaptation to effective running was proposed by Smith & Savage (1956), who described larger infraspinous fossae than supraspinous fossae in mammals adapted to running. Thus, it was expected that **I** scaled faster than **S** in Canidae and terrestrial carnivorans. However, the present results suggest that a faster scaling of the infraspinous fossa is a common trend in Carnivora, not a particular adaptation to running efficiently. Oddly enough, Canidae was one of the subsamples deviating from this general trend. Thus, it might be concluded that previously described adaptations to effective running other that limb elongation are present in the scaling of most carnivoran subsamples (not just “effective runners”), which suggests that they are more related to the biomechanical consequences of increasing size than to effective running.

The effect of adaptations to digging and swimming to the scaling pattern of the carnivoran forelimb were hard to ascertain, since 95%CI_b_ were usually too wide in semifossorial, semiaquatic and aquatic carnivorans. In the case of semifossorial carnivorans, they presented high allometric exponents for scapular widths (**S, I, A**) and olecranon length (**O**), but they were not significantly different from any other subsample due to high 95%CI_b_ (Fig. 4; Table 6). Regarding adaptations to swimming, both semiaquatic and aquatic carnivorans tended to present high allometric exponents for scapular widths (**S, I, A**), olecranon length (both absolute, **O**, and relative, **IFA**), and several bone diameters (**d**_**sh**_, **d**_**sr**_, **d**_**tu**_) and bone robusticies (**HR, RR, UR**) (Fig. 4; Table 6). Furthermore, in semiaquatic carnivorans bone lengths scaled slower than in most carnivorans (significantly in the middle segment: **L**_**r**_, **L**_**u**_), while in aquatic carnivorans the third metarcapal scaled faster than in the rest of Carnivora, in both sagittal diameter and length (Fig. 4; Table 6). Most of these adaptations had already been suggested by previous anatomical and morphometrical analyses (Osburn, 1903; Smith & Savage, 1956; English, 1977; Gálvez-López, 2021), and were recovered here as characteristic deviations of the aquatic/semiaquatic scaling pattern: shorter and more robust limb bones, larger olecrana (both **O** and **IFA**), and wider scapulas (although not in semiaquatic carnivorans).

### Differential scaling, phylogeny and locomotor habit

According to Bertram & Biewener (1990), differential scaling might not be evident within the individual carnivoran families due to their narrow body size ranges. Furthermore, they also stated that differences in scaling explained by differences in locomotor habit would probably be overridden by phylogenetic differences in scaling. Those concerns proved irrelevant in the present study, since not only did more significant cases of complex allometry were found in Viverridae (**M**_**b**_ range: 0.54kg – 13.25kg) than in other families with wider body mass ranges (Canidae, Felidae, Mustelidae), but also significant cases of complex allometry were detected in several locomotor type subsamples (again, regardless of body mass range).

Previous studies have suggested that differential scaling could be a consequence of mixing species with different locomotor specializations (Castiella & Casinos, 1990; Gálvez-López & Casinos, 2012). The results of the present study provide arguments both in favour and against this hypothesis. On one hand, significant evidence for complex allometry was found in almost half the variables in the whole sample. Furthermore, several variables presented differential scaling in Mustelidae and Viverridae, both including species with several locomotor types, and the latter also presenting a narrow body mass range. On the other hand, after removing the large, swimming, pinniped species, significant evidence for complex allometry was rarely found. Furthermore, differential scaling was found in some locomotor type categories.

### On the viability of similarity hypotheses and scaling studies

The present and previous results on the scaling of limb bone morphology have made clear that no similarity hypothesis alone can explain the scaling patterns existing in mammalian limb bones (Bou et al., 1987; Bertram & Biewener, 1990; Christiansen, 1999*a,b*; Carrano, 2001; Llorens et al., 2001; Lilje et al., 2003; Casinos et al., 2012). In our understanding, the main problem with any similarity hypothesis is their extremely simplistic approach: each similarity hypothesis chooses one of the many factors determining how limb bone morphology changes with increasing size and defines allometric exponents based on it (geometric similarity: isometric growth; elastic similarity: deformation under gravity; static stress: constant stresses while standing still; dynamic stress: constant stresses during locomotion; McMahon, 1973, 1975*b*; Alexander & Jayes, 1983; Alexander, 2002). Thus, since no such single determining factor exists, all similarity hypotheses are doomed to fail. However, their inability to produce an accurate theoretical allometric exponent is instead excused by stating that variability around that “universal” trend is clouding the results, and thus the observed allometric exponents deviate from the predicted ones.

A further problem is that large and small mammals have different locomotor requirements (Lilje & Fischer, 2001; Seyfarth et al., 2001; Fischer et al., 2002; Schmidt & Fischer, 2009). This results in differential scaling and its oversimplification by establishing a threshold body mass value with which separate those small and large mammals, and thus be able to ascribe them separately to some similarity hypothesis (or a similarity hypothesis with different allometric exponents for small and large mammals; Garcia & da Silva, 2006). But see also Kokshenev (2003, 2007) for a criticism of Garcia-Silva’s model. The thing with differential scaling is that it is indeed differential. As observed in any plot representing complex allometry (Fig. 5), the allometric exponent changes gradually along a wide spectrum of body masses, and no real threshold exists, no matter how justifiable it is (e.g. the 20 kg threshold in Carnivora, which is related to prey size changes; Carbone et al., 1999).

**Figure 5.**
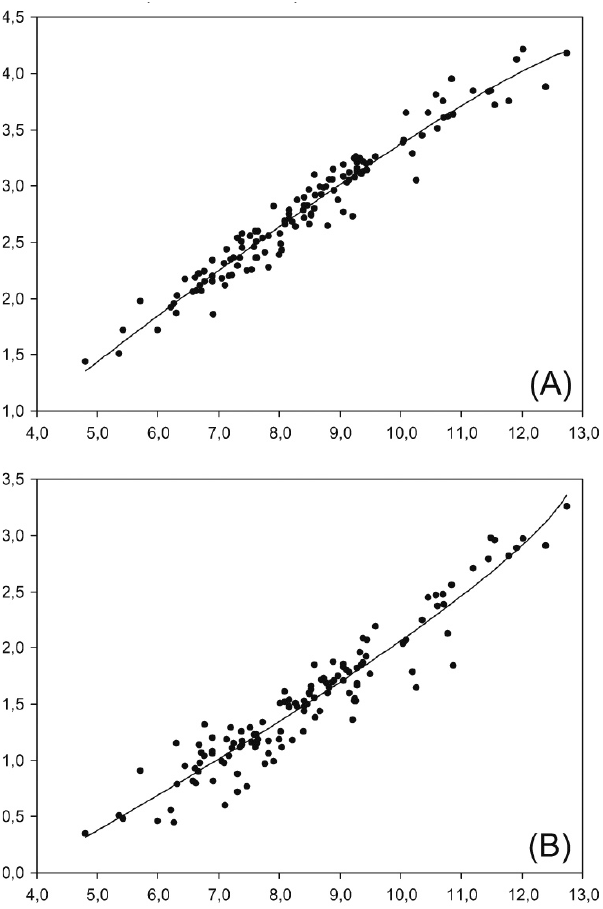
Differential scaling. Complex allometry plots for olecranon length (A) and ulna transverse diameter (B) in the fissiped subsample. As indicated by the curvature of the plot, olecranon length scales faster in small carnivorans than in large carnivorans (i.e., *D* > 1), while the opposite is true for ulna transverse diameter (i.e., *D* < 1).

Another source of variability is the adaptation to performing different modes of locomotion besides walking and running (climbing, swimming, digging). As stated in the introduction, Bou et al. (1987) suggested that similarity hypothesis imply adaptive neutrality, which is not the case, since the present study has proved that adaptations to different locomotor habits do indeed result in different scaling patterns. Furthermore, differences in locomotor habit within the same sample has been proposed as another possible explanation for differential scaling (Castiella & Casinos, 1990; Gálvez-López & Casinos, 2012).

Finally, at least in Carnivora, phylogenetic relatedness also plays an important role in limb bone scaling, as suggested by the different allometric exponents obtained with traditional and PIC regression methods in the present study (contrary to previous studies comparing both methodologies in this and other mammal groups; Christiansen, 2002*b*; Christiansen & Adolfssen, 2005; Gálvez-López & Casinos, 2012).

In conclusion, thus, we propose that either an overcomplicated model should be constructed including all these factors (and the ones we are probably missing), or we finally drop the “universal scaling” searching and focus on solving little problems one at atime, and from the sum of them formulate a generalization (if possible). For instance, how does limb bone morphology change with size in arboreal carnivorans? What about in arboreal didelphids and so on? Can we generalize all those scaling patterns into one scaling pattern for arboreal mammals? We consider that the present study constitutes a first step in that direction.

## Acknowledgements

We would like to thank the curators of the Phylogenetisches Museum (Jena), the Museum für Naturkunde (Berlin), the Museu de Ciències Naturals de la Ciutadella (Barcelona), the Muséum National d’Histoire Naturelle (Paris), the Museo Nacional de Ciencias Naturales (Madrid), the Museo Argentino de Ciencias Naturales (Buenos Aires), the Museo de La Plata (Argentina), and the Naturhistorisches Museum Basel (Switzerland), for granting me access to the collections. We would also show my appreciation to the following organisations for partially funding this research: la Caixa; Deutscher Akademischer Austausch Dienst (DAAD); the University of Barcelona (UB); Agència de Gestió d’Ajuts Universitaris i de Recerca (AGAUR); Departament d’Innovació, Universitats i Empresa de la Generalitat de Catalunya; and the European Social Fund (ESF). Finally, this work was completed with the assistance of funds from research grants CGL2005-04402/BOS and CGL2008-00832/BOS from the former Ministerio de Educación y Ciencia (MEC) of Spain.

