## Supplementary Material for "Scaling pattern of the carnivoran forelimb: Locomotor types, differential scaling and thoughts on a dying similarity"

**Table S1. Branch length transformations used for phylogenetically independent contrasts.** Variable names are listed in Table 3. Abbreviations: exp., exponential transformation; Gra., Grafen's transformation; ln, natural logarithm transformation; Nee, Nee's transformation; $\rho_x$, Grafen's rho transform, where x indicates the value of rho; untr., untransformed branch lengths.

| | | | family | | | | | | | locomotor type | | | | | | |
|---|---|---|---|---|---|---|---|---|---|---|---|---|---|---|---|---|
| | **whole sample** | **fissipeds** | **Canidae** | **Mustelidae** | **Procyonidae** | **Ursidae** | **Felidae** | **Herpestidae** | **Viverridae** | **arboreal** | **semiarboreal** | **scansorial** | **terrestrial** | **semifossorial** | **semiaquatic** | **aquatic** |
| **$L_s$** | $\rho_{0.5}$ | Nee | $\rho_{0.5}$ | $\rho_{0.5}$ | untr. | $\rho_{0.1}$ | $\rho_{0.5}$ | ln | Nee | exp. | $\rho_{0.8}$ | Nee | ln | $\rho_{0.3}$ | Nee | $\rho_{0.8}$ |
| **S** | $\rho_{0.5}$ | Nee | Nee | $\rho_{0.5}$ | untr. | $\rho_{0.1}$ | $\rho_{0.5}$ | ln | $\rho_{1.7}$ | exp. | $\rho_{0.8}$ | Nee | ln | untr. | Nee | untr. |
| **I** | Nee | Nee | $\rho_{0.5}$ | $\rho_{0.5}$ | ln | $\rho_{0.5}$ | $\rho_{0.5}$ | ln | $\rho_{1.7}$ | $\rho_{0.1}$ | $\rho_{0.8}$ | Nee | ln | $\rho_{0.3}$ | Nee | $\rho_{0.8}$ |
| **A** | Nee | Nee | $\rho_{0.5}$ | $\rho_{0.5}$ | untr. | $\rho_{0.1}$ | $\rho_{0.5}$ | ln | Nee | $\rho_{0.1}$ | $\rho_{0.8}$ | Nee | ln | $\rho_{0.3}$ | Nee | $\rho_{0.8}$ |
| **$H_S$** | $\rho_{0.5}$ | $\rho_{0.5}$ | Nee | $\rho_{0.5}$ | untr. | $\rho_{0.1}$ | $\rho_{0.5}$ | ln | Nee | $\rho_{0.1}$ | $\rho_{0.8}$ | Nee | ln | Gra. | Nee | ln |
| **$L_h$** | Nee | $\rho_{0.5}$ | ln | $\rho_{0.5}$ | untr. | $\rho_{0.1}$ | $\rho_{0.5}$ | $\rho_{0.5}$ | Nee | $\rho_{0.1}$ | $\rho_{0.8}$ | Nee | ln | $\rho_{0.3}$ | Nee | Nee |
| **$d_{sh}$** | Nee | $\rho_{0.5}$ | Nee | $\rho_{0.5}$ | untr. | $\rho_{0.1}$ | $\rho_{0.5}$ | untr. | Nee | $\rho_{0.1}$ | ln | Nee | ln | $\rho_{0.3}$ | exp. | ln |
| **$d_{th}$** | Nee | $\rho_{0.5}$ | ln | $\rho_{0.5}$ | Nee | $\rho_{0.1}$ | $\rho_{0.5}$ | untr. | $\rho_{1.7}$ | $\rho_{0.1}$ | $\rho_{0.8}$ | Nee | ln | $\rho_{0.3}$ | Nee | $\rho_{0.8}$ |
| **T** | exp. | exp. | Gra. | $\rho_{0.5}$ | $\rho_{0.5}$ | Nee | untr. | $\rho_{2.0}$ | $\rho_{1.7}$ | exp. | Nee | exp. | exp. | untr. | Gra. | untr. |
| **HR** | exp. | exp. | Nee | $\rho_{0.5}$ | untr. | $\rho_{0.5}$ | $\rho_{0.5}$ | untr. | Nee | $\rho_{0.1}$ | Gra. | Nee | $\rho_{0.5}$ | Gra. | Nee | $\rho_{0.8}$ |
| **$L_r$** | Nee | $\rho_{0.5}$ | ln | $\rho_{0.5}$ | Nee | $\rho_{0.1}$ | untr. | $\rho_{0.5}$ | $\rho_{1.7}$ | $\rho_{0.1}$ | $\rho_{0.8}$ | Nee | ln | $\rho_{0.3}$ | Gra. | $\rho_{0.8}$ |
| **$d_{sr}$** | Nee | Nee | $\rho_{0.5}$ | $\rho_{0.5}$ | untr. | $\rho_{0.1}$ | $\rho_{0.5}$ | untr. | $\rho_{1.7}$ | $\rho_{0.1}$ | $\rho_{0.8}$ | Nee | ln | $\rho_{0.3}$ | Nee | $\rho_{0.8}$ |
| **$d_{tr}$** | Nee | $\rho_{0.5}$ | $\rho_{0.5}$ | ln | Nee | $\rho_{0.5}$ | $\rho_{0.5}$ | $\rho_{0.5}$ | $\rho_{1.7}$ | $\rho_{0.1}$ | $\rho_{0.8}$ | Nee | ln | $\rho_{0.3}$ | Nee | $\rho_{0.8}$ |
| **P** | Nee | Nee | Nee | $\rho_{0.5}$ | untr. | Nee | $\rho_{0.5}$ | $\rho_{0.5}$ | Nee | $\rho_{0.1}$ | $\rho_{0.8}$ | Nee | exp. | untr. | Gra. | ln |
| **RR** | exp. | exp. | Nee | $\rho_{0.5}$ | untr. | $\rho_{0.1}$ | $\rho_{0.5}$ | ln | Nee | $\rho_{0.1}$ | ln | Nee | ln | untr. | exp. | Nee |
| **$L_u$** | Nee | $\rho_{0.5}$ | ln | $\rho_{0.5}$ | Nee | $\rho_{0.1}$ | $\rho_{0.5}$ | ln | $\rho_{1.7}$ | $\rho_{0.1}$ | $\rho_{0.8}$ | Nee | ln | $\rho_{0.3}$ | Gra. | $\rho_{0.8}$ |
| **$d_{su}$** | $\rho_{0.5}$ | $\rho_{0.5}$ | ln | $\rho_{0.5}$ | Nee | Nee | $\rho_{0.5}$ | $\rho_{0.5}$ | $\rho_{1.7}$ | $\rho_{0.1}$ | $\rho_{0.8}$ | Nee | ln | untr. | Nee | $\rho_{0.8}$ |
| **$d_{tu}$** | Nee | exp. | $\rho_{0.5}$ | $\rho_{0.5}$ | untr. | $\rho_{0.1}$ | $\rho_{0.5}$ | ln | $\rho_{1.7}$ | exp. | $\rho_{0.8}$ | Nee | ln | untr. | exp. | $\rho_{0.8}$ |
| **O** | $\rho_{0.5}$ | $\rho_{0.5}$ | Nee | $\rho_{0.5}$ | untr. | $\rho_{0.1}$ | $\rho_{0.5}$ | ln | Nee | exp. | $\rho_{0.8}$ | Nee | ln | untr. | Nee | ln |
| **θ** | exp. | exp. | Nee | exp. | $\rho_{0.5}$ | Nee | untr. | ln | exp. | exp. | Nee | $\rho_{0.5}$ | exp. | $\rho_{0.5}$ | exp. | ln |
| **α** | $\rho_{0.5}$ | $\rho_{0.5}$ | $\rho_{0.5}$ | ln | ln | $\rho_{0.1}$ | $\rho_{0.5}$ | ln | Nee | $\rho_{0.1}$ | ln | Nee | ln | $\rho_{0.3}$ | Nee | Nee |
| **UR** | $\rho_{0.5}$ | $\rho_{0.5}$ | Nee | ln | Nee | $\rho_{0.1}$ | $\rho_{0.5}$ | untr. | Nee | exp. | Nee | exp. | ln | Gra. | exp. | $\rho_{0.8}$ |
| **IFA** | $\rho_{0.5}$ | $\rho_{0.5}$ | ln | $\rho_{0.5}$ | Nee | exp. | $\rho_{0.5}$ | ln | Nee | exp. | $\rho_{0.8}$ | exp. | ln | $\rho_{0.3}$ | Nee | $\rho_{0.8}$ |
| **$L_m$** | Nee | $\rho_{0.5}$ | $\rho_{0.5}$ | $\rho_{0.5}$ | Nee | $\rho_{0.1}$ | $\rho_{0.5}$ | $\rho_{0.5}$ | $\rho_{1.7}$ | $\rho_{0.1}$ | ln | Nee | ln | Nee | exp. | $\rho_{0.8}$ |
| **$d_{sm}$** | Nee | Nee | ln | $\rho_{0.5}$ | untr. | Nee | $\rho_{0.5}$ | ln | Nee | $\rho_{0.1}$ | ln | Nee | ln | $\rho_{0.3}$ | Gra. | Nee |
| **$d_{tm}$** | Nee | $\rho_{0.5}$ | Nee | ln | ln | $\rho_{0.1}$ | $\rho_{0.5}$ | $\rho_{0.5}$ | $\rho_{1.7}$ | $\rho_{0.1}$ | $\rho_{0.8}$ | Nee | ln | $\rho_{0.3}$ | Gra. | Nee |
| **MR** | $\rho_{0.5}$ | $\rho_{0.5}$ | ln | $\rho_{0.5}$ | ln | $\rho_{0.1}$ | $\rho_{0.5}$ | $\rho_{0.5}$ | Nee | $\rho_{0.5}$ | Gra. | $\rho_{0.5}$ | ln | $\rho_{0.3}$ | Nee | ln |
| **$\%_{prox}$** | Nee | $\rho_{0.5}$ | Nee | ln | untr. | $\rho_{0.1}$ | $\rho_{0.5}$ | ln | Nee | $\rho_{0.1}$ | $\rho_{0.8}$ | $\rho_{0.5}$ | ln | Gra. | exp. | Nee |
| **$\%_{mid}$** | Nee | Nee | ln | Nee | untr. | $\rho_{0.5}$ | untr. | untr. | $\rho_{1.7}$ | exp. | Nee | Nee | ln | $\rho_{0.3}$ | exp. | Nee |
| **$\%_{dist}$** | $\rho_{0.5}$ | exp. | ln | exp. | Nee | $\rho_{0.1}$ | $\rho_{0.5}$ | untr. | exp. | $\rho_{0.1}$ | exp. | Nee | ln | Gra. | exp. | $\rho_{0.8}$ |

### Tables SR1 to SR30 – Results of traditional and PIC regressions

As indicated in Table 3, the following tables present the regression results for each variable. Both the results using traditional regression methods and phylogenetically independent contrasts (PIC) are shown for the whole sample and each of the subsamples (fissipeds, by Family, and by locomotor type). In each case, it is indicated (in the "sim." columns) whether the theoretical values proposed by the geometric similarity hypothesis (G), the elastic similarity hypothesis (E), or both (B), are included in the 95% confidence interval for the slope *b* (95% $CI_b$). Furthermore, when neither theoretical value is included in the 95% $CI_b$, it is indicated whether there is positive allometry (+; *b* is higher than both theoretical values), negative allometry (–; *b* is lower than both theoretical values), or both (nei.; *b* is higher than one theoretical values and lower than the other). Finally, the results of the comparison between the allometric coefficients obtained with each methodology are presented in the last column ($b_{trad} \neq b_{PIC}$): a cross (×) indicates no significant differences, while a tick (✓) denotes that the slopes are significantly different from each other ($p < 0.05$).

Variable names and abbreviations are given in Table 3, while the following abbreviations are common to all following tables: 95% $CI_a$, 95% confidence interval for the coefficient *a*; 95% $CI_b$, 95% confidence interval for the allometric coefficient *b*; n, sample size; n.s., unable to test differences due to non-significant regression; R, correlation coefficient; sim., similarity. Results in ***grey italics*** denote non-significant regressions.

| SR1 – $L_s$ | traditional regression | | | | | | | PIC regression | | | | | |
|---|---|---|---|---|---|---|---|---|---|---|---|---|---|
| | n | *a* | 95% CI$_a$ | $b_{trad}$ | 95% CI$_b$ | R | sim. | n | $b_{PIC}$ | 95% CI$_b$ | R | sim. | $b_{trad} \neq b_{PIC}$ |
| **whole sample** | 137 | 3.098 | 2.662 – 3.682 | 0.356 | 0.336 – 0.374 | 0.957 | + | 136 | 0.338 | 0.318 – 0.358 | 0.940 | G | × |
| **fissipeds** | 130 | 2.667 | 2.280 – 3.166 | 0.376 | 0.356 – 0.394 | 0.957 | + | 129 | 0.346 | 0.325 – 0.367 | 0.940 | G | ✓ |
| **Family** | | | | | | | | | | | | | |
| Canidae | 17 | 1.622 | 1.043 – 3.210 | 0.445 | 0.368 – 0.493 | 0.973 | + | 16 | 0.435 | 0.369 – 0.501 | 0.961 | + | × |
| Mustelidae | 32 | 2.970 | 2.214 – 3.985 | 0.341 | 0.304 – 0.378 | 0.956 | G | 31 | 0.330 | 0.283 – 0.377 | 0.923 | G | × |
| Procyonidae | 7 | 2.121 | 0.266 – 16.901 | 0.396 | 0.139 – 0.653 | 0.825 | B | 6 | 0.376 | 0.105 – 0.647 | 0.815 | B | × |
| Ursidae | 7 | 6.492 | 2.710 – 15.551 | 0.284 | 0.210 – 0.359 | 0.974 | B | 6 | 0.287 | 0.203 – 0.371 | 0.972 | B | × |
| Felidae | 26 | 5.271 | 4.163 – 6.674 | 0.313 | 0.288 – 0.338 | 0.982 | G | 25 | 0.317 | 0.283 – 0.351 | 0.968 | G | × |
| Herpestidae | 12 | 4.577 | 3.107 – 6.741 | 0.308 | 0.254 – 0.362 | 0.968 | G | 11 | 0.300 | 0.251 – 0.349 | 0.974 | G | × |
| Eupleridae | 5 | 4.173 | 0.881 – 19.767 | 0.329 | 0.115 – 0.543 | 0.935 | B | | | | | | |
| Viverridae | 14 | 5.077 | 2.677 – 9.630 | 0.299 | 0.220 – 0.378 | 0.907 | B | 13 | 0.313 | 0.223 – 0.403 | 0.893 | B | × |
| **Locomotor type** | | | | | | | | | | | | | |
| arboreal | 7 | 2.219 | 0.414 – 11.900 | 0.382 | 0.174 – 0.590 | 0.881 | B | 6 | 0.216 | 0.201 – 0.231 | 0.998 | – | × |
| semiarboreal | 10 | 5.179 | 3.296 – 8.138 | 0.301 | 0.242 – 0.359 | 0.971 | B | 9 | 0.288 | 0.185 – 0.391 | 0.903 | B | × |
| scansorial | 45 | 4.017 | 3.297 – 4.894 | 0.335 | 0.314 – 0.356 | 0.978 | G | 44 | 0.334 | 0.309 – 0.359 | 0.970 | G | × |
| terrestrial | 49 | 2.056 | 1.540 – 2.961 | 0.411 | 0.367 – 0.447 | 0.963 | + | 48 | 0.339 | 0.301 – 0.377 | 0.925 | G | ✓ |
| semifossorial | 7 | 3.086 | 1.197 – 7.958 | 0.345 | 0.226 – 0.464 | 0.954 | B | 6 | 0.355 | 0.220 – 0.490 | 0.952 | B | × |
| semiaquatic | 11 | 3.334 | 1.226 – 9.068 | 0.329 | 0.212 – 0.446 | 0.881 | B | 10 | 0.349 | 0.231 – 0.467 | 0.897 | B | × |
| aquatic | 8 | 3.484 | 0.494 – 24.560 | 0.324 | 0.161 – 0.488 | 0.863 | B | 7 | 0.358 | 0.154 – 0.562 | 0.841 | B | × |

| SR2 – S | traditional regression | | | | | | | PIC regression | | | | | |
|---|---|---|---|---|---|---|---|---|---|---|---|---|---|
| | n | *a* | 95% CI$_a$ | *b*$_{trad}$ | 95% CI$_b$ | R | sim. | n | *b*$_{PIC}$ | 95% CI$_b$ | R | sim. | *b*$_{trad}$ ≠ *b*$_{PIC}$ |
| **whole sample** | 137 | 0.897 | 0.728 – 1.112 | 0.378 | 0.351 – 0.403 | 0.967 | + | 136 | 0.348 | 0.326 – 0.370 | 0.930 | G | ✓ |
| **fissipeds** | 130 | 1.045 | 0.915 – 1.216 | 0.358 | 0.342 – 0.373 | 0.971 | + | 129 | 0.335 | 0.314 – 0.356 | 0.933 | G | ✓ |
| **Family** | | | | | | | | | | | | | |
| Canidae | 17 | 0.498 | 0.305 – 1.212 | 0.432 | 0.335 – 0.486 | 0.956 | + | 16 | 0.438 | 0.360 – 0.516 | 0.947 | + | × |
| Mustelidae | 32 | 1.089 | 0.838 – 1.416 | 0.354 | 0.321 – 0.388 | 0.967 | G | 31 | 0.337 | 0.291 – 0.383 | 0.932 | G | × |
| Procyonidae | 7 | 1.615 | 0.315 – 8.270 | 0.311 | 0.109 – 0.513 | 0.825 | B | *6* | *0.292* | *0.086 – 0.498* | *0.822* | *B* | n.s. |
| Ursidae | 7 | 5.843 | 3.181 – 10.732 | 0.212 | 0.161 – 0.264 | 0.977 | E | 6 | 0.215 | 0.158 – 0.272 | 0.977 | E | × |
| Felidae | 26 | 1.608 | 1.239 – 2.086 | 0.320 | 0.292 – 0.347 | 0.979 | G | 25 | 0.322 | 0.282 – 0.362 | 0.956 | G | × |
| Herpestidae | 12 | 1.430 | 1.110 – 1.842 | 0.308 | 0.272 – 0.343 | 0.987 | G | 11 | 0.298 | 0.260 – 0.336 | 0.984 | G | × |
| Eupleridae | 5 | 1.291 | 0.725 – 2.299 | 0.336 | 0.257 – 0.416 | 0.992 | G | | | | | | |
| Viverridae | 14 | 1.720 | 0.963 – 3.072 | 0.298 | 0.226 – 0.370 | 0.924 | B | *13* | *0.298* | *0.134 – 0.462* | *0.497* | *B* | n.s. |
| **Locomotor type** | | | | | | | | | | | | | |
| arboreal | 7 | 0.733 | 0.350 – 1.538 | 0.407 | 0.315 – 0.498 | 0.981 | G | 6 | 0.343 | 0.342 – 0.344 | 1.000 | + | × |
| semiarboreal | 10 | 1.614 | 0.972 – 2.680 | 0.318 | 0.252 – 0.384 | 0.967 | G | 9 | 0.302 | 0.188 – 0.416 | 0.892 | B | × |
| scansorial | 45 | 1.266 | 1.066 – 1.503 | 0.343 | 0.325 – 0.362 | 0.984 | G | 44 | 0.330 | 0.299 – 0.361 | 0.953 | G | × |
| terrestrial | 49 | 1.154 | 0.995 – 1.369 | 0.338 | 0.318 – 0.357 | 0.983 | G | 48 | 0.336 | 0.304 – 0.368 | 0.945 | G | × |
| semifossorial | 7 | 0.537 | 0.097 – 2.959 | 0.420 | 0.205 – 0.635 | 0.895 | B | 6 | 0.389 | 0.166 – 0.612 | 0.888 | B | × |
| semiaquatic | 11 | 1.038 | 0.545 – 1.977 | 0.362 | 0.287 – 0.438 | 0.961 | G | 10 | 0.401 | 0.295 – 0.507 | 0.939 | G | × |
| aquatic | *8* | *0.203* | *0.001 – 28.536* | *0.517* | *0.103 – 0.931* | *0.597* | *B* | *7* | *0.388* | *0.130 – 0.646* | *0.773* | *B* | n.s. |

| **SR3 – I** | **traditional regression** | | | | | | | **PIC regression** | | | | | |
|---|---|---|---|---|---|---|---|---|---|---|---|---|---|
| | **n** | ***a*** | **95% CI$_a$** | **$b_{trad}$** | **95% CI$_b$** | **R** | **sim.** | **n** | **$b_{PIC}$** | **95% CI$_b$** | **R** | **sim.** | **$b_{trad} \neq b_{PIC}$** |
| **whole sample** | 137 | 0.716 | 0.538 – 0.939 | 0.423 | 0.392 – 0.453 | 0.927 | + | 136 | 0.386 | 0.356 – 0.416 | 0.892 | + | ✓ |
| **fissipeds** | 130 | 0.614 | 0.460 – 0.806 | 0.443 | 0.412 – 0.474 | 0.924 | + | 129 | 0.384 | 0.358 – 0.410 | 0.922 | + | ✓ |
| **Family** | | | | | | | | | | | | | |
| Canidae | 17 | 0.978 | 0.509 – 2.697 | 0.399 | 0.286 – 0.472 | 0.933 | G | 16 | 0.418 | 0.324 – 0.512 | 0.914 | G | × |
| Mustelidae | 32 | 0.450 | 0.228 – 0.705 | 0.440 | 0.383 – 0.497 | 0.937 | + | 31 | 0.416 | 0.350 – 0.482 | 0.904 | + | × |
| Procyonidae | 7 | 4.282 | 1.564 – 11.724 | 0.233 | 0.108 – 0.357 | 0.885 | B | *6* | *0.238* | *0.066 – 0.410* | *0.812* | *B* | n.s. |
| Ursidae | 7 | 9.192 | 2.518 – 33.546 | 0.204 | 0.094 – 0.314 | 0.883 | E | *6* | *0.202* | *0.086 – 0.318* | *0.866* | *E* | n.s. |
| Felidae | 26 | 1.884 | 1.410 – 2.517 | 0.330 | 0.299 – 0.360 | 0.975 | G | 25 | 0.347 | 0.302 – 0.392 | 0.951 | G | × |
| Herpestidae | 12 | 1.830 | 1.083 – 3.094 | 0.316 | 0.243 – 0.390 | 0.944 | B | 11 | 0.317 | 0.248 – 0.386 | 0.953 | B | × |
| Eupleridae | 5 | 0.624 | 0.068 – 5.761 | 0.451 | 0.145 – 0.758 | 0.929 | B | | | | | | |
| Viverridae | 14 | 2.579 | 1.429 – 4.653 | 0.289 | 0.216 – 0.362 | 0.916 | B | 13 | 0.323 | 0.234 – 0.412 | 0.901 | B | × |
| **Locomotor type** | | | | | | | | | | | | | |
| arboreal | 7 | 1.561 | 0.735 – 3.316 | 0.357 | 0.264 – 0.450 | 0.974 | G | 6 | 0.357 | 0.255 – 0.459 | 0.973 | G | × |
| semiarboreal | 10 | 1.883 | 0.769 – 4.606 | 0.320 | 0.204 – 0.437 | 0.895 | B | 9 | 0.299 | 0.219 – 0.379 | 0.947 | B | × |
| scansorial | 45 | 1.113 | 0.850 – 1.458 | 0.385 | 0.356 – 0.414 | 0.969 | + | 44 | 0.385 | 0.340 – 0.430 | 0.925 | + | × |
| terrestrial | 49 | 0.450 | 0.294 – 0.763 | 0.477 | 0.418 – 0.526 | 0.932 | + | 48 | 0.362 | 0.317 – 0.407 | 0.907 | G | ✓ |
| semifossorial | 7 | 0.497 | 0.089 – 2.781 | 0.446 | 0.229 – 0.662 | 0.906 | B | 6 | 0.475 | 0.249 – 0.701 | 0.924 | B | × |
| semiaquatic | 11 | 0.425 | 0.085 – 2.115 | 0.449 | 0.261 – 0.637 | 0.832 | G | 10 | 0.459 | 0.293 – 0.625 | 0.883 | G | × |
| aquatic | *8* | *0.151* | *0.001 – 16.286* | *0.527* | *0.135 – 0.919* | *0.667* | *B* | *7* | *0.556* | *0.024 – 1.088* | *0.412* | *B* | n.s. |

| **SR4 – A** | **traditional regression** | | | | | | | **PIC regression** | | | | | |
|---|---|---|---|---|---|---|---|---|---|---|---|---|---|
| | **n** | ***a*** | **95% CI$_a$** | ***b*****trad** | **95% CI$_b$** | **R** | **sim.** | **n** | ***b*****PIC** | **95% CI$_b$** | **R** | **sim.** | ***b*****trad ≠ *b*****PIC** |
| **whole sample** | 137 | 1.774 | 1.500 – 2.112 | 0.392 | 0.373 – 0.410 | 0.970 | + | 136 | 0.355 | 0.333 – 0.377 | 0.931 | G | ✓ |
| **fissipeds** | 130 | 1.785 | 1.518 – 2.087 | 0.391 | 0.374 – 0.409 | 0.965 | + | 129 | 0.353 | 0.332 – 0.374 | 0.940 | G | ✓ |
| **Family** | | | | | | | | | | | | | |
| Canidae | 17 | 1.391 | 0.776 – 3.607 | 0.419 | 0.316 – 0.483 | 0.954 | G | 16 | 0.428 | 0.348 – 0.508 | 0.941 | + | × |
| Mustelidae | 32 | 1.564 | 1.134 – 2.157 | 0.387 | 0.346 – 0.428 | 0.959 | + | 31 | 0.366 | 0.313 – 0.414 | 0.921 | G | × |
| Procyonidae | 7 | 4.772 | 1.770 – 12.868 | 0.285 | 0.162 – 0.408 | 0.927 | B | *6* | *0.286* | *0.111 – 0.461* | *0.870* | *B* | n.s. |
| Ursidae | 7 | 18.339 | 8.036 – 41.855 | 0.191 | 0.121 – 0.261 | 0.948 | E | 6 | 0.191 | 0.116 – 0.266 | 0.949 | E | × |
| Felidae | 26 | 3.476 | 2.743 – 4.405 | 0.327 | 0.302 – 0.352 | 0.983 | G | 25 | 0.335 | 0.298 – 0.372 | 0.965 | G | × |
| Herpestidae | 12 | 3.643 | 2.675 – 4.961 | 0.301 | 0.258 – 0.344 | 0.979 | G | 11 | 0.300 | 0.255 – 0.345 | 0.978 | G | × |
| Eupleridae | 5 | 2.216 | 0.745 – 6.586 | 0.369 | 0.218 – 0.519 | 0.975 | B | | | | | | |
| Viverridae | 14 | 4.444 | 2.757 – 7.164 | 0.288 | 0.229 – 0.347 | 0.945 | B | 13 | 0.297 | 0.237 – 0.357 | 0.948 | B | × |
| **Locomotor type** | | | | | | | | | | | | | |
| arboreal | 7 | 2.544 | 1.433 – 4.517 | 0.360 | 0.288 – 0.431 | 0.985 | G | 6 | 0.358 | 0.278 – 0.438 | 0.984 | G | × |
| semiarboreal | 10 | 4.124 | 2.636 – 6.452 | 0.300 | 0.241 – 0.358 | 0.971 | B | 9 | 0.277 | 0.210 – 0.344 | 0.958 | B | × |
| scansorial | 45 | 2.386 | 1.964 – 2.899 | 0.365 | 0.344 – 0.386 | 0.982 | + | 44 | 0.360 | 0.325 – 0.395 | 0.951 | G | × |
| terrestrial | 49 | 1.612 | 1.281 – 2.121 | 0.399 | 0.369 – 0.425 | 0.969 | + | 48 | 0.345 | 0.309 – 0.381 | 0.935 | G | ✓ |
| semifossorial | 7 | 1.304 | 0.392 – 4.336 | 0.409 | 0.258 – 0.560 | 0.947 | G | 6 | 0.420 | 0.262 – 0.578 | 0.953 | G | × |
| semiaquatic | 11 | 1.449 | 0.598 – 3.512 | 0.399 | 0.295 – 0.502 | 0.939 | G | 10 | 0.425 | 0.317 – 0.533 | 0.943 | G | × |
| aquatic | 8 | 0.997 | 0.038 – 26.417 | 0.439 | 0.165 – 0.714 | 0.780 | B | *7* | *0.373* | *0.090 – 0.656* | *0.693* | *B* | n.s. |

| SR5 – $H_S$ | traditional regression | | | | | | | PIC regression | | | | | |
|---|---|---|---|---|---|---|---|---|---|---|---|---|---|
| | n | *a* | 95% $CI_a$ | $b_{trad}$ | 95% $CI_b$ | R | sim. | n | $b_{PIC}$ | 95% $CI_b$ | R | sim. | $b_{trad} \neq b_{PIC}$ |
| **whole sample** | 137 | 0.171 | 0.100 – 0.292 | 0.454 | 0.393 – 0.515 | 0.822 | + | 136 | 0.463 | 0.416 – 0.510 | 0.804 | + | × |
| **fissipeds** | 130 | 0.114 | 0.068 – 0.198 | 0.510 | 0.449 – 0.568 | 0.897 | + | 129 | 0.474 | 0.429 – 0.519 | 0.839 | + | × |
| **Family** | | | | | | | | | | | | | |
| Canidae | 17 | 0.166 | 0.100 – 0.440 | 0.480 | 0.374 – 0.534 | 0.947 | + | 16 | 0.441 | 0.353 – 0.529 | 0.933 | + | × |
| Mustelidae | 32 | 0.073 | 0.032 – 0.165 | 0.548 | 0.444 – 0.653 | 0.860 | + | 31 | 0.497 | 0.384 – 0.610 | 0.791 | + | × |
| Procyonidae | 7 | 0.111 | 0.008 – 1.459 | 0.531 | 0.212 – 0.850 | 0.852 | B | *6* | *0.523* | *0.152 – 0.894* | *0.821* | *B* | n.s. |
| Ursidae | 7 | 2.019 | 0.341 – 11.971 | 0.237 | 0.086 – 0.389 | 0.831 | B | *6* | *0.235* | *0.071 – 0.399* | *0.829* | *B* | n.s. |
| Felidae | 26 | 0.320 | 0.192 – 0.532 | 0.407 | 0.353 – 0.461 | 0.949 | + | 25 | 0.437 | 0.361 – 0.513 | 0.913 | + | × |
| Herpestidae | 12 | 0.827 | 0.434 – 1.573 | 0.278 | 0.188 – 0.368 | 0.887 | B | 11 | 0.250 | 0.158 – 0.342 | 0.858 | B | × |
| Eupleridae | 5 | 0.310 | 0.036 – 2.665 | 0.397 | 0.101 – 0.693 | 0.914 | B | | | | | | |
| Viverridae | 14 | 0.223 | 0.102 – 0.484 | 0.431 | 0.335 – 0.528 | 0.935 | + | 13 | 0.450 | 0.329 – 0.571 | 0.906 | G | × |
| **Locomotor type** | | | | | | | | | | | | | |
| arboreal | 7 | 0.219 | 0.044 – 1.098 | 0.429 | 0.229 – 0.628 | 0.914 | B | 6 | 0.425 | 0.215 – 0.635 | 0.917 | B | × |
| semiarboreal | 10 | 0.220 | 0.084 – 0.576 | 0.442 | 0.317 – 0.567 | 0.938 | G | 9 | 0.441 | 0.232 – 0.650 | 0.825 | B | × |
| scansorial | 45 | 0.270 | 0.201 – 0.364 | 0.420 | 0.388 – 0.452 | 0.968 | + | 44 | 0.435 | 0.384 – 0.486 | 0.924 | + | × |
| terrestrial | 49 | 0.096 | 0.044 – 0.224 | 0.535 | 0.434 – 0.625 | 0.916 | + | 48 | 0.455 | 0.388 – 0.522 | 0.866 | + | × |
| semifossorial | *7* | *0.005* | *0.000 – 2.586* | *0.866* | *0.089 – 1.644* | *0.626* | *B* | *6* | *0.564* | *0.068 – 1.060* | *0.706* | *B* | n.s. |
| semiaquatic | 11 | 0.024 | 0.002 – 0.369 | 0.667 | 0.349 – 0.985 | 0.775 | + | *10* | *0.847* | *0.338 – 1.356* | *0.623* | *+* | n.s. |
| aquatic | *8* | *0.038* | *0.000 – 11.844* | *0.483* | *0.000 – 0.965* | *0.004* | *B* | *7* | *0.580* | *-0.007 – 1.167* | *0.263* | *B* | n.s. |

| **SR6 – $L_h$** | **traditional regression** | | | | | | | **PIC regression** | | | | | |
|---|---|---|---|---|---|---|---|---|---|---|---|---|---|
| | **n** | ***a*** | **95% CI$_a$** | **$b_{trad}$** | **95% CI$_b$** | **R** | **sim.** | **n** | **$b_{PIC}$** | **95% CI$_b$** | **R** | **sim.** | **$b_{trad} \neq b_{PIC}$** |
| **whole sample** | 137 | 5.994 | 4.878 – 7.355 | 0.318 | 0.293 – 0.344 | 0.909 | G | 136 | 0.311 | 0.289 – 0.333 | 0.911 | nei. | × |
| **fissipeds** | 130 | 4.580 | 3.959 – 5.399 | 0.354 | 0.334 – 0.370 | 0.946 | + | 129 | 0.311 | 0.290 – 0.332 | 0.924 | nei. | ✓ |
| **Family** | | | | | | | | | | | | | |
| Canidae | 17 | 3.350 | 1.705 – 11.350 | 0.401 | 0.263 – 0.478 | 0.920 | G | 16 | 0.415 | 0.314 – 0.516 | 0.898 | G | × |
| Mustelidae | 32 | 6.018 | 4.208 – 8.608 | 0.298 | 0.252 – 0.344 | 0.912 | G | 31 | 0.292 | 0.244 – 0.340 | 0.897 | B | × |
| Procyonidae | 7 | 13.706 | 5.597 – 33.563 | 0.224 | 0.113 – 0.335 | 0.902 | B | *6* | ***0.219*** | ***0.081 – 0.357*** | ***0.861*** | ***B*** | n.s. |
| Ursidae | 7 | 13.772 | 6.344 – 29.899 | 0.252 | 0.186 – 0.318 | 0.974 | E | 6 | 0.256 | 0.186 – 0.326 | 0.975 | E | × |
| Felidae | 26 | 9.680 | 7.805 – 12.006 | 0.285 | 0.263 – 0.308 | 0.982 | nei. | 25 | 0.295 | 0.264 – 0.326 | 0.969 | nei. | × |
| Herpestidae | 12 | 6.732 | 3.892 – 11.644 | 0.299 | 0.222 – 0.375 | 0.931 | B | 11 | 0.285 | 0.213 – 0.357 | 0.935 | B | × |
| Eupleridae | 5 | 5.553 | 3.834 – 8.042 | 0.338 | 0.287 – 0.389 | 0.997 | G | | | | | | |
| Viverridae | 14 | 11.070 | 6.159 – 19.896 | 0.248 | 0.175 – 0.321 | 0.885 | E | 13 | 0.282 | 0.204 – 0.360 | 0.901 | B | × |
| **Locomotor type** | | | | | | | | | | | | | |
| arboreal | 7 | 5.956 | 2.087 – 16.997 | 0.334 | 0.204 – 0.463 | 0.941 | B | 6 | 0.334 | 0.193 – 0.475 | 0.940 | B | × |
| semiarboreal | 10 | 10.101 | 6.958 – 14.664 | 0.270 | 0.222 – 0.319 | 0.975 | E | 9 | 0.241 | 0.147 – 0.335 | 0.884 | B | × |
| scansorial | 45 | 6.878 | 5.800 – 8.156 | 0.316 | 0.298 – 0.335 | 0.982 | G | 44 | 0.314 | 0.289 – 0.339 | 0.966 | G | × |
| terrestrial | 49 | 3.645 | 2.899 – 5.137 | 0.381 | 0.338 – 0.408 | 0.959 | + | 48 | 0.323 | 0.282 – 0.364 | 0.900 | G | ✓ |
| semifossorial | 7 | 4.750 | 1.793 – 12.584 | 0.324 | 0.201 – 0.446 | 0.944 | B | 6 | 0.336 | 0.197 – 0.475 | 0.942 | B | × |
| semiaquatic | 11 | 8.410 | 3.903 – 18.128 | 0.256 | 0.166 – 0.346 | 0.885 | B | 10 | 0.269 | 0.183 – 0.355 | 0.911 | B | × |
| aquatic | 8 | 10.127 | 1.925 – 53.281 | 0.222 | 0.083 – 0.361 | 0.779 | B | *7* | ***0.237*** | ***0.090 – 0.384*** | ***0.808*** | ***B*** | n.s. |

| **SR7 – $d_{sh}$** | **traditional regression** | | | | | | | **PIC regression** | | | | | |
|---|---|---|---|---|---|---|---|---|---|---|---|---|---|
| | **n** | ***a*** | **95% CI$_a$** | **$b_{trad}$** | **95% CI$_b$** | **R** | **sim.** | **n** | **$b_{PIC}$** | **95% CI$_b$** | **R** | **sim.** | **$b_{trad} \neq b_{PIC}$** |
| **whole sample** | 137 | 0.396 | 0.352 – 0.454 | 0.376 | 0.362 – 0.389 | 0.975 | E | 136 | 0.383 | 0.359 – 0.407 | 0.933 | E | × |
| **fissipeds** | 130 | 0.371 | 0.329 – 0.432 | 0.385 | 0.368 – 0.399 | 0.971 | E | 129 | 0.387 | 0.361 – 0.413 | 0.927 | E | × |
| **Family** | | | | | | | | | | | | | |
| Canidae | 17 | 0.138 | 0.071 – 0.467 | 0.488 | 0.355 – 0.560 | 0.959 | E | 16 | 0.454 | 0.372 – 0.536 | 0.945 | E | × |
| Mustelidae | 32 | 0.326 | 0.226 – 0.469 | 0.399 | 0.352 – 0.445 | 0.950 | E | 31 | 0.386 | 0.319 – 0.453 | 0.883 | B | × |
| Procyonidae | 7 | 0.496 | 0.117 – 2.092 | 0.370 | 0.191 – 0.548 | 0.908 | B | *6* | *0.366* | *0.147 – 0.585* | *0.876* | *B* | n.s. |
| Ursidae | 7 | 0.962 | 0.607 – 1.522 | 0.293 | 0.254 – 0.332 | 0.993 | – | 6 | 0.295 | 0.252 – 0.338 | 0.993 | G | × |
| Felidae | 26 | 0.353 | 0.271 – 0.459 | 0.393 | 0.365 – 0.421 | 0.986 | E | 25 | 0.382 | 0.341 – 0.423 | 0.968 | E | × |
| Herpestidae | 12 | 0.676 | 0.424 – 1.078 | 0.302 | 0.237 – 0.368 | 0.952 | G | 11 | 0.322 | 0.262 – 0.382 | 0.966 | B | × |
| Eupleridae | 5 | 0.151 | 0.043 – 0.529 | 0.515 | 0.342 – 0.688 | 0.983 | E | | | | | | |
| Viverridae | 14 | 0.474 | 0.209 – 1.077 | 0.349 | 0.248 – 0.451 | 0.887 | B | 13 | 0.338 | 0.263 – 0.413 | 0.937 | B | × |
| **Locomotor type** | | | | | | | | | | | | | |
| arboreal | 7 | 0.443 | 0.269 – 0.730 | 0.375 | 0.313 – 0.437 | 0.990 | B | 6 | 0.371 | 0.305 – 0.437 | 0.990 | B | × |
| semiarboreal | 10 | 0.320 | 0.189 – 0.542 | 0.416 | 0.347 – 0.485 | 0.979 | E | 9 | 0.418 | 0.333 – 0.503 | 0.970 | B | × |
| scansorial | 45 | 0.374 | 0.302 – 0.465 | 0.384 | 0.360 – 0.407 | 0.980 | E | 44 | 0.382 | 0.347 – 0.417 | 0.956 | E | × |
| terrestrial | 49 | 0.372 | 0.300 – 0.500 | 0.381 | 0.347 – 0.406 | 0.968 | E | 48 | 0.354 | 0.312 – 0.396 | 0.917 | B | × |
| semifossorial | 7 | 0.502 | 0.211 – 1.197 | 0.337 | 0.227 – 0.446 | 0.959 | B | 6 | 0.350 | 0.211 – 0.489 | 0.948 | B | × |
| semiaquatic | 11 | 0.294 | 0.094 – 0.922 | 0.418 | 0.284 – 0.552 | 0.906 | B | 10 | 0.527 | 0.279 – 0.775 | 0.791 | B | × |
| aquatic | 8 | 0.249 | 0.025 – 2.525 | 0.407 | 0.213 – 0.601 | 0.879 | B | 7 | 0.443 | 0.261 – 0.625 | 0.920 | B | n.s. |

| SR8 – $d_{th}$ | traditional regression | | | | | | | PIC regression | | | | | |
|---|---|---|---|---|---|---|---|---|---|---|---|---|---|
| | n | *a* | 95% $CI_a$ | $b_{trad}$ | 95% $CI_b$ | R | sim. | n | $b_{PIC}$ | 95% $CI_b$ | R | sim. | $b_{trad} \neq b_{PIC}$ |
| **whole sample** | 137 | 0.319 | 0.281 – 0.365 | 0.369 | 0.354 – 0.384 | 0.973 | E | 136 | 0.347 | 0.324 – 0.370 | 0.923 | G | × |
| **fissipeds** | 130 | 0.322 | 0.285 – 0.372 | 0.368 | 0.353 – 0.383 | 0.968 | E | 129 | 0.341 | 0.319 – 0.363 | 0.931 | G | ✓ |
| **Family** | | | | | | | | | | | | | |
| Canidae | 17 | 0.185 | 0.084 – 0.505 | 0.430 | 0.320 – 0.517 | 0.941 | B | 16 | 0.426 | 0.328 – 0.524 | 0.910 | B | × |
| Mustelidae | 32 | 0.466 | 0.348 – 0.623 | 0.307 | 0.270 – 0.344 | 0.946 | G | 31 | 0.319 | 0.270 – 0.368 | 0.910 | G | × |
| Procyonidae | 7 | 0.810 | 0.370 – 1.772 | 0.267 | 0.170 – 0.364 | 0.949 | G | 6 | 0.261 | 0.167 – 0.355 | 0.957 | G | × |
| Ursidae | 7 | 2.289 | 0.776 – 6.751 | 0.208 | 0.116 – 0.300 | 0.923 | – | 6 | 0.209 | 0.109 – 0.309 | 0.922 | – | × |
| Felidae | 26 | 0.381 | 0.288 – 0.505 | 0.354 | 0.324 – 0.384 | 0.980 | B | 25 | 0.339 | 0.301 – 0.377 | 0.964 | B | × |
| Herpestidae | 12 | 0.811 | 0.494 – 1.330 | 0.242 | 0.173 – 0.312 | 0.913 | – | 11 | 0.254 | 0.189 – 0.319 | 0.934 | – | × |
| Eupleridae | 5 | 0.204 | 0.084 – 0.496 | 0.439 | 0.316 – 0.561 | 0.988 | B | | | | | | |
| Viverridae | 14 | 0.430 | 0.225 – 0.825 | 0.331 | 0.251 – 0.412 | 0.922 | B | 13 | 0.488 | 0.308 – 0.668 | 0.814 | B | × |
| **Locomotor type** | | | | | | | | | | | | | |
| arboreal | 7 | 0.533 | 0.280 – 1.015 | 0.318 | 0.238 – 0.398 | 0.976 | B | 6 | 0.315 | 0.228 – 0.402 | 0.975 | B | × |
| semiarboreal | 10 | 0.363 | 0.184 – 0.716 | 0.362 | 0.274 – 0.451 | 0.954 | B | 9 | 0.374 | 0.233 – 0.515 | 0.892 | B | × |
| scansorial | 45 | 0.303 | 0.253 – 0.363 | 0.379 | 0.359 – 0.398 | 0.986 | E | 44 | 0.349 | 0.320 – 0.378 | 0.963 | B | ✓ |
| terrestrial | 49 | 0.335 | 0.281 – 0.426 | 0.361 | 0.334 – 0.382 | 0.967 | E | 48 | 0.342 | 0.301 – 0.383 | 0.915 | B | × |
| semifossorial | 7 | 0.432 | 0.196 – 0.954 | 0.322 | 0.222 – 0.421 | 0.963 | B | 6 | 0.335 | 0.188 – 0.482 | 0.935 | B | × |
| semiaquatic | 11 | 0.671 | 0.312 – 1.444 | 0.263 | 0.173 – 0.353 | 0.892 | G | 10 | 0.307 | 0.194 – 0.420 | 0.878 | B | × |
| aquatic | 8 | 0.084 | 0.005 – 1.511 | 0.479 | 0.237 – 0.720 | 0.863 | B | *7* | ***0.450*** | ***0.152 – 0.748*** | ***0.775*** | ***B*** | n.s. |

| **SR9 – T** | traditional regression | | | | | | | PIC regression | | | | | |
|---|---|---|---|---|---|---|---|---|---|---|---|---|---|
| | **n** | ***a*** | **95% CI$_a$** | **$b_{trad}$** | **95% CI$_b$** | **R** | **sim.** | **n** | **$b_{PIC}$** | **95% CI$_b$** | **R** | **sim.** | **$b_{trad} \neq b_{PIC}$** |
| **whole sample** | 137 | 0.558 | 0.162 – 1.071 | $6.14 \cdot 10^{-5}$ | $2.68 \cdot 10^{-5} – 7.63 \cdot 10^{-5}$ | 0.619 | + | 136 | $4.12 \cdot 10^{-5}$ | $3.43 \cdot 10^{-5} – 4.81 \cdot 10^{-5}$ | 0.190 | + | × |
| **fissipeds** | 130 | 0.447 | 0.137 – 0.927 | $6.56 \cdot 10^{-5}$ | $4.68 \cdot 10^{-6} – 9.11 \cdot 10^{-5}$ | 0.251 | + | *129* | *$4.22 \cdot 10^{-5}$* | *$3.49 \cdot 10^{-6} – 4.95 \cdot 10^{-5}$* | *0.157* | *+* | n.s. |
| **Family** | | | | | | | | | | | | | |
| Canidae | 17 | 1.560 | 0.840 – 2.499 | $1.88 \cdot 10^{-4}$ | $6.55 \cdot 10^{-5} – 2.55 \cdot 10^{-4}$ | 0.763 | + | 16 | $1.34 \cdot 10^{-4}$ | $7.12 \cdot 10^{-5} – 1.97 \cdot 10^{-4}$ | 0.530 | + | × |
| Mustelidae | *32* | *0.545* | *0.127 – 1.824* | *$-1.17 \cdot 10^{-4}$* | *$-4.22 \cdot 10^{-4} – -7.62 \cdot 10^{-5}$* | *0.288* | *–* | 31 | $-1.19 \cdot 10^{-4}$ | $-1.59 \cdot 10^{-4} – -7.87 \cdot 10^{-5}$ | 0.419 | – | n.s. |
| Procyonidae | *7* | *-1.081* | *-2.525 – 0.362* | *$3.00 \cdot 10^{-4}$* | *$2.72 \cdot 10^{-6} – 0.001$* | *0.174* | *+* | *6* | *$3.34 \cdot 10^{-4}$* | *$-7.96 \cdot 10^{-5} – 7.48 \cdot 10^{-4}$* | *0.075* | *B* | n.s. |
| Ursidae | *7* | *-2.861* | *-6.636 – 0.913* | *$2.09 \cdot 10^{-5}$* | *$-3.87 \cdot 10^{-5} – 5.91 \cdot 10^{-5}$* | *0.462* | *B* | *6* | *$2.25 \cdot 10^{-5}$* | *$-3.93 \cdot 10^{-6} – 4.89 \cdot 10^{-5}$* | *0.327* | *B* | n.s. |
| Felidae | 26 | 1.374 | 0.832 – 1.916 | $5.71 \cdot 10^{-5}$ | $1.38 \cdot 10^{-5} – 7.08 \cdot 10^{-5}$ | 0.892 | + | 25 | $5.47 \cdot 10^{-5}$ | $4.24 \cdot 10^{-5} – 6.70 \cdot 10^{-5}$ | 0.847 | + | × |
| Herpestidae | 12 | 0.248 | -0.187 – 0.683 | $4.50 \cdot 10^{-4}$ | $3.00 \cdot 10^{-4} – 0.001$ | 0.710 | + | *11* | *$4.42 \cdot 10^{-4}$* | *$1.75 \cdot 10^{-4} – 7.09 \cdot 10^{-4}$* | *0.530* | *+* | n.s. |
| Eupleridae | *5* | *1.408* | *-0.147 – 6.976* | *$-5.63 \cdot 10^{-4}$* | *$-0.005 – -3.75 \cdot 10^{-4}$* | *0.271* | *–* | | | | | | |
| Viverridae | 14 | -0.339 | -1.351 – 0.674 | $3.28 \cdot 10^{-4}$ | $1.13 \cdot 10^{-4} – 8.69 \cdot 10^{-4}$ | 0.560 | + | 13 | $2.93 \cdot 10^{-4}$ | $1.50 \cdot 10^{-4} – 4.36 \cdot 10^{-4}$ | 0.637 | + | × |
| **Locomotor type** | | | | | | | | | | | | | |
| arboreal | *7* | *-0.584* | *-5.474 – 1.021* | *$1.71 \cdot 10^{-4}$* | *$-6.07 \cdot 10^{-4} – 0.002$* | *0.063* | *B* | 6 | $6.30 \cdot 10^{-5}$ | $4.10 \cdot 10^{-5} – 8.50 \cdot 10^{-5}$ | 0.960 | + | n.s. |
| semiarboreal | *10* | *-0.769* | *-2.612 – -0.087* | *$2.55 \cdot 10^{-4}$* | *$4.97 \cdot 10^{-5} – 9.93 \cdot 10^{-4}$* | *0.508* | *+* | *9* | *$2.53 \cdot 10^{-4}$* | *$1.02 \cdot 10^{-4} – 4.04 \cdot 10^{-4}$* | *0.702* | *+* | n.s. |
| scansorial | 45 | 0.233 | -0.487 – 0.929 | $4.64 \cdot 10^{-5}$ | $2.16 \cdot 10^{-5} – 6.63 \cdot 10^{-5}$ | 0.440 | + | 44 | $5.14 \cdot 10^{-5}$ | $4.33 \cdot 10^{-5} – 5.95 \cdot 10^{-5}$ | 0.861 | + | × |
| terrestrial | *49* | *1.596* | *-0.963 – 3.215* | *$7.61 \cdot 10^{-5}$* | *$-2.15 \cdot 10^{-4} – 1.99 \cdot 10^{-4}$* | *0.131* | *B* | 48 | $1.31 \cdot 10^{-4}$ | $1.10 \cdot 10^{-4} – 1.52 \cdot 10^{-4}$ | 0.838 | + | n.s. |
| semifossorial | 7 | -0.264 | -0.728 – 0.151 | $1.96 \cdot 10^{-4}$ | $1.34 \cdot 10^{-4} – 2.90 \cdot 10^{-4}$ | 0.898 | + | *6* | *$1.89 \cdot 10^{-4}$* | *$7.46 \cdot 10^{-5} – 3.03 \cdot 10^{-4}$* | *0.873* | *+* | n.s. |
| semiaquatic | *11* | *0.705* | *-0.246 – 2.327* | *$-9.49 \cdot 10^{-5}$* | *$-2.74 \cdot 10^{-4} – 7.69 \cdot 10^{-5}$* | *0.115* | *B* | *10* | *$-8.41 \cdot 10^{-5}$* | *$-1.47 \cdot 10^{-4} – -2.11 \cdot 10^{-5}$* | *0.229* | *–* | n.s. |
| aquatic | *8* | *1.444* | *-21.620 – 14.430* | *$5.67 \cdot 10^{-5}$* | *$-2.95 \cdot 10^{-5} – 1.92 \cdot 10^{-4}$* | *0.642* | *B* | *7* | *$4.75 \cdot 10^{-5}$* | *$1.82 \cdot 10^{-5} – 7.68 \cdot 10^{-5}$* | *0.809* | *+* | n.s. |

| SR10 – HR | traditional regression | | | | | | | PIC regression | | | | | |
|---|---|---|---|---|---|---|---|---|---|---|---|---|---|
| | n | *a* | 95% CI$_a$ | $b_{trad}$ | 95% CI$_b$ | R | sim. | n | $b_{PIC}$ | 95% CI$_b$ | R | sim. | $b_{trad} \neq b_{PIC}$ |
| **whole sample** | 137 | 0.029 | 0.023 – 0.036 | 0.156 | 0.128 – 0.184 | 0.499 | + | 136 | 0.221 | 0.186 – 0.256 | 0.388 | + | ✓ |
| **fissipeds** | 130 | 0.034 | 0.028 – 0.044 | 0.132 | 0.101 – 0.157 | 0.289 | E | 129 | 0.222 | 0.186 – 0.258 | 0.381 | + | ✓ |
| **Family** | | | | | | | | | | | | | |
| Canidae | 17 | 0.015 | $4.19 \cdot 10^{-4}$ – 0.046 | 0.198 | 0.071 – 0.589 | 0.500 | E | 16 | 0.216 | 0.097 – 0.335 | 0.073 | E | n.s. |
| Mustelidae | 32 | 0.028 | 0.018 – 0.047 | 0.186 | 0.125 – 0.240 | 0.575 | E | 31 | 0.181 | 0.120 – 0.242 | 0.431 | E | × |
| Procyonidae | 7 | 0.024 | $4.28 \cdot 10^{-4}$ – 0.071 | 0.200 | 0.060 – 0.674 | 0.674 | E | 6 | 0.201 | 0.014 – 0.388 | 0.662 | E | n.s. |
| Ursidae | 7 | 0.046 | 0.004 – 0.106 | 0.076 | 0.005 – 0.292 | 0.575 | E | 6 | 0.073 | -0.006 – 0.152 | 0.495 | B | n.s. |
| Felidae | 26 | 0.030 | 0.023 – 0.040 | 0.130 | 0.098 – 0.154 | 0.823 | E | 25 | 0.134 | 0.090 – 0.178 | 0.626 | E | × |
| Herpestidae | 12 | 0.030 | 0.002 – 0.079 | 0.174 | 0.030 – 0.563 | 0.055 | E | 11 | 0.168 | 0.055 – 0.281 | 0.335 | E | n.s. |
| Eupleridae | 5 | 0.025 | 0.013 – 0.189 | 0.190 | -0.118 – 0.275 | 0.909 | B | | | | | | |
| Viverridae | 14 | 0.031 | 0.020 – 0.067 | 0.141 | 0.043 – 0.193 | 0.651 | E | 13 | 0.140 | 0.061 – 0.219 | 0.456 | E | n.s. |
| **Locomotor type** | | | | | | | | | | | | | |
| arboreal | 7 | 0.034 | $5.96 \cdot 10^{-4}$ – 0.150 | 0.140 | -0.059 – 0.646 | 0.408 | B | 6 | 0.124 | $-3.60 \cdot 10^{-4}$ – 0.248 | 0.584 | B | n.s. |
| semiarboreal | 10 | 0.024 | 0.013 – 0.057 | 0.180 | 0.069 – 0.252 | 0.806 | E | 9 | 0.246 | 0.097 – 0.395 | 0.690 | E | × |
| scansorial | 45 | 0.038 | 0.032 – 0.048 | 0.105 | 0.080 – 0.125 | 0.615 | E | 44 | 0.128 | 0.094 – 0.162 | 0.484 | E | × |
| terrestrial | 49 | 0.042 | 0.006 – 0.055 | 0.109 | 0.079 – 0.362 | 0.028 | E | 48 | 0.134 | 0.096 – 0.172 | 0.265 | E | n.s. |
| semifossorial | 7 | 0.064 | 0.014 – 0.092 | 0.076 | 0.021 – 0.260 | 0.227 | E | 6 | 0.064 | -0.013 – 0.141 | 0.273 | B | n.s. |
| semiaquatic | 11 | 0.015 | 0.006 – 0.115 | 0.260 | 0.022 – 0.367 | 0.583 | E | 10 | 0.291 | 0.111 – 0.471 | 0.592 | E | n.s. |
| aquatic | 8 | 0.007 | $3.94 \cdot 10^{-5}$ – 0.603 | 0.296 | -0.058 – 0.744 | 0.625 | B | 7 | 0.261 | 0.094 – 0.428 | 0.794 | E | n.s. |

| SR11 – $L_r$ | traditional regression | | | | | | | PIC regression | | | | | |
|---|---|---|---|---|---|---|---|---|---|---|---|---|---|
| | **n** | ***a*** | **95% CI$_a$** | ***b*$_{trad}$** | **95% CI$_b$** | **R** | **sim.** | **n** | ***b*$_{PIC}$** | **95% CI$_b$** | **R** | **sim.** | ***b*$_{trad}$ ≠ *b*$_{PIC}$** |
| **whole sample** | 137 | 3.503 | 2.778 – 4.515 | 0.362 | 0.331 – 0.390 | 0.896 | G | 136 | 0.352 | 0.326 – 0.378 | 0.899 | G | × |
| **fissipeds** | 130 | 2.692 | 2.158 – 3.436 | 0.397 | 0.368 – 0.423 | 0.910 | + | 129 | 0.344 | 0.318 – 0.370 | 0.900 | G | ✓ |
| **Family** | | | | | | | | | | | | | |
| Canidae | 17 | 2.019 | 0.816 –10.965 | 0.454 | 0.269 – 0.557 | 0.879 | G | 16 | 0.476 | 0.332 – 0.620 | 0.839 | G | × |
| Mustelidae | 32 | 3.649 | 2.378 – 5.600 | 0.325 | 0.270 – 0.379 | 0.893 | G | 31 | 0.318 | 0.266 – 0.370 | 0.897 | G | × |
| Procyonidae | 7 | 5.160 | 1.453 – 18.328 | 0.325 | 0.168 – 0.482 | 0.907 | B | 6 | 0.300 | 0.181 – 0.419 | 0.946 | B | × |
| Ursidae | 7 | 10.348 | 5.301 – 20.199 | 0.260 | 0.203 – 0.317 | 0.982 | E | 6 | 0.261 | 0.197 – 0.325 | 0.980 | E | × |
| Felidae | 26 | 8.534 | 5.789 – 12.579 | 0.288 | 0.246 – 0.329 | 0.940 | E | 25 | 0.337 | 0.282 – 0.392 | 0.922 | G | × |
| Herpestidae | 12 | 3.392 | 1.410 – 8.158 | 0.368 | 0.245 – 0.492 | 0.880 | B | 11 | 0.354 | 0.234 – 0.474 | 0.881 | B | × |
| Eupleridae | 5 | 8.373 | 1.676 – 41.831 | 0.269 | 0.048 – 0.491 | 0.894 | B | | | | | | |
| Viverridae | 14 | 8.179 | 4.764 – 14.044 | 0.262 | 0.195 – 0.329 | 0.914 | B | 13 | 0.291 | 0.156 – 0.426 | 0.682 | B | × |
| **Locomotor type** | | | | | | | | | | | | | |
| arboreal | 7 | 3.606 | 0.891 – 14.601 | 0.363 | 0.190 – 0.536 | 0.910 | B | 6 | 0.368 | 0.184 – 0.552 | 0.915 | B | × |
| semiarboreal | 10 | 9.473 | 6.558 – 13.683 | 0.250 | 0.202 – 0.298 | 0.972 | E | 9 | 0.226 | 0.133 – 0.319 | 0.872 | E | × |
| scansorial | 45 | 5.991 | 4.633 – 7.748 | 0.317 | 0.290 – 0.345 | 0.958 | G | 44 | 0.336 | 0.300 – 0.372 | 0.939 | G | × |
| terrestrial | 49 | 1.709 | 1.167 – 2.768 | 0.457 | 0.396 – 0.506 | 0.941 | + | 48 | 0.361 | 0.312 – 0.410 | 0.890 | G | ✓ |
| semifossorial | 7 | 3.434 | 0.908 – 12.985 | 0.342 | 0.175 – 0.510 | 0.905 | B | 6 | 0.362 | 0.184 – 0.540 | 0.918 | B | × |
| semiaquatic | 11 | 4.515 | 1.267 – 16.083 | 0.293 | 0.144 – 0.442 | 0.739 | B | 10 | 0.239 | 0.142 – 0.336 | 0.850 | B | × |
| aquatic | 8 | 2.372 | 0.424 – 13.268 | 0.347 | 0.203 – 0.491 | 0.909 | B | 7 | 0.338 | 0.192 – 0.484 | 0.911 | B | × |

| **SR12 – $d_{sr}$** | **traditional regression** | | | | | | | **PIC regression** | | | | | |
|---|---|---|---|---|---|---|---|---|---|---|---|---|---|
| | **n** | ***a*** | **95% CI$_a$** | **$b_{trad}$** | **95% CI$_b$** | **R** | **sim.** | **n** | **$b_{PIC}$** | **95% CI$_b$** | **R** | **sim.** | **$b_{trad} \neq b_{PIC}$** |
| **whole sample** | 137 | 0.282 | 0.244 – 0.327 | 0.335 | 0.318 – 0.352 | 0.961 | G | 136 | 0.358 | 0.331 – 0.385 | 0.894 | B | × |
| **fissipeds** | 130 | 0.270 | 0.233 – 0.316 | 0.341 | 0.323 – 0.357 | 0.956 | G | 129 | 0.357 | 0.330 – 0.384 | 0.902 | B | × |
| **Family** | | | | | | | | | | | | | |
| Canidae | 17 | 0.096 | 0.052 – 0.327 | 0.465 | 0.329 – 0.531 | 0.936 | B | 16 | 0.427 | 0.326 – 0.528 | 0.905 | B | × |
| Mustelidae | 32 | 0.291 | 0.202 – 0.421 | 0.326 | 0.279 – 0.373 | 0.922 | G | 31 | 0.342 | 0.277 – 0.407 | 0.863 | B | × |
| Procyonidae | 7 | 0.485 | 0.154 – 1.528 | 0.275 | 0.133 – 0.417 | 0.893 | B | 6 | 0.254 | 0.129 – 0.379 | 0.917 | B | × |
| Ursidae | 7 | 0.314 | 0.035 – 2.851 | 0.320 | 0.132 – 0.508 | 0.860 | B | ***6*** | ***0.328*** | ***0.127 – 0.529*** | ***0.870*** | ***B*** | n.s. |
| Felidae | 26 | 0.154 | 0.107 – 0.221 | 0.391 | 0.353 – 0.430 | 0.972 | E | 25 | 0.395 | 0.339 – 0.451 | 0.942 | E | × |
| Herpestidae | 12 | 0.339 | 0.150 – 0.770 | 0.321 | 0.206 – 0.436 | 0.862 | B | 11 | 0.367 | 0.248 – 0.486 | 0.891 | B | × |
| Eupleridae | 5 | 0.325 | 0.129 – 0.819 | 0.318 | 0.191 – 0.445 | 0.976 | B | | | | | | |
| Viverridae | 14 | 0.331 | 0.197 – 0.558 | 0.314 | 0.250 – 0.379 | 0.945 | B | 13 | 0.349 | 0.217 – 0.481 | 0.803 | B | × |
| **Locomotor type** | | | | | | | | | | | | | |
| arboreal | 7 | 0.219 | 0.068 – 0.705 | 0.361 | 0.216 – 0.506 | 0.937 | B | 6 | 0.362 | 0.208 – 0.516 | 0.939 | B | × |
| semiarboreal | 10 | 0.672 | 0.443 – 1.021 | 0.228 | 0.173 – 0.282 | 0.956 | – | 9 | 0.232 | 0.164 – 0.300 | 0.937 | – | × |
| scansorial | 45 | 0.247 | 0.196 – 0.312 | 0.346 | 0.321 – 0.371 | 0.971 | G | 44 | 0.367 | 0.327 – 0.407 | 0.935 | B | × |
| terrestrial | 49 | 0.248 | 0.195 – 0.343 | 0.358 | 0.320 – 0.387 | 0.959 | B | 48 | 0.342 | 0.299 – 0.385 | 0.903 | B | × |
| semifossorial | 7 | 0.255 | 0.098 – 0.662 | 0.345 | 0.225 – 0.465 | 0.953 | B | 6 | 0.362 | 0.220 – 0.504 | 0.949 | B | × |
| semiaquatic | 11 | 0.183 | 0.072 – 0.466 | 0.382 | 0.273 – 0.492 | 0.925 | B | 10 | 0.459 | 0.286 – 0.632 | 0.872 | B | × |
| aquatic | 8 | 0.020 | 0.001 – 0.483 | 0.545 | 0.277 – 0.813 | 0.871 | B | ***7*** | ***0.544*** | ***0.136 – 0.952*** | ***0.700*** | ***B*** | n.s. |

| **SR13 – $d_{tr}$** | traditional regression | | | | | | | PIC regression | | | | | |
|---|---|---|---|---|---|---|---|---|---|---|---|---|---|
| | **n** | ***a*** | **95% CI$_a$** | **$b_{trad}$** | **95% CI$_b$** | **R** | **sim.** | **n** | **$b_{PIC}$** | **95% CI$_b$** | **R** | **sim.** | **$b_{trad} \neq b_{PIC}$** |
| **whole sample** | 137 | 0.140 | 0.117 – 0.171 | 0.442 | 0.420 – 0.463 | 0.962 | + | 136 | 0.403 | 0.375 – 0.431 | 0.910 | + | ✓ |
| **fissipeds** | 130 | 0.160 | 0.136 – 0.198 | 0.425 | 0.400 – 0.444 | 0.953 | + | 129 | 0.389 | 0.361 – 0.417 | 0.914 | E | ✓ |
| **Family** | | | | | | | | | | | | | |
| Canidae | 17 | 0.134 | 0.081 – 0.348 | 0.465 | 0.364 – 0.519 | 0.947 | E | 16 | 0.478 | 0.388 – 0.570 | 0.938 | + | × |
| Mustelidae | 32 | 0.239 | 0.169 – 0.338 | 0.352 | 0.308 – 0.397 | 0.942 | B | 31 | 0.382 | 0.308 – 0.456 | 0.857 | B | × |
| Procyonidae | 7 | 0.320 | 0.077 – 1.335 | 0.355 | 0.178 – 0.531 | 0.901 | B | 6 | 0.347 | 0.163 – 0.531 | 0.904 | B | × |
| Ursidae | 7 | 0.301 | 0.038 – 2.387 | 0.357 | 0.181 – 0.534 | 0.903 | B | 6 | 0.317 | 0.167 – 0.467 | 0.925 | B | × |
| Felidae | 26 | 0.240 | 0.188 – 0.306 | 0.389 | 0.363 – 0.415 | 0.988 | E | 25 | 0.379 | 0.345 – 0.413 | 0.977 | E | × |
| Herpestidae | 12 | 0.336 | 0.112 – 1.002 | 0.318 | 0.165 – 0.472 | 0.729 | B | 11 | 0.310 | 0.158 – 0.462 | 0.728 | B | × |
| Eupleridae | 5 | 0.161 | 0.072 – 0.362 | 0.438 | 0.326 – 0.549 | 0.990 | B | | | | | | |
| Viverridae | 14 | 0.253 | 0.116 – 0.552 | 0.360 | 0.263 – 0.456 | 0.904 | B | 13 | 0.420 | 0.265 – 0.575 | 0.816 | B | × |
| **Locomotor type** | | | | | | | | | | | | | |
| arboreal | 7 | 0.245 | 0.066 – 0.909 | 0.383 | 0.220 – 0.545 | 0.929 | B | 6 | 0.386 | 0.216 – 0.556 | 0.935 | B | × |
| semiarboreal | 10 | 0.213 | 0.090 – 0.502 | 0.397 | 0.286 – 0.509 | 0.939 | B | 9 | 0.366 | 0.253 – 0.479 | 0.929 | B | × |
| scansorial | 45 | 0.203 | 0.162 – 0.256 | 0.401 | 0.376 – 0.425 | 0.980 | + | 44 | 0.387 | 0.353 – 0.421 | 0.958 | E | × |
| terrestrial | 49 | 0.127 | 0.093 – 0.195 | 0.458 | 0.406 – 0.494 | 0.948 | + | 48 | 0.375 | 0.320 – 0.430 | 0.865 | B | ✓ |
| semifossorial | 7 | 0.209 | 0.051 – 0.860 | 0.376 | 0.157 – 0.554 | 0.911 | B | 6 | 0.401 | 0.182 – 0.620 | 0.898 | B | × |
| semiaquatic | 11 | 0.385 | 0.185 – 0.800 | 0.294 | 0.208 – 0.380 | 0.922 | B | 10 | 0.320 | 0.228 – 0.412 | 0.928 | B | × |
| aquatic | 8 | 0.031 | 0.003 – 0.277 | 0.581 | 0.398 – 0.764 | 0.949 | + | 7 | 0.500 | 0.264 – 0.736 | 0.893 | B | × |

| SR14 – P | traditional regression | | | | | | | PIC regression | | | | | |
|---|---|---|---|---|---|---|---|---|---|---|---|---|---|
| | **n** | ***a*** | **95% CI*a*** | ***b*trad** | **95% CI*b*** | **R** | **sim.** | **n** | ***b*PIC** | **95% CI*b*** | **R** | **sim.** | ***b*trad ≠ *b*PIC** |
| **whole sample** | 137 | 0.101 | 0.085 – 0.122 | 0.426 | 0.404 – 0.446 | 0.956 | + | 136 | 0.412 | 0.380 – 0.444 | 0.892 | + | × |
| **fissipeds** | 130 | 0.088 | 0.074 – 0.108 | 0.444 | 0.421 – 0.464 | 0.952 | + | 129 | 0.414 | 0.381 – 0.447 | 0.888 | + | × |
| **Family** | | | | | | | | | | | | | |
| Canidae | 17 | 0.046 | 0.016 – 0.095 | 0.514 | 0.428 – 0.624 | 0.960 | + | 16 | 0.488 | 0.396 – 0.580 | 0.940 | + | × |
| Mustelidae | 32 | 0.161 | 0.110 – 0.236 | 0.350 | 0.302 – 0.399 | 0.929 | G | 31 | 0.347 | 0.276 – 0.418 | 0.838 | G | × |
| Procyonidae | 7 | 0.186 | 0.028 – 1.241 | 0.352 | 0.117 – 0.587 | 0.814 | B | *6* | ***0.325*** | ***0.095 – 0.555*** | ***0.820*** | ***B*** | n.s. |
| Ursidae | 7 | 0.147 | 0.016 – 1.360 | 0.399 | 0.210 – 0.589 | 0.911 | B | 6 | 0.357 | 0.159 – 0.555 | 0.894 | B | × |
| Felidae | 26 | 0.182 | 0.131 – 0.253 | 0.380 | 0.345 – 0.415 | 0.976 | + | 25 | 0.369 | 0.325 – 0.413 | 0.959 | G | × |
| Herpestidae | 12 | 0.164 | 0.075 – 0.361 | 0.374 | 0.264 – 0.484 | 0.908 | G | 11 | 0.353 | 0.241 – 0.465 | 0.897 | B | × |
| Eupleridae | 5 | 0.027 | 0.001 – 1.001 | 0.611 | 0.113 – 1.108 | 0.896 | B | | | | | | |
| Viverridae | 14 | 0.096 | 0.029 – 0.321 | 0.424 | 0.275 – 0.574 | 0.828 | G | 13 | 0.483 | 0.292 – 0.674 | 0.783 | G | × |
| **Locomotor type** | | | | | | | | | | | | | |
| arboreal | 7 | 0.026 | 0.001 – 0.455 | 0.588 | 0.233 – 0.943 | 0.851 | B | *6* | ***0.589*** | ***0.211 – 0.967*** | ***0.935*** | ***B*** | n.s. |
| semiarboreal | 10 | 0.063 | 0.022 – 0.180 | 0.496 | 0.360 – 0.633 | 0.941 | + | 9 | 0.475 | 0.274 – 0.676 | 0.929 | G | × |
| scansorial | 45 | 0.089 | 0.066 – 0.119 | 0.448 | 0.416 – 0.480 | 0.972 | + | 44 | 0.421 | 0.373 – 0.469 | 0.958 | + | × |
| terrestrial | 49 | 0.092 | 0.070 – 0.130 | 0.442 | 0.400 – 0.475 | 0.956 | + | 48 | 0.294 | 0.255 – 0.333 | 0.865 | G | ✓ |
| semifossorial | 7 | 0.280 | 0.110 – 0.709 | 0.292 | 0.175 – 0.409 | 0.937 | B | 6 | 0.313 | 0.178 – 0.448 | 0.898 | B | × |
| semiaquatic | 11 | 0.117 | 0.028 – 0.478 | 0.386 | 0.220 – 0.551 | 0.823 | B | *10* | ***0.330*** | ***0.100 – 0.560*** | ***0.928*** | ***B*** | n.s. |
| aquatic | 8 | 0.037 | 0.004 – 0.354 | 0.487 | 0.298 – 0.676 | 0.922 | G | 7 | 0.481 | 0.277 – 0.685 | 0.893 | G | × |

| **SR15 – RR** | **traditional regression** | | | | | | | **PIC regression** | | | | | |
|---|---|---|---|---|---|---|---|---|---|---|---|---|---|
| | **n** | ***a*** | **95% CI$_a$** | **$b_{trad}$** | **95% CI$_b$** | **R** | **sim.** | **n** | **$b_{PIC}$** | **95% CI$_b$** | **R** | **sim.** | **$b_{trad} \neq b_{PIC}$** |
| **whole sample** | 137 | 0.250 | 0.209 – 4.676 | -0.158 | -0.496 – -0.135 | 0.021 | – | 136 | 0.200 | 0.169 – 0.231 | 0.409 | + | n.s. |
| **fissipeds** | 130 | 0.258 | 0.196 – 0.322 | -0.169 | -0.196 – -0.134 | 0.214 | – | 129 | 0.198 | 0.167 – 0.229 | 0.422 | + | ✓ |
| **Family** | | | | | | | | | | | | | |
| Canidae | 17 | 0.005 | $1.27 \cdot 10^{-5}$ – 0.025 | 0.260 | 0.080 – 0.915 | 0.141 | E | 16 | -0.324 | -0.499 – -0.149 | 0.223 | – | n.s. |
| Mustelidae | 32 | 0.028 | 0.003 – 0.039 | 0.139 | 0.097 – 0.432 | 0.080 | E | 31 | 0.161 | 0.101 – 0.221 | 0.062 | E | n.s. |
| Procyonidae | 7 | 0.121 | 0.033 – 0.375 | -0.081 | -0.224 – 0.070 | 0.585 | B | 6 | -0.088 | -0.180 – 0.004 | 0.542 | B | n.s. |
| Ursidae | 7 | 0.008 | $1.43 \cdot 10^{-5}$ – 0.055 | 0.169 | 0.002 – 0.712 | 0.082 | E | 6 | 0.168 | -0.038 – 0.374 | 0.142 | B | n.s. |
| Felidae | 26 | 0.011 | 0.008 – 0.017 | 0.155 | 0.112 – 0.189 | 0.709 | E | 25 | 0.165 | 0.103 – 0.227 | 0.458 | E | × |
| Herpestidae | 12 | 0.449 | 0.155 – 46.774 | -0.259 | -0.922 – -0.116 | 0.177 | – | 11 | 0.243 | 0.070 – 0.416 | 0.096 | E | n.s. |
| Eupleridae | 5 | 0.028 | 0.005 – 0.107 | 0.093 | -0.113 – 0.348 | 0.735 | B | | | | | | |
| Viverridae | 14 | 0.026 | 0.019 – 0.046 | 0.108 | 0.038 – 0.147 | 0.543 | E | 13 | 0.108 | 0.045 – 0.171 | 0.396 | E | n.s. |
| **Locomotor type** | | | | | | | | | | | | | |
| arboreal | 7 | 0.009 | $5.40 \cdot 10^{-7}$ – 0.029 | 0.237 | 0.106 – 1.454 | 0.047 | E | 6 | -0.190 | -0.425 – 0.045 | 0.030 | B | n.s. |
| semiarboreal | 10 | 0.124 | 0.085 – 0.691 | -0.095 | -0.326 – -0.051 | 0.263 | – | 9 | 0.106 | 0.018 – 0.194 | 0.115 | E | n.s. |
| scansorial | 45 | 0.016 | 0.011 – 0.023 | 0.129 | 0.093 – 0.171 | 0.242 | E | 44 | 0.142 | 0.099 – 0.185 | 0.189 | E | n.s. |
| terrestrial | 49 | 0.247 | 0.163 – 0.328 | -0.164 | -0.201 – -0.110 | 0.528 | – | 48 | -0.165 | -0.214 – -0.116 | 0.073 | – | n.s. |
| semifossorial | 7 | 0.033 | 0.004 – 0.067 | 0.104 | 0.002 – 0.361 | 0.193 | E | 6 | 0.120 | -0.029 – 0.269 | 0.064 | B | n.s. |
| semiaquatic | 11 | 0.010 | 0.003 – 0.079 | 0.252 | 0.027 – 0.387 | 0.547 | E | 10 | 0.248 | 0.131 – 0.365 | 0.789 | + | n.s. |
| aquatic | 8 | 0.004 | $3.55 \cdot 10^{-6}$ – 0.205 | 0.260 | -0.061 – 0.832 | 0.613 | B | 7 | 0.266 | 0.239 – 0.293 | 0.276 | + | n.s. |

| **SR16 – $L_u$** | **traditional regression** | | | | | | | **PIC regression** | | | | | |
|---|---|---|---|---|---|---|---|---|---|---|---|---|---|
| | **n** | ***a*** | **95% $CI_a$** | **$b_{trad}$** | **95% $CI_b$** | **R** | **sim.** | **n** | **$b_{PIC}$** | **95% $CI_b$** | **R** | **sim.** | **$b_{trad} \neq b_{PIC}$** |
| **whole sample** | 136 | 3.841 | 3.046 – 4.922 | 0.358 | 0.327 – 0.385 | 0.897 | G | 135 | 0.347 | 0.321 – 0.373 | 0.897 | G | × |
| **fissipeds** | 129 | 2.926 | 2.411 – 3.644 | 0.394 | 0.367 – 0.418 | 0.916 | + | 128 | 0.340 | 0.314 – 0.366 | 0.902 | G | ✓ |
| **Family** | | | | | | | | | | | | | |
| Canidae | 16 | 2.170 | 0.856 – 12.076 | 0.452 | 0.258 – 0.556 | 0.884 | G | 15 | 0.474 | 0.326 – 0.622 | 0.841 | G | × |
| Mustelidae | 32 | 4.037 | 2.665 – 6.115 | 0.322 | 0.269 – 0.375 | 0.898 | G | 31 | 0.315 | 0.263 – 0.367 | 0.896 | G | × |
| Procyonidae | 7 | 5.692 | 1.569 – 20.645 | 0.318 | 0.159 – 0.478 | 0.900 | B | 6 | 0.293 | 0.174 – 0.412 | 0.945 | B | × |
| Ursidae | 7 | 10.831 | 6.606 – 17.759 | 0.262 | 0.220 – 0.304 | 0.990 | E | 6 | 0.263 | 0.216 – 0.310 | 0.990 | E | × |
| Felidae | 26 | 9.200 | 6.415 – 13.195 | 0.286 | 0.248 – 0.324 | 0.948 | E | 25 | 0.317 | 0.267 – 0.367 | 0.928 | G | × |
| Herpestidae | 12 | 3.710 | 1.629 – 8.452 | 0.363 | 0.247 – 0.478 | 0.892 | B | 11 | 0.334 | 0.230 – 0.438 | 0.900 | B | × |
| Eupleridae | 5 | 8.643 | 2.236 – 33.406 | 0.270 | 0.083 – 0.456 | 0.927 | B | | | | | | |
| Viverridae | 14 | 8.785 | 5.053 – 15.275 | 0.260 | 0.191 – 0.328 | 0.908 | E | 13 | 0.291 | 0.147 – 0.435 | 0.631 | B | × |
| **Locomotor type** | | | | | | | | | | | | | |
| arboreal | 7 | 3.573 | 0.863 – 14.800 | 0.371 | 0.195 – 0.547 | 0.911 | B | 6 | 0.377 | 0.190 – 0.564 | 0.917 | B | × |
| semiarboreal | 10 | 9.563 | 6.305 – 14.505 | 0.257 | 0.203 – 0.311 | 0.966 | E | 9 | 0.227 | 0.130 – 0.324 | 0.860 | E | × |
| scansorial | 45 | 6.234 | 4.849 – 8.013 | 0.319 | 0.292 – 0.346 | 0.961 | G | 44 | 0.332 | 0.299 – 0.365 | 0.945 | G | × |
| terrestrial | 48 | 1.893 | 1.332 – 3.071 | 0.451 | 0.392 – 0.497 | 0.944 | + | 47 | 0.358 | 0.309 – 0.407 | 0.888 | G | ✓ |
| semifossorial | 7 | 3.904 | 1.139 – 13.375 | 0.333 | 0.178 – 0.488 | 0.914 | B | 6 | 0.349 | 0.180 – 0.518 | 0.921 | B | × |
| semiaquatic | 11 | 5.026 | 1.558 – 16.210 | 0.290 | 0.153 – 0.427 | 0.779 | B | 10 | 0.240 | 0.145 – 0.335 | 0.856 | B | × |
| aquatic | 8 | 4.077 | 0.572 – 29.063 | 0.303 | 0.139 – 0.468 | 0.840 | B | 7 | 0.298 | 0.123 – 0.473 | 0.830 | B | × |

| SR17 – $d_{su}$ | traditional regression | | | | | | | PIC regression | | | | | |
|---|---|---|---|---|---|---|---|---|---|---|---|---|---|
| | **n** | ***a*** | **95% CI$_a$** | **$b_{trad}$** | **95% CI$_b$** | **R** | **sim.** | **n** | **$b_{PIC}$** | **95% CI$_b$** | **R** | **sim.** | **$b_{trad} \neq b_{PIC}$** |
| **whole sample** | 136 | 0.263 | 0.223 – 0.320 | 0.369 | 0.348 – 0.387 | 0.948 | E | 135 | 0.365 | 0.336 – 0.394 | 0.886 | E | × |
| **fissipeds** | 129 | 0.257 | 0.216 – 0.312 | 0.372 | 0.350 – 0.392 | 0.936 | E | 128 | 0.364 | 0.334 – 0.394 | 0.885 | E | × |
| **Family** | | | | | | | | | | | | | |
| Canidae | 16 | 0.057 | 0.018 – 0.291 | 0.505 | 0.325 – 0.626 | 0.915 | B | 15 | 0.433 | 0.308 – 0.558 | 0.865 | B | × |
| Mustelidae | 32 | 0.320 | 0.231 – 0.442 | 0.345 | 0.304 – 0.386 | 0.947 | B | 31 | 0.353 | 0.289 – 0.417 | 0.873 | B | × |
| Procyonidae | 7 | 0.167 | 0.011 – 2.592 | 0.457 | 0.117 – 0.796 | 0.763 | B | *6* | *0.456* | *0.075 – 0.837* | *0.741* | *B* | n.s. |
| Ursidae | *7* | *1.015* | *0.098 – 10.539* | *0.258* | *0.059 – 0.457* | *0.741* | *B* | *6* | *0.246* | *0.040 – 0.452* | *0.738* | *B* | n.s. |
| Felidae | 26 | 0.309 | 0.210 – 0.455 | 0.358 | 0.317 – 0.399 | 0.963 | B | 25 | 0.349 | 0.294 – 0.404 | 0.928 | B | × |
| Herpestidae | 12 | 0.265 | 0.101 – 0.697 | 0.363 | 0.228 – 0.499 | 0.848 | B | 11 | 0.357 | 0.215 – 0.499 | 0.831 | B | × |
| Eupleridae | 5 | 0.184 | 0.056 – 0.603 | 0.423 | 0.259 – 0.586 | 0.978 | B | | | | | | |
| Viverridae | 14 | 0.374 | 0.142 – 0.985 | 0.330 | 0.210 – 0.450 | 0.815 | B | *13* | *0.365* | *0.168 – 0.562* | *0.527* | *B* | n.s. |
| **Locomotor type** | | | | | | | | | | | | | |
| arboreal | 7 | 0.192 | 0.048 – 0.770 | 0.427 | 0.254 – 0.599 | 0.936 | B | 6 | 0.435 | 0.242 – 0.628 | 0.934 | B | × |
| semiarboreal | 10 | 0.555 | 0.320 – 0.964 | 0.295 | 0.223 – 0.367 | 0.954 | G | 9 | 0.275 | 0.173 – 0.377 | 0.896 | B | × |
| scansorial | 45 | 0.266 | 0.198 – 0.357 | 0.376 | 0.344 – 0.408 | 0.961 | E | 44 | 0.366 | 0.315 – 0.417 | 0.891 | B | × |
| terrestrial | 48 | 0.298 | 0.226 – 0.407 | 0.338 | 0.304 – 0.372 | 0.931 | G | 47 | 0.359 | 0.304 – 0.414 | 0.857 | B | × |
| semifossorial | 7 | 0.258 | 0.127 – 0.524 | 0.379 | 0.290 – 0.468 | 0.979 | B | 6 | 0.382 | 0.266 – 0.498 | 0.970 | B | × |
| semiaquatic | 11 | 0.319 | 0.166 – 0.614 | 0.344 | 0.267 – 0.420 | 0.955 | B | 10 | 0.345 | 0.236 – 0.454 | 0.912 | B | × |
| aquatic | 8 | 0.115 | 0.007 – 1.916 | 0.434 | 0.198 – 0.670 | 0.839 | B | *7* | *0.462* | *0.143 – 0.781* | *0.753* | *B* | n.s. |

| SR18 – $d_{tu}$ | traditional regression | | | | | | | PIC regression | | | | | |
|---|---|---|---|---|---|---|---|---|---|---|---|---|---|
| | **n** | ***a*** | **95% CI$_a$** | **$b_{trad}$** | **95% CI$_b$** | **R** | **sim.** | **n** | **$b_{PIC}$** | **95% CI$_b$** | **R** | **sim.** | **$b_{trad} \neq b_{PIC}$** |
| **whole sample** | 136 | 0.199 | 0.169 – 0.238 | 0.370 | 0.351 – 0.389 | 0.958 | E | 135 | 0.374 | 0.346 – 0.402 | 0.897 | E | × |
| **fissipeds** | 129 | 0.182 | 0.152 – 0.215 | 0.382 | 0.363 – 0.403 | 0.951 | E | 128 | 0.482 | 0.427 – 0.537 | 0.760 | + | ✓ |
| **Family** | | | | | | | | | | | | | |
| Canidae | 16 | 0.150 | 0.055 – 0.518 | 0.410 | 0.274 – 0.518 | 0.904 | B | 15 | 0.411 | 0.290 – 0.532 | 0.861 | B | × |
| Mustelidae | 32 | 0.247 | 0.166 – 0.367 | 0.334 | 0.283 – 0.384 | 0.914 | B | 31 | 0.368 | 0.294 – 0.442 | 0.844 | B | × |
| Procyonidae | 7 | 0.435 | 0.122 – 1.557 | 0.270 | 0.112 – 0.427 | 0.861 | B | *6* | ***0.281*** | ***0.090 – 0.472*** | ***0.837*** | ***B*** | n.s. |
| Ursidae | 7 | 0.173 | 0.010 – 3.127 | 0.395 | 0.148 – 0.641 | 0.839 | B | *6* | ***0.409*** | ***0.138 – 0.680*** | ***0.845*** | ***B*** | n.s. |
| Felidae | 26 | 0.119 | 0.068 – 0.208 | 0.419 | 0.360 – 0.479 | 0.942 | E | 25 | 0.388 | 0.314 – 0.462 | 0.892 | B | × |
| Herpestidae | 12 | 0.233 | 0.096 – 0.569 | 0.370 | 0.245 – 0.495 | 0.877 | B | 11 | 0.367 | 0.246 – 0.488 | 0.887 | B | × |
| Eupleridae | 5 | 0.225 | 0.154 – 0.328 | 0.363 | 0.310 – 0.415 | 0.997 | B | | | | | | |
| Viverridae | 14 | 0.323 | 0.166 – 0.629 | 0.320 | 0.237 – 0.402 | 0.912 | B | 13 | 0.282 | 0.152 – 0.412 | 0.689 | B | × |
| **Locomotor type** | | | | | | | | | | | | | |
| arboreal | 7 | 0.250 | 0.068 – 0.914 | 0.342 | 0.181 – 0.502 | 0.913 | B | 6 | 0.222 | 0.219 – 0.225 | 1.000 | – | × |
| semiarboreal | 10 | 0.353 | 0.194 – 0.644 | 0.310 | 0.232 – 0.388 | 0.951 | B | 9 | 0.287 | 0.182 – 0.392 | 0.900 | B | × |
| scansorial | 45 | 0.139 | 0.101 – 0.191 | 0.408 | 0.374 – 0.442 | 0.961 | E | 44 | 0.409 | 0.352 – 0.466 | 0.893 | E | × |
| terrestrial | 48 | 0.186 | 0.148 – 0.242 | 0.386 | 0.357 – 0.412 | 0.964 | E | 47 | 0.375 | 0.329 – 0.421 | 0.909 | B | × |
| semifossorial | 7 | 0.106 | 0.053 – 0.524 | 0.374 | 0.230 – 0.518 | 0.942 | B | 6 | 0.377 | 0.225 – 0.529 | 0.945 | B | × |
| semiaquatic | 11 | 0.325 | 0.105 – 1.007 | 0.307 | 0.174 – 0.439 | 0.820 | B | 10 | 0.480 | 0.213 – 0.747 | 0.691 | B | × |
| aquatic | 8 | 0.033 | 0.006 – 0.181 | 0.503 | 0.361 – 0.645 | 0.959 | E | 7 | 0.479 | 0.336 – 0.622 | 0.959 | E | × |

| **SR19 – O** | traditional regression | | | | | | | PIC regression | | | | | |
|---|---|---|---|---|---|---|---|---|---|---|---|---|---|
| | **n** | ***a*** | **95% CI$_a$** | **$b_{trad}$** | **95% CI$_b$** | **R** | **sim.** | **n** | **$b_{PIC}$** | **95% CI$_b$** | **R** | **sim.** | **$b_{trad} \neq b_{PIC}$** |
| **whole sample** | 136 | 0.721 | 0.633 – 0.831 | 0.368 | 0.351 – 0.384 | 0.976 | + | 135 | 0.369 | 0.349 – 0.389 | 0.950 | + | × |
| **fissipeds** | 129 | 0.674 | 0.588 – 0.779 | 0.377 | 0.359 – 0.393 | 0.974 | + | 128 | 0.370 | 0.350 – 0.390 | 0.949 | + | × |
| **Family** | | | | | | | | | | | | | |
| Canidae | 16 | 0.425 | 0.263 – 0.902 | 0.436 | 0.352 – 0.488 | 0.959 | + | 15 | 0.378 | 0.298 – 0.458 | 0.930 | G | × |
| Mustelidae | 32 | 0.641 | 0.486 – 0.846 | 0.372 | 0.337 – 0.407 | 0.967 | + | 31 | 0.369 | 0.323 – 0.415 | 0.942 | G | × |
| Procyonidae | 7 | 0.543 | 0.068 – 4.328 | 0.400 | 0.143 – 0.657 | 0.829 | B | *6* | *0.409* | *0.077 – 0.741* | *0.756* | *B* | n.s. |
| Ursidae | 7 | 1.978 | 0.319 – 12.275 | 0.269 | 0.113 – 0.424 | 0.864 | B | *6* | *0.276* | *0.097 – 0.455* | *0.853* | *B* | n.s. |
| Felidae | 26 | 0.776 | 0.615 – 0.977 | 0.370 | 0.345 – 0.394 | 0.987 | + | 25 | 0.362 | 0.327 – 0.397 | 0.974 | G | × |
| Herpestidae | 12 | 0.962 | 0.588 – 1.573 | 0.334 | 0.265 – 0.403 | 0.956 | G | 11 | 0.347 | 0.288 – 0.406 | 0.971 | G | × |
| Eupleridae | 5 | 0.612 | 0.229 – 1.632 | 0.402 | 0.267 – 0.537 | 0.983 | G | | | | | | |
| Viverridae | 14 | 0.626 | 0.350 – 1.122 | 0.384 | 0.312 – 0.456 | 0.954 | G | 13 | 0.405 | 0.323 – 0.487 | 0.948 | G | × |
| **Locomotor type** | | | | | | | | | | | | | |
| arboreal | 7 | 0.402 | 0.093 – 1.735 | 0.440 | 0.259 – 0.621 | 0.934 | B | 6 | 0.377 | 0.376 – 0.378 | 1.000 | + | × |
| semiarboreal | 10 | 0.560 | 0.311 – 1.008 | 0.401 | 0.325 – 0.478 | 0.972 | G | 9 | 0.378 | 0.245 – 0.511 | 0.906 | B | × |
| scansorial | 45 | 0.789 | 0.633 – 0.983 | 0.360 | 0.337 – 0.384 | 0.976 | + | 44 | 0.358 | 0.323 – 0.393 | 0.948 | G | × |
| terrestrial | 48 | 0.605 | 0.494 – 0.789 | 0.391 | 0.358 – 0.417 | 0.975 | + | 47 | 0.359 | 0.327 – 0.391 | 0.953 | G | × |
| semifossorial | 7 | 0.596 | 0.316 – 1.124 | 0.400 | 0.320 – 0.480 | 0.985 | G | 6 | 0.423 | 0.289 – 0.557 | 0.967 | G | × |
| semiaquatic | 11 | 0.640 | 0.281 – 1.459 | 0.375 | 0.278 – 0.471 | 0.940 | G | 10 | 0.419 | 0.295 – 0.543 | 0.923 | G | × |
| aquatic | 8 | 0.186 | 0.012 – 2.791 | 0.469 | 0.242 – 0.696 | 0.875 | B | 7 | 0.458 | 0.284 – 0.632 | 0.932 | G | × |

| SR20 – θ | traditional regression | | | | | | | PIC regression | | | | | |
|---|---|---|---|---|---|---|---|---|---|---|---|---|---|
| | n | *a* | 95% CI$_a$ | $b_{trad}$ | 95% CI$_b$ | R | sim. | n | $b_{PIC}$ | 95% CI$_b$ | R | sim. | $b_{trad} \neq b_{PIC}$ |
| **whole sample** | 136 | 0.078 | 0.008 – 0.090 | $9.70 \cdot 10^{-7}$ | $4.60 \cdot 10^{-7}$ – $3.56 \cdot 10^{-7}$ | 0.048 | – | 135 | $2.44 \cdot 10^{-6}$ | $2.02 \cdot 10^{-6}$ – $2.86 \cdot 10^{-6}$ | 0.047 | – | n.s. |
| **fissipeds** | 129 | 0.132 | 0.115 – 0.148 | $-1.52 \cdot 10^{-6}$ | $-2.06 \cdot 10^{-6}$ – $-9.73 \cdot 10^{-10}$ | 0.100 | – | 128 | $2.58 \cdot 10^{-6}$ | $2.13 \cdot 10^{-6}$ – $3.03 \cdot 10^{-6}$ | 0.018 | – | n.s. |
| **Family** | | | | | | | | | | | | | |
| Canidae | 16 | 0.019 | -0.059 – 0.042 | $3.63 \cdot 10^{-6}$ | $5.98 \cdot 10^{-7}$ – $1.22 \cdot 10^{-5}$ | 0.447 | – | 15 | $3.63 \cdot 10^{-6}$ | $1.99 \cdot 10^{-6}$ – $5.27 \cdot 10^{-6}$ | 0.623 | – | n.s. |
| Mustelidae | 32 | 0.110 | -0.041 – 0.141 | $1.17 \cdot 10^{-5}$ | $-1.51 \cdot 10^{-7}$ – $3.71 \cdot 10^{-5}$ | 0.064 | – | 31 | $-2.93 \cdot 10^{-5}$ | $-4.02 \cdot 10^{-5}$ – $-1.84 \cdot 10^{-5}$ | 0.081 | – | n.s. |
| Procyonidae | 7 | 0.025 | -0.047 – 0.096 | $1.95 \cdot 10^{-5}$ | $7.02 \cdot 10^{-6}$ – $3.15 \cdot 10^{-5}$ | 0.689 | – | 6 | $2.08 \cdot 10^{-5}$ | $-2.00 \cdot 10^{-8}$ – $4.16 \cdot 10^{-5}$ | 0.594 | – | n.s. |
| Ursidae | 7 | 0.033 | -0.020 – 0.085 | $2.52 \cdot 10^{-7}$ | $-2.84 \cdot 10^{-7}$ – $1.63 \cdot 10^{-6}$ | 0.241 | – | 6 | $2.71 \cdot 10^{-7}$ | $-6.21 \cdot 10^{-8}$ – $6.04 \cdot 10^{-7}$ | 0.154 | – | n.s. |
| Felidae | 26 | 0.059 | -0.022 – 0.082 | $1.12 \cdot 10^{-6}$ | $-4.11 \cdot 10^{-7}$ – $4.92 \cdot 10^{-6}$ | 0.073 | – | 25 | $-1.43 \cdot 10^{-6}$ | $-2.03 \cdot 10^{-6}$ – $-8.26 \cdot 10^{-7}$ | 0.006 | – | n.s. |
| Herpestidae | 12 | 0.027 | -0.020 – 0.071 | $4.82 \cdot 10^{-5}$ | $6.09 \cdot 10^{-6}$ – $6.51 \cdot 10^{-5}$ | 0.588 | – | 11 | $5.04 \cdot 10^{-5}$ | $1.92 \cdot 10^{-5}$ – $8.16 \cdot 10^{-5}$ | 0.497 | – | n.s. |
| Eupleridae | 5 | 0.085 | 0.018 – 0.152 | $2.40 \cdot 10^{-5}$ | $1.85 \cdot 10^{-4}$ – $1.28 \cdot 10^{-4}$ | 0.866 | – | | | | | | |
| Viverridae | 14 | 0.113 | 0.090 – 0.179 | $-7.14 \cdot 10^{-6}$ | $-2.80 \cdot 10^{-5}$ – $-3.26 \cdot 10^{-6}$ | 0.428 | – | 13 | $-5.48 \cdot 10^{-5}$ | $-8.94 \cdot 10^{-5}$ – $-2.02 \cdot 10^{-5}$ | 0.134 | – | n.s. |
| **Locomotor type** | | | | | | | | | | | | | |
| arboreal | 7 | 0.095 | 0.004 – 0.260 | $-7.49 \cdot 10^{-6}$ | $-6.38 \cdot 10^{-5}$ – $2.51 \cdot 10^{-5}$ | 0.462 | – | 6 | $-7.17 \cdot 10^{-6}$ | $-7.76 \cdot 10^{-6}$ – $-6.58 \cdot 10^{-6}$ | 0.998 | – | n.s. |
| semiarboreal | 10 | 0.202 | 0.108 – 0.378 | $-2.22 \cdot 10^{-5}$ | $-9.98 \cdot 10^{-5}$ – $3.73 \cdot 10^{-5}$ | 0.168 | – | 9 | $-1.74 \cdot 10^{-5}$ | $-3.18 \cdot 10^{-5}$ – $-3.00 \cdot 10^{-6}$ | 0.131 | – | n.s. |
| scansorial | 45 | 0.135 | 0.112 – 0.196 | $-1.04 \cdot 10^{-6}$ | $-2.87 \cdot 10^{-6}$ – $-2.12 \cdot 10^{-7}$ | 0.180 | – | 44 | $-1.33 \cdot 10^{-6}$ | $-1.74 \cdot 10^{-6}$ – $-9.20 \cdot 10^{-7}$ | 0.038 | – | n.s. |
| terrestrial | 48 | 0.111 | 0.060 – 0.167 | $-1.45 \cdot 10^{-6}$ | $-3.93 \cdot 10^{-6}$ – $6.65 \cdot 10^{-6}$ | 0.052 | – | 47 | $1.50 \cdot 10^{-5}$ | $-2.87 \cdot 10^{-5}$ – $5.87 \cdot 10^{-5}$ | 0.193 | – | n.s. |
| semifossorial | 7 | 0.047 | -0.013 – 0.088 | $1.75 \cdot 10^{-5}$ | $1.11 \cdot 10^{-5}$ – $2.56 \cdot 10^{-5}$ | 0.885 | – | 6 | $1.68 \cdot 10^{-5}$ | $3.50 \cdot 10^{-6}$ – $3.01 \cdot 10^{-5}$ | 0.772 | – | n.s. |
| semiaquatic | 11 | 0.106 | -0.064 – 0.160 | $8.20 \cdot 10^{-6}$ | $-4.47 \cdot 10^{-6}$ – $3.40 \cdot 10^{-5}$ | 0.070 | – | 10 | $-1.63 \cdot 10^{-5}$ | $-2.82 \cdot 10^{-5}$ – $-4.45 \cdot 10^{-6}$ | 0.315 | – | n.s. |
| aquatic | 8 | 0.035 | -0.318 – 0.215 | $5.07 \cdot 10^{-7}$ | $-7.40 \cdot 10^{-7}$ – $2.50 \cdot 10^{-6}$ | 0.314 | – | 7 | $5.75 \cdot 10^{-7}$ | $1.28 \cdot 10^{-7}$ – $1.02 \cdot 10^{-6}$ | 0.673 | – | n.s. |

| SR21 – α | traditional regression | | | | | | | PIC regression | | | | | |
|---|---|---|---|---|---|---|---|---|---|---|---|---|---|
| | n | *a* | 95% CI$_a$ | $b_{trad}$ | 95% CI$_b$ | R | sim. | n | $b_{PIC}$ | 95% CI$_b$ | R | sim. | $b_{trad} \neq b_{PIC}$ |
| **whole sample** | 136 | 0.020 | 0.012 – 0.035 | 0.314 | 0.253 – 0.366 | 0.589 | G | 135 | 0.388 | 0.329 – 0.447 | 0.451 | G | ✓ |
| **fissipeds** | 129 | 0.017 | 0.010 – 0.033 | 0.334 | 0.257 – 0.395 | 0.535 | G | 128 | 0.398 | 0.336 – 0.460 | 0.450 | + | ✓ |
| **Family** | | | | | | | | | | | | | |
| Canidae | 16 | 0.033 | 0.009 – 0.284 | 0.301 | 0.060 – 0.450 | 0.523 | B | *15* | *0.323* | *0.150 – 0.496* | *0.376* | *B* | n.s. |
| Mustelidae | *32* | *0.023* | *0.009 – 0.059* | *0.333* | *0.211 – 0.455* | *0.180* | *B* | *31* | *0.417* | *0.265 – 0.569* | *0.214* | *G* | n.s. |
| Procyonidae | 7 | 0.002 | 0.000 – 0.041 | 0.597 | 0.190 – 1.003 | 0.806 | B | *6* | *0.547* | *0.131 – 0.963* | *0.791* | *B* | n.s. |
| Ursidae | *7* | *0.005* | *0.000 – 0.974* | *0.385* | *-0.058 – 0.828* | *0.003* | *B* | *6* | *0.393* | *-0.095 – 0.881* | *0.022* | *B* | n.s. |
| Felidae | 26 | 0.016 | 0.008 – 0.031 | 0.309 | 0.235 – 0.383 | 0.822 | B | 25 | 0.369 | 0.269 – 0.469 | 0.766 | G | × |
| Herpestidae | *12* | *0.008* | *0.001 – 0.061* | *0.476* | *0.186 – 0.766* | *0.500* | *B* | *11* | *0.444* | *0.169 – 0.719* | *0.499* | *B* | n.s. |
| Eupleridae | *5* | *0.019* | *0.001 – 0.470* | *0.346* | *-0.095 – 0.788* | *0.720* | *B* | | | | | | |
| Viverridae | 14 | 0.003 | 0.000 – 0.025 | 0.490 | 0.241 – 0.739 | 0.588 | B | *13* | *0.484* | *0.188 – 0.780* | *0.266* | *B* | n.s. |
| **Locomotor type** | | | | | | | | | | | | | |
| arboreal | *7* | *0.003* | *0.000 – 0.222* | *0.475* | *-0.062 – 1.012* | *0.178* | *B* | *6* | *0.475* | *-0.108 – 1.058* | *0.148* | *B* | n.s. |
| semiarboreal | *10* | *0.005* | *0.000 – 0.079* | *0.474* | *0.124 – 0.824* | *0.422* | *B* | *9* | *0.429* | *0.100 – 0.758* | *0.395* | *B* | n.s. |
| scansorial | 45 | 0.024 | 0.016 – 0.038 | 0.268 | 0.221 – 0.314 | 0.822 | E | 44 | 0.326 | 0.252 – 0.400 | 0.674 | G | × |
| terrestrial | 48 | 0.023 | 0.007 – 0.104 | 0.334 | 0.157 – 0.470 | 0.556 | B | *47* | *0.404* | *0.289 – 0.519* | *0.281* | *G* | n.s. |
| semifossorial | *7* | *0.010* | *0.000 – 0.332* | *0.405* | *-0.032 – 0.842* | *0.347* | *B* | *6* | *0.397* | *-0.088 – 0.882* | *0.185* | *B* | n.s. |
| semiaquatic | *11* | *0.013* | *0.002 – 0.100* | *0.363* | *0.128 – 0.599* | *0.513* | *B* | *10* | *0.468* | *0.141 – 0.795* | *0.414* | *B* | n.s. |
| aquatic | *8* | *0.002* | *0.000 – 0.506* | *0.497* | *0.011 – 0.984* | *0.203* | *B* | *7* | *0.437* | *-0.022 – 0.896* | *0.000* | *B* | n.s. |

| **SR22 – UR** | **traditional regression** | | | | | | | **PIC regression** | | | | | |
|---|---|---|---|---|---|---|---|---|---|---|---|---|---|
| | **n** | ***a*** | **95% CI$_a$** | **$b_{trad}$** | **95% CI$_b$** | **R** | **sim.** | **n** | **$b_{PIC}$** | **95% CI$_b$** | **R** | **sim.** | **$b_{trad} \neq b_{PIC}$** |
| **whole sample** | *136* | *0.014* | *$4.28 \cdot 10^{-4}$ – 0.019* | *0.198* | *0.166 – 0.605* | *0.141* | *+* | *135* | *0.188* | *0.156 – 0.220* | *0.142* | *+* | n.s. |
| **fissipeds** | *129* | *0.408* | *0.300 – 15.740* | *-0.389* | *-0.634 – -0.166* | *0.067* | *–* | *128* | *0.188* | *0.155 – 0.221* | *0.083* | *+* | n.s. |
| **Family** | | | | | | | | | | | | | |
| Canidae | *16* | *0.003* | *$3.17 \cdot 10^{-6}$ – 0.019* | *0.302* | *0.077 – 1.045* | *0.212* | *E* | *15* | *-0.380* | *-0.597 – -0.163* | *0.167* | *–* | n.s. |
| Mustelidae | *32* | *0.032* | *0.004 – 0.052* | *0.139* | *0.084 – 0.407* | *0.272* | *E* | *31* | *0.143* | *0.090 – 0.196* | *0.175* | *E* | n.s. |
| Procyonidae | *7* | *0.005* | *$1.06 \cdot 10^{-8}$ – 0.026* | *0.358* | *0.159 – 1.907* | *0.199* | *+* | *6* | *0.340* | *-0.074 – 0.754* | *0.201* | *B* | n.s. |
| Ursidae | *7* | *0.742* | *0.051 – 273.527* | *-0.181* | *-0.688 – 0.062* | *0.397* | *G* | *6* | *-0.179* | *-0.377 – 0.019* | *0.457* | *B* | n.s. |
| Felidae | 26 | 0.015 | 0.009 – 0.024 | 0.162 | 0.108 – 0.216 | 0.458 | E | *25* | *0.198* | *0.115 – 0.281* | *0.154* | *E* | n.s. |
| Herpestidae | *12* | *0.403* | *0.123 – 41.976* | *-0.243* | *-0.889 – -0.073* | *0.067* | *–* | *11* | *-0.234* | *-0.401 – -0.067* | *0.043* | *–* | n.s. |
| Eupleridae | *5* | *0.012* | *$4.22 \cdot 10^{-5}$ – 0.055* | *0.238* | *-0.008 – 1.038* | *0.693* | *B* | | | | | | |
| Viverridae | *14* | *0.024* | *0.002 – 0.043* | *0.143* | *0.068 – 0.484* | *0.229* | *E* | *13* | *0.130* | *0.054 – 0.206* | *0.405* | *E* | n.s. |
| **Locomotor type** | | | | | | | | | | | | | |
| arboreal | *7* | *0.018* | *$1.75 \cdot 10^{-5}$ – 0.037* | *0.194* | *0.110 – 1.069* | *0.319* | *E* | 6 | 0.179 | 0.157 – 0.201 | 0.995 | + | n.s. |
| semiarboreal | *10* | *0.034* | *0.003 – 0.052* | *0.109* | *0.044 – 0.418* | *0.312* | *E* | *9* | *0.106* | *0.033 – 0.179* | *0.578* | *E* | n.s. |
| scansorial | 45 | 0.018 | 0.013 – 0.027 | 0.149 | 0.105 – 0.186 | 0.361 | E | *44* | *0.280* | *0.194 – 0.366* | *0.082* | *+* | n.s. |
| terrestrial | 48 | 0.416 | 0.232 – 0.623 | -0.233 | -0.289 – -0.156 | 0.477 | – | *47* | *-0.189* | *-0.245 – -0.133* | *0.050* | *–* | n.s. |
| semifossorial | *7* | *0.028* | *0.003 – 0.066* | *0.156* | *0.053 – 0.433* | *0.431* | *E* | *6* | *0.132* | *-0.029 – 0.293* | *0.175* | *B* | n.s. |
| semiaquatic | *11* | *0.019* | *0.006 – 0.098* | *0.199* | *0.014 – 0.317* | *0.516* | *E* | 10 | 0.113 | 0.056 – 0.170 | 0.754 | E | n.s. |
| aquatic | *8* | *0.013* | *$2.70 \cdot 10^{-5}$ – 0.313* | *0.197* | *-0.076 – 0.700* | *0.552* | *B* | *7* | *0.233* | *0.011 – 0.455* | *0.424* | *E* | n.s. |

| SR23 – IFA | traditional regression | | | | | | | PIC regression | | | | | |
|---|---|---|---|---|---|---|---|---|---|---|---|---|---|
| | n | *a* | 95% CI$_a$ | $b_{trad}$ | 95% CI$_b$ | R | sim. | n | $b_{PIC}$ | 95% CI$_b$ | R | sim. | $b_{trad} \neq b_{PIC}$ |
| **whole sample** | 136 | 0.062 | 0.053 – 0.075 | 0.138 | 0.117 – 0.156 | 0.275 | + | 135 | 0.161 | 0.135 – 0.187 | 0.332 | + | × |
| **fissipeds** | *129* | *0.064* | *0.006 – 0.079* | *0.134* | *0.109 – 0.418* | *0.041* | *+* | 128 | 0.161 | 0.134 – 0.188 | 0.278 | + | n.s. |
| **Family** | | | | | | | | | | | | | |
| Canidae | *16* | *0.020* | *$5.22 \cdot 10^{-5}$ – 0.040* | *0.239* | *0.149 – 0.893* | *0.079* | *+* | *15* | *-0.285* | *-0.449 – -0.121* | *0.072* | *–* | n.s. |
| Mustelidae | 32 | 0.070 | 0.049 – 0.109 | 0.155 | 0.103 – 0.198 | 0.456 | + | 31 | 0.149 | 0.099 – 0.199 | 0.441 | + | × |
| Procyonidae | *7* | *0.025* | *$3.36 \cdot 10^{-6}$ – 0.374* | *0.249* | *-0.064 – 1.310* | *0.216* | *B* | *6* | *0.249* | *-0.054 – 0.552* | *0.197* | *B* | n.s. |
| Ursidae | *7* | *1.246* | *0.007 – 119.95* | *-0.157* | *-0.540 – 0.292* | *0.180* | *B* | *6* | *0.434* | *0.008 – 0.860* | *0.614* | *+* | n.s. |
| Felidae | 26 | 0.055 | 0.043 – 0.073 | 0.130 | 0.099 – 0.154 | 0.726 | + | 25 | 0.127 | 0.080 – 0.174 | 0.463 | + | × |
| Herpestidae | *12* | *0.838* | *0.566 – 35.400* | *-0.194* | *-0.724 – -0.138* | *0.021* | *–* | *11* | *0.177* | *0.053 – 0.301* | *0.209* | *+* | n.s. |
| Eupleridae | 5 | 0.059 | 0.011 – 0.212 | 0.158 | -0.038 – 0.400 | 0.919 | B | | | | | | |
| Viverridae | 14 | 0.046 | 0.026 – 0.089 | 0.179 | 0.088 – 0.251 | 0.731 | + | 13 | 0.180 | 0.085 – 0.275 | 0.565 | + | × |
| **Locomotor type** | | | | | | | | | | | | | |
| arboreal | *7* | *0.037* | *$3.77 \cdot 10^{-5}$ – 0.264* | *0.206* | *-0.052 – 1.067* | *0.350* | *B* | 6 | -0.056 | -0.066 – -0.046 | 0.990 | – | n.s. |
| semiarboreal | 10 | 0.049 | 0.030 – 0.091 | 0.168 | 0.080 – 0.231 | 0.849 | + | 9 | 0.184 | 0.092 – 0.276 | 0.802 | + | × |
| scansorial | 45 | 0.078 | 0.066 – 0.097 | 0.094 | 0.072 – 0.113 | 0.479 | + | *44* | *0.183* | *0.128 – 0.238* | *0.241* | *+* | n.s. |
| terrestrial | 48 | 0.535 | 0.379 – 0.686 | -0.123 | -0.155 – -0.079 | 0.355 | – | *47* | *0.155* | *0.109 – 0.201* | *0.156* | *+* | n.s. |
| semifossorial | *7* | *0.074* | *0.015 – 0.170* | *0.158* | *0.052 – 0.358* | *0.562* | *+* | *6* | *0.161* | *-0.016 – 0.338* | *0.471* | *B* | n.s. |
| semiaquatic | 11 | 0.053 | 0.024 – 0.213 | 0.189 | 0.034 – 0.275 | 0.673 | + | 10 | 0.249 | 0.114 – 0.384 | 0.707 | + | × |
| aquatic | 8 | 0.027 | $1.72 \cdot 10^{-4}$ – 0.062 | 0.210 | 0.134 – 0.622 | 0.737 | + | 7 | 0.207 | 0.073 – 0.341 | 0.785 | + | × |

| SR24 – $L_m$ | traditional regression | | | | | | | PIC regression | | | | | |
|---|---|---|---|---|---|---|---|---|---|---|---|---|---|
| | **n** | ***a*** | **95% CI$_a$** | ***b*$_{trad}$** | **95% CI$_b$** | **R** | **sim.** | **n** | ***b*$_{PIC}$** | **95% CI$_b$** | **R** | **sim.** | **$b_{trad} \neq b_{PIC}$** |
| **whole sample** | 136 | 1.498 | 1.174 – 1.972 | 0.351 | 0.315 – 0.380 | 0.873 | G | 135 | 0.347 | 0.319 – 0.375 | 0.885 | G | × |
| **fissipeds** | 129 | 1.171 | 0.907 – 1.567 | 0.383 | 0.347 – 0.415 | 0.884 | + | 128 | 0.348 | 0.320 – 0.376 | 0.887 | G | × |
| **Family** | | | | | | | | | | | | | |
| Canidae | 16 | 0.915 | 0.440 – 4.034 | 0.449 | 0.285 – 0.532 | 0.926 | G | 15 | 0.458 | 0.344 – 0.572 | 0.902 | + | × |
| Mustelidae | 32 | 2.017 | 1.281 – 3.176 | 0.294 | 0.236 – 0.352 | 0.849 | B | 31 | 0.299 | 0.240 – 0.358 | 0.847 | B | × |
| Procyonidae | 7 | 2.983 | 0.809 – 10.995 | 0.251 | 0.089 – 0.412 | 0.828 | B | 6 | 0.225 | 0.095 – 0.355 | 0.883 | B | × |
| Ursidae | 7 | 0.998 | 0.156 – 6.389 | 0.351 | 0.193 – 0.509 | 0.920 | B | 6 | 0.345 | 0.178 – 0.512 | 0.921 | B | × |
| Felidae | 26 | 2.709 | 1.964 – 3.738 | 0.310 | 0.275 – 0.344 | 0.965 | G | 25 | 0.336 | 0.292 – 0.380 | 0.951 | G | × |
| Herpestidae | 12 | 1.369 | 0.721 – 2.599 | 0.373 | 0.283 – 0.463 | 0.940 | G | 11 | 0.354 | 0.263 – 0.445 | 0.934 | G | × |
| Eupleridae | 5 | 4.309 | 1.684 – 11.025 | 0.217 | 0.088 – 0.346 | 0.946 | B | | | | | | |
| Viverridae | 14 | 1.429 | 0.695 – 2.939 | 0.343 | 0.254 – 0.433 | 0.911 | G | 13 | 0.336 | 0.212 – 0.460 | 0.815 | B | × |
| **Locomotor type** | | | | | | | | | | | | | |
| arboreal | *7* | *1.114* | *0.093 – 13.348* | *0.369* | *0.062 – 0.677* | *0.689* | *B* | *6* | *0.379* | *0.049 – 0.709* | *0.714* | *B* | n.s. |
| semiarboreal | 10 | 2.040 | 0.656 – 6.344 | 0.315 | 0.167 – 0.462 | 0.818 | B | 9 | 0.276 | 0.146 – 0.406 | 0.826 | B | × |
| scansorial | 45 | 2.240 | 1.535 – 3.268 | 0.314 | 0.273 – 0.354 | 0.905 | G | 44 | 0.318 | 0.282 – 0.354 | 0.929 | G | × |
| terrestrial | 48 | 0.788 | 0.507 – 1.360 | 0.497 | 0.377 – 0.505 | 0.934 | + | 47 | 0.355 | 0.310 – 0.400 | 0.904 | G | ✓ |
| semifossorial | 7 | 1.646 | 0.455 – 5.947 | 0.301 | 0.140 – 0.463 | 0.884 | B | 6 | 0.307 | 0.143 – 0.471 | 0.903 | B | × |
| semiaquatic | 11 | 2.832 | 1.128 – 7.113 | 0.257 | 0.149 – 0.365 | 0.831 | B | 10 | 0.281 | 0.166 – 0.396 | 0.847 | B | × |
| aquatic | 8 | 0.153 | 0.029 – 0.813 | 0.491 | 0.352 – 0.631 | 0.959 | + | 7 | 0.471 | 0.286 – 0.656 | 0.927 | G | × |

| SR25 – $d_{sm}$ | traditional regression | | | | | | | PIC regression | | | | | |
|---|---|---|---|---|---|---|---|---|---|---|---|---|---|
| | n | *a* | 95% $CI_a$ | $b_{trad}$ | 95% $CI_b$ | R | sim. | n | $b_{PIC}$ | 95% $CI_b$ | R | sim. | $b_{trad} \neq b_{PIC}$ |
| **whole sample** | 136 | 0.192 | 0.167 – 0.225 | 0.337 | 0.320 – 0.352 | 0.963 | G | 135 | 0.337 | 0.313 – 0.361 | 0.907 | G | × |
| **fissipeds** | 129 | 0.178 | 0.153 – 0.214 | 0.346 | 0.326 – 0.363 | 0.957 | G | 128 | 0.335 | 0.310 – 0.360 | 0.909 | G | × |
| **Family** | | | | | | | | | | | | | |
| Canidae | 16 | 0.051 | 0.021 – 0.126 | 0.492 | 0.385 – 0.588 | 0.939 | + | 15 | 0.433 | 0.335 – 0.531 | 0.920 | E | × |
| Mustelidae | 32 | 0.212 | 0.152 – 0.294 | 0.320 | 0.278 – 0.362 | 0.937 | G | 31 | 0.311 | 0.259 – 0.363 | 0.894 | G | × |
| Procyonidae | 7 | 0.183 | 0.025 – 1.354 | 0.336 | 0.088 – 0.583 | 0.767 | B | *6* | *0.316* | *0.057 – 0.575* | *0.751* | *B* | n.s. |
| Ursidae | 7 | 0.884 | 0.307 – 2.549 | 0.197 | 0.107 – 0.287 | 0.918 | – | 6 | 0.176 | 0.086 – 0.266 | 0.912 | – | × |
| Felidae | 26 | 0.209 | 0.171 – 0.257 | 0.336 | 0.314 – 0.358 | 0.988 | G | 25 | 0.323 | 0.291 – 0.355 | 0.971 | G | × |
| Herpestidae | 12 | 0.328 | 0.174 – 0.618 | 0.281 | 0.192 – 0.370 | 0.893 | G | 11 | 0.281 | 0.193 – 0.369 | 0.898 | G | × |
| Eupleridae | 5 | 0.084 | 0.016 – 0.430 | 0.429 | 0.203 – 0.654 | 0.958 | B | | | | | | |
| Viverridae | 14 | 0.197 | 0.092 – 0.420 | 0.326 | 0.232 – 0.420 | 0.889 | B | 13 | 0.362 | 0.234 – 0.490 | 0.830 | B | × |
| **Locomotor type** | | | | | | | | | | | | | |
| arboreal | 7 | 0.114 | 0.020 – 0.648 | 0.387 | 0.173 – 0.602 | 0.876 | B | 6 | 0.383 | 0.155 – 0.611 | 0.878 | B | × |
| semiarboreal | 10 | 0.174 | 0.075 – 0.405 | 0.352 | 0.242 – 0.462 | 0.924 | B | 9 | 0.350 | 0.178 – 0.522 | 0.810 | B | × |
| scansorial | 45 | 0.174 | 0.139 – 0.217 | 0.349 | 0.326 – 0.373 | 0.975 | G | 44 | 0.335 | 0.299 – 0.371 | 0.937 | G | × |
| terrestrial | 48 | 0.182 | 0.142 – 0.265 | 0.346 | 0.303 – 0.376 | 0.948 | B | 47 | 0.324 | 0.283 – 0.365 | 0.902 | G | × |
| semifossorial | 7 | 0.214 | 0.064 – 0.719 | 0.321 | 0.168 – 0.473 | 0.911 | B | 6 | 0.325 | 0.160 – 0.490 | 0.912 | B | × |
| semiaquatic | 11 | 0.160 | 0.068 – 0.378 | 0.359 | 0.259 – 0.460 | 0.929 | B | 10 | 0.364 | 0.227 – 0.501 | 0.871 | B | × |
| aquatic | 8 | 0.026 | 0.014 – 0.048 | 0.488 | 0.438 – 0.539 | 0.995 | + | 7 | 0.489 | 0.437 – 0.541 | 0.995 | + | × |

| **SR26 – $d_{tm}$** | traditional regression | | | | | | | PIC regression | | | | | |
|---|---|---|---|---|---|---|---|---|---|---|---|---|---|
| | **n** | ***a*** | **95% CI$_a$** | **$b_{trad}$** | **95% CI$_b$** | **R** | **sim.** | **n** | **$b_{PIC}$** | **95% CI$_b$** | **R** | **sim.** | **$b_{trad} \neq b_{PIC}$** |
| **whole sample** | 136 | 0.200 | 0.176 – 0.232 | 0.339 | 0.321 – 0.354 | 0.964 | G | 135 | 0.364 | 0.341 – 0.387 | 0.932 | E | ✓ |
| **fissipeds** | 129 | 0.180 | 0.155 – 0.209 | 0.353 | 0.334 – 0.370 | 0.962 | nei. | 128 | 0.362 | 0.338 – 0.386 | 0.930 | E | × |
| **Family** | | | | | | | | | | | | | |
| Canidae | 16 | 0.091 | 0.051 – 0.346 | 0.436 | 0.295 – 0.496 | 0.919 | B | 15 | 0.406 | 0.314 – 0.498 | 0.919 | B | × |
| Mustelidae | 32 | 0.191 | 0.144 – 0.254 | 0.341 | 0.305 – 0.377 | 0.959 | B | 31 | 0.375 | 0.326 – 0.424 | 0.937 | B | × |
| Procyonidae | *7* | *0.471* | *0.115 – 1.927* | *0.223* | *0.048 – 0.397* | *0.733* | *B* | *6* | *0.221* | *0.022 – 0.420* | *0.687* | *B* | n.s. |
| Ursidae | 7 | 0.593 | 0.205 – 1.714 | 0.236 | 0.146 – 0.327 | 0.943 | – | 6 | 0.237 | 0.140 – 0.334 | 0.944 | G | × |
| Felidae | 26 | 0.179 | 0.127 – 0.253 | 0.358 | 0.322 – 0.395 | 0.970 | B | 25 | 0.364 | 0.318 – 0.410 | 0.953 | B | × |
| Herpestidae | 12 | 0.347 | 0.191 – 0.629 | 0.280 | 0.197 – 0.364 | 0.906 | G | 11 | 0.263 | 0.176 – 0.350 | 0.885 | G | × |
| Eupleridae | 5 | 0.129 | 0.030 – 0.559 | 0.391 | 0.189 – 0.594 | 0.959 | B | | | | | | |
| Viverridae | 14 | 0.217 | 0.120 – 0.395 | 0.318 | 0.244 – 0.391 | 0.929 | B | 13 | 0.352 | 0.177 – 0.527 | 0.624 | B | × |
| **Locomotor type** | | | | | | | | | | | | | |
| arboreal | 7 | 0.159 | 0.045 – 0.555 | 0.355 | 0.200 – 0.509 | 0.925 | B | 6 | 0.360 | 0.199 – 0.521 | 0.876 | B | × |
| semiarboreal | 10 | 0.156 | 0.075 – 0.323 | 0.371 | 0.276 – 0.466 | 0.949 | B | 9 | 0.409 | 0.249 – 0.569 | 0.924 | B | × |
| scansorial | 45 | 0.181 | 0.142 – 0.232 | 0.351 | 0.324 – 0.377 | 0.969 | B | 44 | 0.369 | 0.328 – 0.410 | 0.975 | B | × |
| terrestrial | 48 | 0.179 | 0.139 – 0.238 | 0.359 | 0.324 – 0.391 | 0.963 | B | 47 | 0.355 | 0.317 – 0.393 | 0.948 | B | × |
| semifossorial | 7 | 0.153 | 0.048 – 0.489 | 0.372 | 0.226 – 0.518 | 0.940 | B | 6 | 0.375 | 0.210 – 0.540 | 0.911 | B | × |
| semiaquatic | 11 | 0.206 | 0.076 – 0.557 | 0.327 | 0.210 – 0.444 | 0.881 | B | 10 | 0.366 | 0.212 – 0.520 | 0.929 | B | × |
| aquatic | 8 | 0.098 | 0.023 – 0.421 | 0.382 | 0.260 – 0.505 | 0.947 | B | 7 | 0.382 | 0.259 – 0.505 | 0.995 | B | × |

| SR27 – MR | traditional regression | | | | | | | PIC regression | | | | | |
|---|---|---|---|---|---|---|---|---|---|---|---|---|---|
| | n | *a* | 95% CI$_a$ | $b_{trad}$ | 95% CI$_b$ | R | sim. | n | $b_{PIC}$ | 95% CI$_b$ | R | sim. | $b_{trad} \neq b_{PIC}$ |
| **whole sample** | 136 | 0.031 | 0.002 – 0.039 | 0.150 | 0.127 – 0.459 | 0.122 | + | 135 | 0.168 | 0.139 – 0.197 | 0.043 | + | n.s. |
| **fissipeds** | 129 | 0.427 | 0.352 – 7.313 | -0.159 | -0.494 – -0.133 | 0.050 | – | 128 | 0.172 | 0.142 – 0.202 | 0.004 | + | n.s. |
| **Family** | | | | | | | | | | | | | |
| Canidae | 16 | 0.012 | $8.81 \cdot 10^{-5}$ – 0.028 | 0.219 | 0.109 – 0.757 | 0.226 | E | 15 | -0.231 | -0.364 – -0.098 | 0.039 | – | n.s. |
| Mustelidae | 32 | 0.038 | 0.022 – 0.065 | 0.157 | 0.095 – 0.224 | 0.315 | E | 31 | 0.167 | 0.105 – 0.229 | 0.139 | E | n.s. |
| Procyonidae | 7 | 0.022 | $1.21 \cdot 10^{-4}$ – 0.057 | 0.214 | 0.093 – 0.833 | 0.284 | E | 6 | 0.238 | -0.049 – 0.525 | 0.228 | B | n.s. |
| Ursidae | 7 | 1.315 | 0.225 – 4.209 | -0.188 | -0.290 – -0.035 | 0.773 | – | 6 | -0.187 | -0.337 – -0.037 | 0.765 | – | × |
| Felidae | 26 | 0.036 | 0.005 – 0.055 | 0.109 | 0.064 – 0.331 | 0.302 | E | 25 | -0.137 | -0.195 – -0.079 | 0.048 | – | n.s. |
| Herpestidae | 12 | 0.496 | 0.288 – 11.350 | -0.195 | -0.634 – -0.112 | 0.503 | – | 11 | -0.188 | -0.310 – -0.066 | 0.422 | – | n.s. |
| Eupleridae | 5 | 0.016 | $1.42 \cdot 10^{-4}$ – 0.471 | 0.243 | -0.279 – 0.902 | 0.851 | B | | | | | | |
| Viverridae | 14 | 0.384 | -0.715 – 0.975 | -0.145 | -0.544 – -0.060 | 0.143 | – | 13 | -0.176 | -0.284 – -0.068 | 0.260 | – | n.s. |
| **Locomotor type** | | | | | | | | | | | | | |
| arboreal | 7 | 0.019 | $4.10 \cdot 10^{-4}$ – 0.460 | 0.227 | -0.179 – 0.689 | 0.379 | B | 6 | -0.203 | -0.453 – 0.047 | 0.114 | B | n.s. |
| semiarboreal | 10 | 0.017 | $1.99 \cdot 10^{-4}$ – 0.087 | 0.245 | 0.022 – 0.855 | 0.278 | E | 9 | 0.269 | 0.063 – 0.475 | 0.404 | E | n.s. |
| scansorial | 45 | 0.033 | 0.026 – 0.042 | 0.130 | 0.102 – 0.156 | 0.428 | E | 44 | 0.142 | 0.099 – 0.185 | 0.130 | E | n.s. |
| terrestrial | 48 | 0.393 | 0.272 – 0.527 | -0.165 | -0.205 – -0.116 | 0.539 | – | 47 | -0.175 | -0.226 – -0.124 | 0.164 | G | n.s. |
| semifossorial | 7 | 0.069 | 0.010 – 0.130 | 0.101 | 0.021 – 0.351 | 0.263 | E | 6 | 0.118 | -0.026 – 0.262 | 0.163 | B | n.s. |
| semiaquatic | 11 | 0.038 | 0.028 – 0.073 | 0.148 | 0.070 – 0.187 | 0.811 | E | 10 | 0.154 | 0.055 – 0.253 | 0.546 | E | n.s. |
| aquatic | 8 | 0.041 | $1.73 \cdot 10^{-4}$ – 0.226 | 0.118 | -0.029 – 0.571 | 0.122 | B | 7 | 0.148 | $4.25 \cdot 10^{-4}$ – 0.296 | 0.320 | E | n.s. |

| SR28 – $\%_{prox}$ | traditional regression | | | | | | | PIC regression | | | | | |
|---|---|---|---|---|---|---|---|---|---|---|---|---|---|
| | n | $a$ | 95% CI$_a$ | $b_{trad}$ | 95% CI$_b$ | R | sim. | n | $b_{PIC}$ | 95% CI$_b$ | R | sim. | $b_{trad} \neq b_{PIC}$ |
| **whole sample** | 137 | 16.881 | 15.596 – 18.365 | 0.058 | 0.048 – 0.067 | 0.416 | + | 136 | 0.066 | 0.055 – 0.077 | 0.240 | + | × |
| **fissipeds** | *130* | *17.894* | *16.368 – 19.789* | *0.050* | *0.039 – 0.060* | *0.172* | *+* | 129 | 0.061 | 0.051 – 0.071 | 0.272 | + | n.s. |
| **Family** | | | | | | | | | | | | | |
| Canidae | *17* | *11.899* | *1.718 – 17.783* | *0.090* | *0.043 – 0.304* | *0.398* | *+* | *16* | *-0.112* | *-0.174 – -0.050* | *0.024* | *–* | n.s. |
| Mustelidae | 32 | 17.535 | 15.276 – 20.941 | 0.062 | 0.041 – 0.078 | 0.529 | + | 31 | 0.049 | 0.032 – 0.066 | 0.409 | + | × |
| Procyonidae | *7* | *7.280* | *0.619 – 29.174* | *0.154* | *-0.009 – 0.474* | *0.416* | *B* | *6* | *0.151* | *-0.023 – 0.324* | *0.384* | *B* | n.s. |
| Ursidae | *7* | *19.134* | *9.449 – 23.714* | *0.030* | *0.012 – 0.089* | *0.556* | *+* | *6* | *0.032* | *-0.002 – 0.066* | *0.511* | *B* | n.s. |
| Felidae | 26 | 18.184 | 16.218 – 20.654 | 0.043 | 0.030 – 0.054 | 0.525 | + | *25* | *0.042* | *0.025 – 0.059* | *0.245* | *+* | n.s. |
| Herpestidae | *12* | *43.421* | *39.811 – 124.738* | *-0.060* | *-0.208 – -0.045* | *0.004* | *–* | *11* | *0.056* | *0.016 – 0.096* | *0.183* | *+* | n.s. |
| Eupleridae | *5* | *19.996* | *10.046 – 33.963* | *0.041* | *-0.043 – 0.130* | *0.267* | *B* | | | | | | |
| Viverridae | *14* | *10.940* | *1.239 – 16.482* | *0.114* | *0.061 – 0.382* | *0.258* | *+* | *13* | *0.114* | *0.042 – 0.186* | *0.066* | *+* | n.s. |
| **Locomotor type** | | | | | | | | | | | | | |
| arboreal | *7* | *8.619* | *0.504 – 37.239* | *0.126* | *-0.071 – 0.478* | *0.095* | *B* | *6* | *-0.115* | *-0.253 – 0.023* | *0.261* | *B* | n.s. |
| semiarboreal | *10* | *17.235* | *9.829 – 25.527* | *0.056* | *-0.001 – 0.125* | *0.496* | *B* | 9 | 0.060 | 0.030 – 0.090 | 0.799 | + | n.s. |
| scansorial | 45 | 19.829 | 18.408 – 21.429 | 0.033 | 0.025 – 0.041 | 0.426 | + | *44* | *0.042* | *0.029 – 0.055* | *0.251* | *+* | n.s. |
| terrestrial | *49* | *38.380* | *35.156 – 81.283* | *-0.040* | *-0.132 – -0.029* | *0.002* | *–* | *48* | *0.051* | *0.036 – 0.066* | *0.130* | *+* | n.s. |
| semifossorial | *7* | *20.893* | *9.643 – 26.546* | *0.043* | *0.016 – 0.141* | *0.346* | *+* | *6* | *0.038* | *-0.008 – 0.084* | *0.249* | *B* | n.s. |
| semiaquatic | 11 | 18.143 | 15.704 –25.942 | 0.058 | 0.017 – 0.074 | 0.822 | + | 10 | 0.058 | 0.028 – 0.088 | 0.725 | + | × |
| aquatic | *8* | *20.970* | *4.901 – 36.224* | *0.046* | *0.001 – 0.164* | *0.486* | *+* | *7* | *0.049* | *$-8.77 \cdot 10^{-4}$ – 0.099* | *0.240* | *B* | n.s. |

| SR29 – $\%_{mid}$ | traditional regression | | | | | | | PIC regression | | | | | |
|---|---|---|---|---|---|---|---|---|---|---|---|---|---|
| | n | $a$ | 95% $CI_a$ | $b_{trad}$ | 95% $CI_b$ | R | sim. | n | $b_{PIC}$ | 95% $CI_b$ | R | sim. | $b_{trad} \neq b_{PIC}$ |
| **whole sample** | 137 | 57.597 | 54.325 – 61.094 | -0.046 | -0.053 – -0.039 | 0.595 | – | 136 | -0.044 | -0.050 – -0.038 | 0.508 | – | × |
| **fissipeds** | 130 | 53.530 | 50.699 – 55.976 | -0.037 | -0.042 – -0.030 | 0.418 | – | 129 | -0.039 | -0.045 – -0.033 | 0.454 | – | × |
| **Family** | | | | | | | | | | | | | |
| Canidae | 17 | 52.881 | 44.361 – 59.704 | -0.039 | -0.053 – -0.019 | 0.720 | – | 16 | -0.039 | -0.057 – -0.021 | 0.531 | – | × |
| Mustelidae | 32 | 51.916 | 48.529 – 55.081 | -0.031 | -0.039 – -0.022 | 0.701 | – | 31 | -0.028 | -0.036 – -0.020 | 0.681 | – | × |
| Procyonidae | 7 | 82.092 | 44.055 – 142.561 | -0.088 | -0.154 – -0.016 | 0.705 | – | 6 | -0.084 | -0.156 – -0.012 | 0.721 | – | n.s. |
| Ursidae | 7 | 63.416 | 34.514 – 175.388 | -0.040 | -0.127 – 0.013 | 0.290 | B | 6 | -0.046 | -0.102 – 0.010 | 0.070 | B | n.s. |
| Felidae | 26 | 46.957 | 42.756 – 72.277 | -0.022 | -0.068 – -0.012 | 0.193 | – | 25 | -0.030 | -0.042 – -0.018 | 0.328 | – | n.s. |
| Herpestidae | 12 | 53.174 | 42.756 – 86.298 | -0.043 | -0.108 – -0.011 | 0.473 | – | 11 | -0.040 | -0.064 – -0.016 | 0.554 | – | n.s. |
| Eupleridae | 5 | 23.025 | 4.328 – 37.757 | 0.070 | -0.011 – 0.309 | 0.581 | B | | | | | | |
| Viverridae | 14 | 71.945 | 50.003 – 272.898 | -0.074 | -0.239 – -0.030 | 0.299 | – | 13 | -0.057 | -0.090 – -0.024 | 0.424 | – | n.s. |
| **Locomotor type** | | | | | | | | | | | | | |
| arboreal | 7 | 65.253 | 27.290 – 158.489 | -0.052 | -0.158 – 0.061 | 0.184 | B | 6 | 0.024 | 0.019 – 0.029 | 0.984 | + | n.s. |
| semiarboreal | 10 | 50.548 | 41.305 – 87.297 | -0.028 | -0.099 – 0.002 | 0.017 | B | 9 | -0.026 | -0.047 – -0.005 | 0.241 | – | n.s. |
| scansorial | 45 | 47.676 | 45.082 – 73.790 | -0.022 | -0.071 – -0.016 | 0.102 | – | 44 | -0.031 | -0.040 – -0.022 | 0.328 | – | n.s. |
| terrestrial | 49 | 54.125 | 50.699 – 57.544 | -0.042 | -0.050 – -0.033 | 0.720 | – | 48 | -0.032 | -0.040 – -0.024 | 0.508 | – | ✓ |
| semifossorial | 7 | 46.644 | 42.658 – 71.450 | -0.025 | -0.078 – -0.012 | 0.324 | – | 6 | -0.026 | -0.055 – 0.003 | 0.444 | B | n.s. |
| semiaquatic | 11 | 56.312 | 48.529 – 131.522 | -0.039 | -0.136 – -0.018 | 0.397 | – | 10 | 0.030 | 0.011 – 0.049 | 0.604 | + | n.s. |
| aquatic | 8 | 92.066 | 68.391 – 163.682 | -0.091 | -0.139 – -0.068 | 0.929 | – | 7 | -0.089 | -0.137 – -0.041 | 0.857 | – | × |

| SR30 – %dist | traditional regression | | | | | | | PIC regression | | | | | |
|---|---|---|---|---|---|---|---|---|---|---|---|---|---|
| | n | *a* | 95% CI$_a$ | *b*$_{trad}$ | 95% CI$_b$ | R | sim. | n | *b*$_{PIC}$ | 95% CI$_b$ | R | sim. | *b*$_{trad}$ ≠ *b*$_{PIC}$ |
| **whole sample** | 137 | 22.631 | 21.429 – 24.099 | 0.044 | 0.037 – 0.051 | 0.187 | + | *136* | *0.050* | *0.041 – 0.059* | *0.098* | + | n.s. |
| **fissipeds** | 130 | 21.807 | 20.464 – 23.496 | 0.050 | 0.040 – 0.057 | 0.222 | + | *129* | *-0.063* | *-0.074 – -0.052* | *0.001* | – | n.s. |
| **Family** | | | | | | | | | | | | | |
| Canidae | *17* | *19.738* | *4.642 – 29.923* | *0.067* | *0.023 – 0.231* | *0.032* | + | *16* | *0.082* | *0.037 – 0.127* | *0.070* | + | n.s. |
| Mustelidae | *32* | *42.374* | *37.844 – 84.333* | *-0.042* | *-0.133 – -0.030* | *0.077* | – | *31* | *0.067* | *0.042 – 0.092* | *0.041* | + | n.s. |
| Procyonidae | *7* | *20.663* | *8.674 – 75.858* | *0.063* | *-0.088 – 0.172* | *0.475* | *B* | *6* | *0.062* | *-0.005 – 0.129* | *0.490* | *B* | n.s. |
| Ursidae | *7* | *47.610* | *34.995 – 168.267* | *-0.032* | *-0.140 – -0.003* | *0.005* | – | *6* | *-0.031* | *-0.069 – 0.007* | *0.071* | *B* | n.s. |
| Felidae | *26* | *51.109* | *44.771 – 115.611* | *-0.042* | *-0.131 – -0.029* | *0.331* | – | *25* | *0.046* | *0.027 – 0.065* | *0.035* | + | n.s. |
| Herpestidae | *12* | *18.030* | *4.642 – 22.961* | *0.082* | *0.044 – 0.273* | *0.312* | + | *11* | *0.076* | *0.024 – 0.128* | *0.287* | + | n.s. |
| Eupleridae | *5* | *57.438* | *24.831 – 189.234* | *-0.069* | *-0.237 – 0.062* | *0.797* | *B* | | | | | | |
| Viverridae | *14* | *45.930* | *36.644 – 109.901* | *-0.042* | *-0.148 – -0.011* | *0.053* | – | *13* | *0.104* | *0.039 – 0.169* | *0.191* | + | n.s. |
| **Locomotor type** | | | | | | | | | | | | | |
| arboreal | *7* | *20.142* | *3.648 – 79.433* | *0.062* | *-0.111 – 0.278* | *0.099* | *B* | *6* | *0.049* | *-0.010 – 0.108* | *0.264* | *B* | n.s. |
| semiarboreal | 10 | 40.105 | 36.644 – 42.855 | -0.026 | -0.034 – -0.014 | 0.792 | – | *9* | *0.072* | *0.012 – 0.132* | *0.026* | + | n.s. |
| scansorial | *45* | *45.867* | *42.658 – 80.538* | *-0.032* | *-0.095 – -0.024* | *0.267* | – | *44* | *0.048* | *0.033 – 0.063* | *0.069* | + | n.s. |
| terrestrial | 49 | 21.434 | 19.861 – 23.933 | 0.055 | 0.041 – 0.066 | 0.634 | + | *48* | *0.053* | *0.038 – 0.068* | *0.276* | + | n.s. |
| semifossorial | *7* | *44.066* | *37.325 – 95.060* | *-0.040* | *-0.146 – -0.016* | *0.094* | – | *6* | *0.029* | *-0.002 – 0.060* | *0.514* | *B* | n.s. |
| semiaquatic | *11* | *51.098* | *27.925 –80.353* | *-0.064* | *-0.113 – 0.002* | *0.400* | *B* | 10 | -0.072 | -0.088 – -0.056 | 0.957 | – | n.s. |
| aquatic | 8 | 12.791 | 2.191 – 25.351 | 0.078 | 0.035 – 0.226 | 0.749 | + | *7* | *0.079* | *0.001 – 0.157* | *0.368* | + | n.s. |

**Tables SR31 to SR59 – Results of the complex allometry tests**

In each case, it is indicated (in the "*D* ≠ *1*" column) whether the exponent of complex allometry (*D*) is significantly different from 1. Results in *grey italics* denote non-significant regressions. Variable names are listed in Table II.3. Abbreviations: 95% $CI_C$, 95% confidence interval for the coefficient (*C*); 95% $CI_D$, 95% confidence interval for the exponent of complex allometry (*D*); 95% $CI_{ln A}$, 95% confidence interval for *ln A*; n, sample size; n.c., the model did not converge in a realistic solution; n.s., although the model did converge in a realistic solution, it was not significant according to the associated correlation coefficient (R).

| **SR31 – $L_S$** | **n** | ***ln A*** | **95% $CI_{ln A}$** | ***C*** | **95% $CI_C$** | ***D*** | **95% $CI_D$** | **R** | ***D* ≠ 1** |
|---|---|---|---|---|---|---|---|---|---|
| **whole sample** | 137 | 5.527 | 5.377 – 5.677 | 0.181 | 0.108 – 0.253 | 1.284 | 1.102 – 1.465 | 0.960 | ✓ (*D*>1) |
| **fissipeds** | 130 | 5.568 | 5.390 – 5.746 | 0.270 | 0.167 – 0.373 | 1.129 | 0.957 – 1.300 | 0.958 | × |
| **Family** | | | | | | | | | |
| Canidae | 17 | 5.133 | 4.964 – 5.301 | 0.449 | 0.272 – 0.625 | 0.971 | 0.675 – 1.266 | 0.973 | × |
| Mustelidae | 32 | 4.604 | 4.392 – 4.817 | 0.383 | 0.188 – 0.578 | 0.911 | 0.641 – 1.182 | 0.957 | × |
| Procyonidae | 7 | 4.245 | 3.576 – 4.915 | 0.409 | -0.440 – 1.258 | 0.754 | 1.207 – 2.715 | 0.833 | × |
| Ursidae | 7 | 5.438 | 5.365 – 5.511 | 0.200 | 0.114 – 0.287 | 1.472 | 0.930 – 2.014 | 0.991 | × |
| Felidae | 26 | 5.458 | 5.330 – 5.586 | 0.360 | 0.236 – 0.484 | 0.908 | 0.713 – 1.103 | 0.982 | × |
| Herpestidae | 12 | 4.153 | 4.013 – 4.293 | 0.433 | 0.266 – 0.600 | 0.647 | 0.341 – 0.953 | 0.981 | ✓ (*D*<1) |
| Eupleridae | 5 | 4.236 | 3.718 – 4.753 | 0.166 | -0.361 – 0.693 | 1.728 | -1.648 – 5.104 | 0.962 | × |
| Viverridae | 14 | 4.400 | 4.215 – 4.585 | 0.237 | 0.040 – 0.434 | 1.120 | 0.223 – 1.818 | 0.908 | × |
| **Locomotor type** | | | | | | | | | |
| arboreal | 7 | 4.287 | 3.913 – 4.661 | 0.187 | -0.154 – 0.529 | 1.748 | -0.231 – 3.728 | 0.917 | × |
| semiarboreal | 10 | 4.518 | 4.326 – 4.710 | 0.368 | 0.150 – 0.586 | 0.820 | 0.393 – 1.246 | 0.974 | × |
| scansorial | 45 | 5.426 | 5.296 – 5.557 | 0.238 | 0.140 – 0.336 | 1.168 | 0.952 – 1.385 | 0.979 | × |
| terrestrial | 49 | 5.705 | 5.388 – 6.021 | 0.286 | 0.122 – 0.451 | 1.139 | 0.892 – 1.387 | 0.964 | × |
| semifossorial | 7 | 4.297 | 3.963 – 4.630 | 0.320 | -0.274 – 0.913 | 1.026 | -0.637 – 2.689 | 0.953 | × |
| semiaquatic | 11 | 4.312 | 4.025 – 4.598 | 0.093 | -0.117 – 0.302 | 1.863 | 0.151 – 3.575 | 0.904 | × |
| aquatic | 8 | 5.402 | 5.006 – 5.799 | 0.174 | -0.226 – 0.573 | 1.410 | -0.438 – 3.258 | 0.881 | × |

| **SR32 – S** | **n** | ***ln A*** | **95% $CI_{ln A}$** | ***C*** | **95% $CI_C$** | ***D*** | **95% $CI_D$** | **R** | ***D* ≠ 1** |
|---|---|---|---|---|---|---|---|---|---|
| **whole sample** | 137 | 4.941 | 4.737 – 5.146 | 0.443 | 0.301 – 0.585 | 0.918 | 0.784 – 1.052 | 0.967 | × |
| **fissipeds** | 130 | 4.581 | 4.421 – 4.742 | 0.360 | 0.253 – 0.466 | 0.985 | 0.857 – 1.113 | 0.971 | × |
| **Family** | | | | | | | | | |
| Canidae | 17 | 3.827 | 3.616 – 4.037 | 0.445 | 0.223 – 0.667 | 0.942 | 0.570 – 1.314 | 0.957 | × |
| Mustelidae | 32 | 3.826 | 3.630 – 4.021 | 0.485 | 0.297 – 0.672 | 0.814 | 0.615 – 1.012 | 0.971 | × |
| Procyonidae | 7 | 3.243 | 2.680 – 3.806 | 0.346 | -0.344 – 1.037 | 0.683 | -1.074 – 2.440 | 0.838 | × |
| Ursidae | 7 | 4.449 | 4.359 – 4.540 | 0.183 | 0.074 – 0.292 | 1.173 | 0.453 – 1.893 | 0.979 | × |
| Felidae | 26 | 4.272 | 4.140 – 4.405 | 0.282 | 0.163 – 0.401 | 1.062 | 0.814 – 1.310 | 0.979 | × |
| Herpestidae | 12 | 2.913 | 2.831 – 2.995 | 0.324 | 0.223 – 0.425 | 0.932 | 0.634 – 1.231 | 0.987 | × |
| Eupleridae | 5 | 3.210 | 2.951 – 3.468 | 0.351 | -0.014 – 0.716 | 0.940 | -0.169 – 2.049 | 0.992 | × |
| Viverridae | 14 | 3.444 | 3.225 – 3.663 | 0.422 | 0.163 – 0.680 | 0.643 | 0.199 – 1.088 | 0.935 | × |
| **Locomotor type** | | | | | | | | | |
| arboreal | 7 | 3.528 | 3.331 – 3.725 | 0.372 | 0.149 – 0.595 | 1.085 | 0.426 – 1.744 | 0.981 | × |
| semiarboreal | 10 | 3.552 | 3.347 – 3.757 | 0.435 | 0.199 – 0.672 | 0.737 | 0.358 – 1.115 | 0.975 | × |
| scansorial | 45 | 4.400 | 4.282 – 4.518 | 0.265 | 0.174 – 0.357 | 1.126 | 0.947 – 1.305 | 0.985 | × |
| terrestrial | 49 | 4.550 | 4.363 – 4.737 | 0.421 | 0.298 – 0.544 | 0.901 | 0.782 – 1.020 | 0.984 | × |
| semifossorial | 7 | 3.548 | 1.737 – 5.360 | 0.921 | -1.285 – 3.127 | 0.372 | -0.756 – 1.500 | 0.933 | × |
| semiaquatic | 11 | 3.771 | 3.501 – 4.040 | 0.425 | 0.147 – 0.704 | 0.861 | 0.439 – 1.284 | 0.963 | × |
| aquatic | 8 | – | – | – | – | – | – | – | n.c. |

| **SR33 – I** | **n** | ***ln A*** | **95% CI$_{ln A}$** | ***C*** | **95% CI$_C$** | ***D*** | **95% CI$_D$** | **R** | ***D* ≠ 1** |
|---|---|---|---|---|---|---|---|---|---|
| **whole sample** | 137 | 4.807 | 4.580 – 5.034 | 0.194 | 0.088 – 0.300 | 1.315 | 1.067 – 1.563 | 0.931 | ✓ (*D*>1) |
| **fissipeds** | 130 | 4.816 | 4.549 – 5.083 | 0.280 | 0.131 – 0.428 | 1.171 | 0.931 – 1.412 | 0.925 | × |
| **Family** | | | | | | | | | |
| Canidae | 17 | 4.178 | 3.924 – 4.432 | 0.453 | 0.181 – 0.725 | 0.847 | 0.415 – 1.278 | 0.936 | × |
| Mustelidae | 32 | 3.680 | 3.363 – 3.997 | 0.448 | 0.162 – 0.734 | 0.954 | 0.611 – 1.297 | 0.938 | × |
| Procyonidae | 7 | 3.478 | 3.214 – 3.742 | 0.197 | -0.187 – 0.580 | 1.057 | -1.220 – 3.334 | 0.885 | × |
| Ursidae | 7 | 4.806 | 4.578 – 5.035 | 0.200 | -0.073 – 0.473 | 0.859 | -0.649 – 2.367 | 0.885 | × |
| Felidae | 26 | 4.585 | 4.431 – 4.739 | 0.334 | 0.190 – 0.479 | 0.977 | 0.728 – 1.226 | 0.975 | × |
| Herpestidae | 12 | 3.305 | 3.094 – 3.515 | 0.444 | 0.193 – 0.695 | 0.631 | 0.191 – 1.071 | 0.961 | × |
| Eupleridae | 5 | 3.395 | 2.527 – 4.263 | 0.273 | -0.697 – 1.243 | 1.506 | -2.279 – 5.292 | 0.944 | × |
| Viverridae | 14 | 3.791 | 3.543 – 4.040 | 0.456 | 0.163 – 0.749 | 0.557 | 0.125 – 0.989 | 0.932 | ✓ (*D*<1) |
| **Locomotor type** | | | | | | | | | |
| arboreal | 7 | 3.845 | 3.642 – 4.048 | 0.393 | 0.142 – 0.643 | 0.850 | 0.154 – 1.547 | 0.976 | × |
| semiarboreal | 10 | 3.698 | 3.290 – 4.107 | 0.434 | -0.040 – 0.908 | 0.689 | -0.056 – 1.433 | 0.905 | × |
| scansorial | 45 | 4.695 | 4.526 – 4.863 | 0.238 | 0.118 – 0.358 | 1.237 | 0.969 – 1.504 | 0.971 | × |
| terrestrial | 49 | 4.942 | 4.441 – 5.442 | 0.326 | 0.065 – 0.588 | 1.134 | 0.789 – 1.478 | 0.933 | × |
| semifossorial | 7 | 3.291 | 2.880 – 3.701 | 0.121 | -0.232 – 0.474 | 2.165 | -0.506 – 4.835 | 0.934 | × |
| semiaquatic | 11 | – | – | – | – | – | – | – | n.c. |
| aquatic | 8 | – | – | – | – | – | – | – | n.c. |

| **SR34 – A** | **n** | ***ln A*** | **95% CI$_{ln A}$** | ***C*** | **95% CI$_C$** | ***D*** | **95% CI$_D$** | **R** | ***D* ≠ 1** |
|---|---|---|---|---|---|---|---|---|---|
| **whole sample** | 137 | 5.619 | 5.447 – 5.791 | 0.320 | 0.217 – 0.424 | 1.075 | 0.934 – 1.215 | 0.971 | × |
| **fissipeds** | 130 | 5.435 | 5.257 – 5.614 | 0.331 | 0.220 – 0.441 | 1.059 | 0.911 – 1.206 | 0.965 | × |
| **Family** | | | | | | | | | |
| Canidae | 17 | 4.735 | 4.519 – 4.950 | 0.454 | 0.226 – 0.683 | 0.900 | 0.530 – 1.269 | 0.955 | × |
| Mustelidae | 32 | 4.461 | 4.226 – 4.696 | 0.455 | 0.237 – 0.673 | 0.889 | 0.637 – 1.142 | 0.960 | × |
| Procyonidae | 7 | 4.059 | 3.784 – 4.334 | 0.271 | -0.113 – 0.654 | 0.969 | -0.606 – 2.543 | 0.927 | × |
| Ursidae | 7 | 5.347 | 5.202 – 5.492 | 0.200 | 0.027 – 0.373 | 0.869 | -0.092 – 1.830 | 0.949 | × |
| Felidae | 26 | 5.151 | 5.028 – 5.275 | 0.313 | 0.199 – 0.427 | 1.015 | 0.804 – 1.227 | 0.983 | × |
| Herpestidae | 12 | 3.814 | 3.711 – 3.917 | 0.353 | 0.226 – 0.480 | 0.816 | 0.490 – 1.142 | 0.982 | × |
| Eupleridae | 5 | 3.982 | 3.606 – 4.359 | 0.253 | -0.185 – 0.691 | 1.410 | -0.434 – 3.253 | 0.984 | × |
| Viverridae | 14 | 4.284 | 4.116 – 4.451 | 0.390 | 0.193 – 0.587 | 0.695 | 0.317 – 1.073 | 0.954 | × |
| **Locomotor type** | | | | | | | | | |
| arboreal | 7 | 4.344 | 4.187 – 4.501 | 0.361 | 0.177 – 0.545 | 0.976 | 0.414 – 1.538 | 0.985 | × |
| semiarboreal | 10 | 4.341 | 4.177 – 4.505 | 0.452 | 0.262 – 0.643 | 0.670 | 0.386 – 0.955 | 0.984 | ✓ (*D*<1) |
| scansorial | 45 | 5.266 | 5.141 – 5.391 | 0.252 | 0.159 – 0.345 | 1.185 | 0.991 – 1.379 | 0.984 | × |
| terrestrial | 49 | 5.483 | 5.186 – 5.781 | 0.375 | 0.199 – 0.550 | 1.014 | 0.817 – 1.210 | 0.970 | × |
| semifossorial | 7 | 4.052 | 3.548 – 4.556 | 0.458 | -0.440 – 1.356 | 0.850 | -0.811 – 2.511 | 0.948 | × |
| semiaquatic | 11 | 4.290 | 3.971 – 4.609 | 0.250 | -0.043 – 0.544 | 1.293 | 0.461 – 2.124 | 0.943 | × |
| aquatic | 8 | – | – | – | – | – | – | – | n.c. |

| **SR35 – $H_S$** | **n** | ***ln A*** | **95% $CI_{ln A}$** | ***C*** | **95% $CI_C$** | ***D*** | **95% $CI_D$** | **R** | ***D* ≠ 1** |
|---|---|---|---|---|---|---|---|---|---|
| **whole sample** | 137 | 3.235 | 3.010 – 3.460 | 0.041 | $3.76 \cdot 10^{-4}$ – 0.082 | 2.049 | 1.570 – 2.528 | 0.860 | ✓ (*D*>1) |
| **fissipeds** | 130 | 3.719 | 3.412 – 4.025 | 0.213 | 0.072 – 0.354 | 1.352 | 1.043 – 1.661 | 0.902 | ✓ (*D*>1) |
| **Family** | | | | | | | | | |
| Canidae | 17 | 3.258 | 2.991 – 3.524 | 0.536 | 0.252 – 0.821 | 0.871 | 0.486 – 1.257 | 0.950 | × |
| Mustelidae | 32 | 2.579 | 2.141 – 3.018 | 0.248 | -0.057 – 0.553 | 1.375 | 0.660 – 2.091 | 0.868 | × |
| Procyonidae | 7 | 2.424 | 1.717 – 3.131 | 0.459 | -0.535 – 1.453 | 0.984 | -1.447 – 3.415 | 0.852 | × |
| Ursidae | 7 | 3.644 | 3.412 – 3.876 | 0.132 | -0.139 – 0.403 | 1.599 | -1.010 – 4.207 | 0.857 | × |
| Felidae | 26 | 3.507 | 3.308 – 3.705 | 0.182 | 0.043 – 0.320 | 1.471 | 0.996 – 1.946 | 0.960 | × |
| Herpestidae | 12 | 2.253 | 1.857 – 2.649 | 0.480 | 0.035 – 0.925 | 0.448 | -0.101 – 0.997 | 0.923 | ✓ (*D*=0) |
| Eupleridae | 5 | 2.169 | 1.861 – 2.478 | 0.106 | -0.119 – 0.332 | 2.454 | 0.194 – 4.713 | 0.991 | × |
| Viverridae | 14 | 2.471 | 2.272 – 2.669 | 0.279 | 0.084 – 0.475 | 1.339 | 0.742 – 1.936 | 0.943 | × |
| **Locomotor type** | | | | | | | | | |
| arboreal | 7 | 2.457 | 2.043 – 2.870 | 0.307 | -0.129 – 0.743 | 1.308 | -0.243 – 2.859 | 0.921 | × |
| semiarboreal | 10 | 2.679 | 2.257 – 3.100 | 0.514 | 0.035 – 0.992 | 0.833 | 0.159 – 1.508 | 0.939 | × |
| scansorial | 45 | 3.626 | 3.469 – 3.783 | 0.202 | 0.102 – 0.301 | 1.375 | 1.108 – 1.643 | 0.975 | ✓ (*D*>1) |
| terrestrial | 49 | 3.758 | 3.217 – 4.299 | 0.201 | -0.012 – 0.413 | 1.395 | 0.920 – 1.869 | 0.924 | × |
| semifossorial | 7 | – | – | – | – | – | – | – | n.c. |
| semiaquatic | 11 | – | – | – | – | – | – | – | n.c. |
| aquatic | 8 | – | – | – | – | – | – | – | n.c. |

| **SR36 – $L_h$** | **n** | ***ln A*** | **95% $CI_{ln A}$** | ***C*** | **95% $CI_C$** | ***D*** | **95% $CI_D$** | **R** | ***D* ≠ 1** |
|---|---|---|---|---|---|---|---|---|---|
| **whole sample** | 137 | 5.513 | 5.363 – 5.663 | 0.081 | 0.031 – 0.132 | 1.583 | 1.290 – 1.876 | 0.922 | ✓ (*D*>1) |
| **fissipeds** | 130 | 5.868 | 5.673 – 6.063 | 0.277 | 0.159 – 0.395 | 1.084 | 0.895 – 1.274 | 0.947 | × |
| **Family** | | | | | | | | | |
| Canidae | 17 | 5.402 | 5.133 – 5.672 | 0.417 | 0.131 – 0.702 | 0.905 | 0.400 – 1.409 | 0.921 | × |
| Mustelidae | 32 | 4.722 | 4.501 – 4.944 | 0.208 | 0.029 – 0.387 | 1.152 | 0.667 – 1.637 | 0.914 | × |
| Procyonidae | 7 | 4.610 | 4.321 – 4.898 | 0.257 | -0.106 – 0.620 | 0.734 | -0.577 – 2.044 | 0.910 | × |
| Ursidae | 7 | 5.804 | 5.692 – 5.917 | 0.211 | 0.076 – 0.346 | 1.215 | 0.434 – 1.995 | 0.977 | × |
| Felidae | 26 | 5.688 | 5.574 – 5.802 | 0.281 | 0.175 – 0.386 | 0.999 | 0.782 – 1.217 | 0.982 | × |
| Herpestidae | 12 | 4.467 | 4.215 – 4.719 | 0.435 | 0.139 – 0.732 | 0.587 | 0.083 – 1.091 | 0.946 | × |
| Eupleridae | 5 | 4.665 | 4.555 – 4.774 | 0.289 | 0.149 – 0.428 | 1.179 | 0.662 – 1.696 | 0.998 | × |
| Viverridae | 14 | 4.098 | 4.399 – 5.798 | 0.686 | -0.064 – 1.436 | 0.256 | -0.116 – 0.626 | 0.932 | ✓ (*D*=0) |
| **Locomotor type** | | | | | | | | | |
| arboreal | 7 | 4.957 | 4.662 – 5.251 | 0.380 | 0.005 – 0.755 | 0.768 | -0.294 – 1.830 | 0.946 | × |
| semiarboreal | 10 | 4.940 | 4.803 – 5.076 | 0.391 | 0.233 – 0.548 | 0.704 | 0.426 – 0.981 | 0.985 | ✓ (*D*<1) |
| scansorial | 45 | 5.720 | 5.615 – 5.826 | 0.206 | 0.130 – 0.282 | 1.216 | 1.020 – 1.413 | 0.984 | ✓ (*D*>1) |
| terrestrial | 49 | 5.977 | 5.657 – 6.296 | 0.307 | 0.130 – 0.485 | 1.073 | 0.828 – 1.319 | 0.960 | × |
| semifossorial | 7 | 4.536 | 4.181 – 4.891 | 0.311 | -0.325 – 0.947 | 0.986 | -0.832 – 2.803 | 0.944 | × |
| semiaquatic | 11 | 4.555 | 4.330 – 4.780 | 0.078 | -0.092 – 0.247 | 1.807 | 0.164 – 3.450 | 0.906 | × |
| aquatic | 8 | – | – | – | – | – | – | – | n.c. |

| SR37 – $d_{sh}$ | n | *ln A* | 95% $CI_{ln A}$ | *C* | 95% $CI_C$ | *D* | 95% $CI_D$ | R | $D \neq 1$ |
|---|---|---|---|---|---|---|---|---|---|
| **whole sample** | 137 | 3.985 | 3.824 – 4.146 | 0.348 | 0.246 – 0.449 | 1.023 | 0.898 – 1.149 | 0.975 | × |
| **fissipeds** | 130 | 3.842 | 3.675 – 4.008 | 0.358 | 0.251 – 0.465 | 1.019 | 0.888 – 1.150 | 0.971 | × |
| **Family** | | | | | | | | | |
| Canidae | 17 | 3.056 | 2.850 – 3.261 | 0.415 | 0.207 – 0.623 | 1.100 | 0.710 – 1.489 | 0.960 | × |
| Mustelidae | 32 | 3.047 | 2.773 – 3.321 | 0.511 | 0.251 – 0.772 | 0.838 | 0.574 – 1.101 | 0.952 | × |
| Procyonidae | 7 | 2.529 | 2.137 – 2.921 | 0.338 | -0.216 – 0.891 | 0.993 | -0.856 – 2.842 | 0.908 | × |
| Ursidae | 7 | 3.691 | 3.617 – 3.766 | 0.286 | 0.197 – 0.376 | 1.025 | 0.658 – 1.391 | 0.993 | × |
| Felidae | 26 | 3.660 | 3.522 – 3.799 | 0.384 | 0.256 – 0.512 | 1.004 | 0.811 – 1.198 | 0.985 | × |
| Herpestidae | 12 | 2.121 | 1.960 – 2.282 | 0.329 | 0.131 – 0.527 | 0.862 | 0.302 – 1.422 | 0.953 | × |
| Eupleridae | 5 | 2.545 | 2.462 – 2.629 | 0.304 | 0.214 – 0.394 | 1.596 | 1.282 – 1.910 | 0.999 | ✓ (*D*>1) |
| Viverridae | 14 | 2.632 | 2.313 – 2.952 | 0.476 | 0.099 – 0.853 | 0.642 | 0.068 – 1.216 | 0.898 | × |
| **Locomotor type** | | | | | | | | | |
| arboreal | 7 | 2.715 | 2.609 – 2.822 | 0.313 | 0.197 – 0.428 | 1.217 | 0.812 – 1.621 | 0.993 | × |
| semiarboreal | 10 | 2.843 | 2.628 – 3.058 | 0.511 | 0.267 – 0.755 | 0.825 | 0.480 – 1.170 | 0.983 | × |
| scansorial | 45 | 3.625 | 3.487 – 3.764 | 0.261 | 0.158 – 0.363 | 1.192 | 0.986 – 1.399 | 0.982 | × |
| terrestrial | 49 | 3.910 | 3.613 – 4.206 | 0.447 | 0.255 – 0.639 | 0.919 | 0.743 – 1.095 | 0.969 | × |
| semifossorial | 7 | 2.382 | 2.136 – 2.628 | 0.215 | -0.149 – 0.580 | 1.387 | -0.184 – 2.958 | 0.964 | × |
| semiaquatic | 11 | 2.858 | 2.456 – 3.259 | 0.242 | -0.123 – 0.607 | 1.329 | 0.254 – 2.404 | 0.915 | × |
| aquatic | 8 | 4.227 | 3.651 – 4.803 | 0.716 | 0.071 – 1.361 | 0.504 | -0.054 – 1.062 | 0.936 | × |

| SR38 – $d_{th}$ | n | *ln A* | 95% $CI_{ln A}$ | *C* | 95% $CI_C$ | *D* | 95% $CI_D$ | R | $D \neq 1$ |
|---|---|---|---|---|---|---|---|---|---|
| **whole sample** | 137 | 3.808 | 3.626 – 3.990 | 0.439 | 0.312 – 0.565 | 0.915 | 0.795 – 1.035 | 0.973 | × |
| **fissipeds** | 130 | 3.649 | 3.462 – 3.836 | 0.463 | 0.328 – 0.598 | 0.887 | 0.764 – 1.009 | 0.969 | × |
| **Family** | | | | | | | | | |
| Canidae | 17 | 2.849 | 2.539 – 3.105 | 0.491 | 0.217 – 0.765 | 0.850 | 0.448 – 1.253 | 0.944 | × |
| Mustelidae | 32 | 2.379 | 2.170 – 2.588 | 0.328 | 0.138 – 0.518 | 0.933 | 0.624 – 1.242 | 0.947 | × |
| Procyonidae | 7 | 2.143 | 1.923 – 2.362 | 0.267 | -0.036 – 0.569 | 0.941 | -0.299 – 2.182 | 0.949 | × |
| Ursidae | 7 | 3.418 | 3.292 – 3.545 | 0.127 | -0.021 – 0.274 | 1.613 | 0.135 – 3.090 | 0.947 | × |
| Felidae | 26 | 3.246 | 3.103 – 3.389 | 0.315 | 0.186 – 0.444 | 1.059 | 0.819 – 1.298 | 0.980 | × |
| Herpestidae | 12 | 1.762 | 1.614 – 1.909 | 0.210 | 0.029 – 0.391 | 1.054 | 0.193 – 1.916 | 0.914 | × |
| Eupleridae | 5 | 2.246 | 1.853 – 2.638 | 0.407 | -0.116 – 0.929 | 1.074 | -0.301 – 2.450 | 0.988 | × |
| Viverridae | 14 | 2.398 | 2.152 – 2.645 | 0.490 | 0.198 – 0.781 | 0.613 | 0.192 – 1.035 | 0.939 | × |
| **Locomotor type** | | | | | | | | | |
| arboreal | 7 | 2.403 | 2.228 – 2.577 | 0.353 | 0.183 – 0.569 | 0.841 | 0.175 – 1.507 | 0.978 | × |
| semiarboreal | 10 | 2.479 | 2.198 – 2.761 | 0.494 | 0.169 – 0.819 | 0.731 | 0.274 – 1.189 | 0.964 | × |
| scansorial | 45 | 3.435 | 3.303 – 3.566 | 0.323 | 0.217 – 0.428 | 1.076 | 0.908 – 1.244 | 0.986 | × |
| terrestrial | 49 | 3.572 | 3.289 – 3.855 | 0.436 | 0.251 – 0.621 | 0.908 | 0.734 – 1.081 | 0.968 | × |
| semifossorial | 7 | 2.102 | 1.872 – 2.332 | 0.225 | -0.134 – 0.584 | 1.304 | -0.172 – 2.779 | 0.967 | × |
| semiaquatic | 11 | 2.207 | 1.906 – 2.508 | 0.202 | -0.091 – 0.495 | 1.107 | 0.111 – 2.103 | 0.893 | × |
| aquatic | 8 | 3.593 | 3.090 – 4.096 | 0.179 | -0.297 – 0.656 | 1.730 | -0.485 – 3.946 | 0.901 | × |

| **SR39 – HR** | **n** | ***ln A*** | **95% CI$_{ln A}$** | ***C*** | **95% CI$_{C}$** | ***D*** | **95% CI$_{D}$** | **R** | ***D* ≠ 1** |
|---|---|---|---|---|---|---|---|---|---|
| **whole sample** | 137 | 4.224 | -25.155 – 33.603 | 6.108 | -23.324 – 35.540 | 0.038 | -0.144 – 0.219 | 0.607 | ✓ (*D*=0) |
| **fissipeds** | 130 | -2.049 | -2.351 – -1.746 | 0.070 | -0.173 – 0.312 | 0.749 | -0.633 – 2.130 | 0.292 | × |
| **Family** | | | | | | | | | |
| Canidae | 17 | -2.320 | -2.495 – -2.145 | 0.032 | -0.089 – 0.152 | 2.042 | -1.137 – 5.222 | 0.548 | × |
| Mustelidae | 32 | -1.509 | -2.425 – -0.592 | 0.486 | -0.484 – 1.457 | 0.330 | -0.303 – 0.963 | 0.620 | ✓ (*D*=0) |
| Procyonidae | *7* | *-2.055* | *-2.378 – -1.733* | *0.108* | *-0.408 – 0.624* | *1.296* | *-4.827 – 7.419* | *0.681* | n.s. |
| Ursidae | 7 | – | – | – | – | – | – | – | n.c. |
| Felidae | 26 | -2.026 | -2.182 – -1.871 | 0.105 | -0.039 – 0.248 | 1.011 | 0.219 – 1.803 | 0.823 | × |
| Herpestidae | 12 | – | – | – | – | – | – | – | n.c. |
| Eupleridae | 5 | -2.118 | -2.260 – -1.977 | 0.033 | -0.048 – 0.114 | 2.911 | 0.284 – 5.537 | 0.991 | × |
| Viverridae | 14 | -2.209 | -2.365 – -2.053 | 0.078 | -0.087 – 0.242 | 1.155 | -0.628 – 2.939 | 0.653 | × |
| **Locomotor type** | | | | | | | | | |
| arboreal | 7 | – | – | – | – | – | – | – | n.c. |
| semiarboreal | 10 | -2.079 | -2.357 – -1.801 | 0.142 | -0.163 – 0.447 | 1.017 | -0.621 – 2.656 | 0.806 | × |
| scansorial | 45 | -2.099 | -2.268 – -1.930 | 0.053 | -0.080 – 0.186 | 1.101 | -0.192 – 2.394 | 0.616 | × |
| terrestrial | 49 | – | – | – | – | – | – | – | n.c. |
| semifossorial | 7 | – | – | – | – | – | – | – | n.c. |
| semiaquatic | 11 | *-1.743* | *-2.269 – -1.217* | *0.120* | *-0.382 – 0.622* | *1.171* | *-1.742 – 4.083* | *0.586* | n.s. |
| aquatic | 8 | – | – | – | – | – | – | – | n.c. |

| **SR40 – L$_{r}$** | **n** | ***ln A*** | **95% CI$_{ln A}$** | ***C*** | **95% CI$_{C}$** | ***D*** | **95% CI$_{D}$** | **R** | ***D* ≠ 1** |
|---|---|---|---|---|---|---|---|---|---|
| **whole sample** | 137 | 5.470 | 5.287 – 5.653 | 0.088 | 0.027 – 0.149 | 1.600 | 1.277 – 1.924 | 0.909 | ✓ (*D*>1) |
| **fissipeds** | 130 | 5.654 | 5.414 – 5.894 | 0.202 | 0.081 – 0.322 | 1.266 | 0.990 – 1.543 | 0.913 | × |
| **Family** | | | | | | | | | |
| Canidae | 17 | 5.444 | 5.060 – 5.829 | 0.482 | 0.070 – 0.893 | 0.854 | 0.238 – 1.471 | 0.882 | × |
| Mustelidae | 32 | 4.431 | 4.185 – 4.678 | 0.184 | -0.002 – 0.370 | 1.261 | 0.684 – 1.839 | 0.897 | × |
| Procyonidae | 7 | 4.667 | 4.153 – 5.182 | 0.533 | -0.039 – 1.104 | 0.460 | -0.227 – 1.147 | 0.950 | × |
| Ursidae | 7 | 5.613 | 5.534 – 5.693 | 0.209 | 0.114 – 0.304 | 1.285 | 0.723 – 1.846 | 0.988 | × |
| Felidae | 26 | 5.531 | 5.333 – 5.730 | 0.245 | 0.066 – 0.423 | 1.060 | 0.632 – 1.489 | 0.941 | × |
| Herpestidae | 12 | 4.288 | 3.934 – 4.643 | 0.428 | -0.002 – 0.858 | 0.727 | -0.130 – 1.584 | 0.885 | × |
| Eupleridae | 5 | 4.417 | 3.765 – 5.070 | 0.155 | -0.571 – 0.881 | 1.517 | -3.462 – 6.496 | 0.909 | × |
| Viverridae | 14 | 4.724 | 4.484 – 4.964 | 0.463 | 0.183 – 0.743 | 0.488 | 0.114 – 0.862 | 0.944 | ✓ (*D*<1) |
| **Locomotor type** | | | | | | | | | |
| arboreal | 7 | 4.692 | 4.306 – 5.078 | 0.346 | -0.113 – 0.805 | 0.943 | -0.516 – 2.402 | 0.910 | × |
| semiarboreal | 10 | 4.678 | 4.535 – 4.821 | 0.362 | 0.197 – 0.527 | 0.649 | 0.386 – 1.012 | 0.981 | × |
| scansorial | 45 | 5.538 | 5.386 – 5.691 | 0.173 | 0.070 – 0.276 | 1.300 | 0.981 – 1.619 | 0.962 | × |
| terrestrial | 49 | 5.909 | 5.496 – 6.322 | 0.238 | 0.049 – 0.428 | 1.258 | 0.909 – 1.606 | 0.946 | × |
| semifossorial | 7 | – | – | – | – | – | – | – | n.c. |
| semiaquatic | 11 | – | – | – | – | – | – | – | n.c. |
| aquatic | 8 | 5.395 | 5.001 – 5.788 | 0.287 | -0.128 – 0.703 | 1.078 | -0.026 – 2.181 | 0.910 | × |

| SR41 – $d_{sr}$ | n | *ln A* | 95% $CI_{ln A}$ | *C* | 95% $CI_C$ | *D* | 95% $CI_D$ | R | $D \neq 1$ |
|---|---|---|---|---|---|---|---|---|---|
| **whole sample** | 137 | 3.112 | 2.928 – 3.296 | 0.326 | 0.207 – 0.445 | 0.994 | 0.838 – 1.149 | 0.961 | × |
| **fissipeds** | 130 | 3.017 | 2.824 – 3.209 | 0.363 | 0.232 – 0.494 | 0.952 | 0.797 – 1.108 | 0.956 | × |
| **Family** | | | | | | | | | |
| Canidae | 17 | 2.466 | 2.210 – 2.721 | 0.415 | 0.152 – 0.678 | 1.039 | 0.554 – 1.525 | 0.936 | × |
| Mustelidae | 32 | 2.013 | 1.772 – 2.255 | 0.266 | 0.061 – 0.471 | 1.069 | 0.644 – 1.495 | 0.923 | × |
| Procyonidae | 7 | 1.774 | 1.352 – 2.196 | 0.374 | -0.121 – 0.869 | 0.582 | -0.450 – 1.614 | 0.918 | × |
| Ursidae | 7 | 2.780 | 2.541 – 3.018 | 0.120 | -0.138 – 0.379 | 2.206 | -0.579 – 4.992 | 0.902 | × |
| Felidae | 26 | 2.831 | 2.635 – 3.026 | 0.416 | 0.229 – 0.602 | 0.947 | 0.691 – 1.203 | 0.973 | × |
| Herpestidae | 12 | 1.528 | 1.268 – 1.789 | 0.284 | -0.037 – 0.604 | 0.976 | -0.127 – 2.079 | 0.861 | × |
| Eupleridae | 5 | 1.624 | 1.330 – 1.918 | 0.210 | -0.125 – 0.546 | 1.454 | -0.247 – 3.155 | 0.987 | × |
| Viverridae | 14 | 2.024 | 1.845 – 2.203 | 0.526 | 0.316 – 0.737 | 0.546 | 0.280 – 0.812 | 0.972 | ✓ ($D<1$) |
| **Locomotor type** | | | | | | | | | |
| arboreal | 7 | 2.059 | 1.428 – 2.689 | 0.635 | -0.149 – 1.418 | 0.367 | -0.433 – 1.167 | 0.974 | × |
| semiarboreal | 10 | 1.700 | 1.541 – 1.860 | 0.169 | 0.002 – 0.335 | 1.210 | 0.429 – 1.990 | 0.960 | × |
| scansorial | 45 | 2.864 | 2.679 – 3.050 | 0.343 | 0.186 – 0.500 | 0.989 | 0.759 – 1.219 | 0.971 | × |
| terrestrial | 49 | 3.084 | 2.776 – 3.393 | 0.336 | 0.153 – 0.519 | 1.009 | 0.781 – 1.237 | 0.959 | × |
| semifossorial | 7 | 1.769 | 1.509 – 2.029 | 0.198 | -0.162 – 0.558 | 1.485 | -0.203 – 3.174 | 0.960 | × |
| semiaquatic | 11 | 2.114 | 1.746 – 2.482 | 0.309 | -0.049 – 0.668 | 1.096 | 0.301 – 1.890 | 0.926 | × |
| aquatic | 8 | 3.011 | 2.414 – 3.608 | 0.236 | -0.345 – 0.816 | 1.607 | -0.422 – 3.635 | 0.898 | × |

| SR42 – $d_{tr}$ | n | *ln A* | 95% $CI_{ln A}$ | *C* | 95% $CI_C$ | *D* | 95% $CI_D$ | R | $D \neq 1$ |
|---|---|---|---|---|---|---|---|---|---|
| **whole sample** | 137 | 4.031 | 3.757 – 4.304 | 0.590 | 0.391 – 0.789 | 0.862 | 0.725 – 1.000 | 0.963 | × |
| **fissipeds** | 130 | 3.467 | 3.234 – 3.700 | 0.384 | 0.235 – 0.534 | 1.023 | 0.853 – 1.193 | 0.953 | × |
| **Family** | | | | | | | | | |
| Canidae | 17 | 2.897 | 2.634 – 3.160 | 0.538 | 0.256 – 0.820 | 0.844 | 0.468 – 1.220 | 0.950 | × |
| Mustelidae | 32 | 2.365 | 2.088 – 2.642 | 0.580 | 0.304 – 0.856 | 0.707 | 0.474 – 0.940 | 0.950 | ✓ ($D<1$) |
| Procyonidae | 7 | 1.956 | 1.574 – 2.338 | 0.315 | -0.232 – 0.861 | 1.022 | -0.970 – 3.014 | 0.901 | × |
| Ursidae | 7 | 3.330 | 2.976 – 3.685 | 0.341 | -0.083 – 0.766 | 0.927 | -0.487 – 2.341 | 0.904 | × |
| Felidae | 26 | 3.210 | 3.086 – 3.334 | 0.352 | 0.240 – 0.464 | 1.053 | 0.867 – 1.240 | 0.988 | × |
| Herpestidae | 12 | 1.592 | 0.867 – 2.318 | 0.432 | -0.391 – 1.254 | 0.472 | -0.710 – 1.653 | 0.751 | × |
| Eupleridae | 5 | 1.998 | 1.663 – 2.334 | 0.386 | -0.049 – 0.826 | 1.127 | -0.080 – 2.334 | 0.991 | × |
| Viverridae | 14 | 2.303 | 1.897 – 2.709 | 0.735 | 0.271 – 1.199 | 0.401 | 0.066 – 0.736 | 0.951 | ✓ ($D<1$) |
| **Locomotor type** | | | | | | | | | |
| arboreal | 7 | 2.194 | 1.832 – 2.557 | 0.366 | -0.061 – 0.793 | 0.967 | -0.317 – 2.250 | 0.929 | × |
| semiarboreal | 10 | 2.363 | 2.019 – 2.707 | 0.658 | 0.255 – 1.062 | 0.590 | 0.196 – 0.983 | 0.964 | ✓ ($D<1$) |
| scansorial | 45 | 3.240 | 3.088 – 3.392 | 0.290 | 0.174 – 0.405 | 1.159 | 0.951 – 1.368 | 0.981 | × |
| terrestrial | 49 | 3.580 | 3.141 – 4.019 | 0.390 | 0.139 – 0.641 | 1.045 | 0.773 – 1.317 | 0.948 | × |
| semifossorial | 7 | – | – | – | – | – | – | – | n.c. |
| semiaquatic | 11 | 2.130 | 1.806 – 2.453 | 0.422 | 0.077 – 0.767 | 0.704 | 0.206 – 1.203 | 0.933 | × |
| aquatic | 8 | 4.097 | 3.622 – 4.572 | 0.444 | -0.051 – 0.939 | 1.180 | 0.314 – 2.046 | 0.952 | × |

| SR43 – P | n | *ln A* | 95% CI$_{\ln A}$ | *C* | 95% CI$_{C}$ | *D* | 95% CI$_{D}$ | R | *D* ≠ 1 |
|---|---|---|---|---|---|---|---|---|---|
| **whole sample** | 137 | 3.190 | 2.960 – 3.421 | 0.357 | 0.216 – 0.497 | 1.057 | 0.887 – 1.228 | 0.957 | × |
| **fissipeds** | 130 | 3.269 | 2.996 – 3.542 | 0.521 | 0.328 – 0.714 | 0.909 | 0.752 – 1.066 | 0.952 | × |
| **Family** | | | | | | | | | |
| Canidae | 17 | 2.086 | 1.949 – 2.223 | 0.283 | 0.162 – 0.405 | 1.477 | 1.127 – 1.827 | 0.977 | ✓ (*D*>1) |
| Mustelidae | 32 | 1.672 | 1.421 – 1.923 | 0.291 | 0.077 – 0.505 | 1.062 | 0.652 – 1.468 | 0.929 | × |
| Procyonidae | 7 | 1.470 | 0.757 – 2.183 | 0.422 | -0.426 – 1.270 | 0.612 | -0.018 – 2.242 | 0.835 | × |
| Ursidae | 7 | 3.253 | 2.791 – 3.726 | 0.533 | -0.005 – 1.071 | 0.561 | -0.329 – 1.452 | 0.937 | × |
| Felidae | 26 | 2.823 | 2.652 – 2.994 | 0.355 | 0.198 – 0.511 | 1.026 | 0.769 – 1.283 | 0.976 | × |
| Herpestidae | 12 | 1.500 | 1.020 – 1.980 | 0.665 | 0.127 – 1.204 | 0.441 | -0.032 – 0.914 | 0.941 | ✓ (*D*=0) |
| Eupleridae | 5 | 1.480 | 1.136 – 1.823 | 0.114 | -0.093 – 0.321 | 2.844 | 0.910 – 4.778 | 0.995 | × |
| Viverridae | 14 | 1.608 | 1.218 – 1.997 | 0.396 | -0.046 – 0.839 | 0.894 | -0.010 – 1.799 | 0.829 | × |
| **Locomotor type** | | | | | | | | | |
| arboreal | 7 | 1.750 | 1.008 – 2.491 | 0.382 | -0.390 – 1.153 | 1.346 | -0.859 – 3.552 | 0.861 | × |
| semiarboreal | 10 | 1.780 | 1.378 – 2.182 | 0.342 | -0.072 – 0.755 | 1.256 | 0.291 – 2.221 | 0.944 | × |
| scansorial | 45 | 2.889 | 2.714 – 3.063 | 0.254 | 0.135 – 0.373 | 1.286 | 1.034 – 1.537 | 0.976 | ✓ (*D*>1) |
| terrestrial | 49 | 3.349 | 2.947 – 3.751 | 0.566 | 0.295 – 0.836 | 0.879 | 0.685 – 1.072 | 0.957 | × |
| semifossorial | 7 | 1.356 | 1.179 – 1.532 | 0.082 | -0.069 – 0.233 | 2.174 | 0.490 – 3.859 | 0.972 | × |
| semiaquatic | 11 | – | – | – | – | – | – | – | n.c. |
| aquatic | 8 | 2.925 | 2.557 – 3.292 | 0.217 | -0.139 – 0.572 | 1.632 | 0.278 – 2.986 | 0.952 | × |

| SR44 – RR | n | *ln A* | 95% CI$_{\ln A}$ | *C* | 95% CI$_{C}$ | *D* | 95% CI$_{D}$ | R | *D* ≠ 1 |
|---|---|---|---|---|---|---|---|---|---|
| **whole sample** | 137 | -2.792 | -2.856 – -2.728 | $-1.25 \cdot 10^{-7}$ | $-1.85 \cdot 10^{-6}$ – $1.61 \cdot 10^{-6}$ | 7.165 | 0.438 – 13.891 | 0.251 | × |
| **fissipeds** | 130 | -2.838 | -2.912 – -2.763 | $-1.48 \cdot 10^{-5}$ | $-1.37 \cdot 10^{-4}$ – $1.07 \cdot 10^{-4}$ | 5.090 | 0.940 – 9.240 | 0.319 | × |
| **Family** | | | | | | | | | |
| Canidae | *17* | *-2.917* | *-3.140 – -2.693* | *0.005* | *-0.089 – 0.100* | *2.954* | *-11.798 – 17.706* | *0.207* | n.s. |
| Mustelidae | *32* | *-2.477* | *-2.851 – -2.103* | *0.022* | *-0.353 – 0.397* | *0.686* | *-7.398 – 8.769* | *0.089* | n.s. |
| Procyonidae | 7 | – | – | – | – | – | – | – | n.c. |
| Ursidae | 7 | – | – | – | – | – | – | – | n.c. |
| Felidae | 26 | -2.704 | -2.975 – -2.432 | 0.166 | -0.113 – 0.445 | 0.769 | -0.127 – 1.666 | 0.714 | × |
| Herpestidae | 12 | – | – | – | – | – | – | – | n.c. |
| Eupleridae | *5* | *-2.786* | *-3.188 – -2.385* | *0.074* | *-0.506 – 0.655* | *0.908* | *-7.186 – 9.002* | *0.736* | n.s. |
| Viverridae | 14 | -2.678 | -2.858 – -2.499 | 0.088 | -0.124 – 0.300 | 0.665 | -1.106 – 2.437 | 0.556 | × |
| **Locomotor type** | | | | | | | | | |
| arboreal | 7 | – | – | – | – | – | – | – | n.c. |
| semiarboreal | 10 | – | – | – | – | – | – | – | n.c. |
| scansorial | *45* | *-2.742* | *-3.261 – -2.223* | *0.112* | *-0.424 – 0.648* | *0.467* | *-1.287 – 2.221* | *0.266* | n.s. |
| terrestrial | 49 | -2.940 | -3.156 – -2.724 | -0.004 | -0.026 – 0.017 | 2.452 | -0.081 – 4.985 | 0.578 | × |
| semifossorial | 7 | – | – | – | – | – | – | – | n.c. |
| semiaquatic | *11* | *-2.133* | *-2.784 – -1.483* | *0.229* | *-0.469 – 0.928* | *0.674* | *-1.156 – 2.503* | *0.566* | n.s. |
| aquatic | 8 | – | – | – | – | – | – | – | n.c. |

| **SR45 – $L_u$** | **n** | ***ln A*** | **95% $CI_{ln A}$** | ***C*** | **95% $CI_C$** | ***D*** | **95% $CI_D$** | **R** | ***D* ≠ 1** |
|---|---|---|---|---|---|---|---|---|---|
| **whole sample** | 136 | 5.510 | 5.331 – 5.688 | 0.085 | 0.027 – 0.143 | 1.613 | 1.291 – 1.936 | 0.911 | ✓ (*D*>1) |
| **fissipeds** | 129 | 5.736 | 5.499 – 5.972 | 0.216 | 0.092 – 0.339 | 1.235 | 0.972 – 1.497 | 0.919 | × |
| **Family** | | | | | | | | | |
| Canidae | 16 | 5.491 | 5.102 – 5.881 | 0.473 | 0.054 – 0.891 | 0.869 | 0.228 – 1.511 | 0.886 | × |
| Mustelidae | 32 | 4.525 | 4.278 – 4.772 | 0.199 | 0.007 – 0.390 | 1.215 | 0.666 – 1.764 | 0.901 | × |
| Procyonidae | 7 | 4.721 | 4.162 – 5.281 | 0.540 | -0.074 – 1.155 | 0.435 | -0.256 – 1.125 | 0.947 | × |
| Ursidae | 7 | 5.703 | 5.630 – 5.776 | 0.239 | 0.151 – 0.327 | 1.115 | 0.673 – 1.557 | 0.991 | × |
| Felidae | 26 | 5.593 | 5.409 – 5.778 | 0.245 | 0.080 – 0.411 | 1.060 | 0.663 – 1.456 | 0.949 | × |
| Herpestidae | 12 | 4.305 | 4.007 – 4.603 | 0.384 | 0.019 – 0.750 | 0.824 | -0.042 – 1.689 | 0.894 | × |
| Eupleridae | 5 | 4.469 | 3.923 – 5.015 | 0.172 | -0.455 – 0.799 | 1.439 | -2.444 – 5.322 | 0.938 | × |
| Viverridae | 14 | 4.816 | 4.534 – 5.099 | 0.516 | 0.191 – 0.840 | 0.417 | 0.072 – 0.762 | 0.949 | ✓ (*D*<1) |
| **Locomotor type** | | | | | | | | | |
| arboreal | 7 | 4.762 | 4.370 – 5.155 | 0.358 | -0.111 – 0.827 | 0.929 | -0.509 – 2.368 | 0.912 | × |
| semiarboreal | 10 | 4.779 | 4.623 – 4.934 | 0.407 | 0.226 – 0.589 | 0.633 | 0.339 – 0.926 | 0.981 | ✓ (*D*<1) |
| scansorial | 45 | 5.599 | 5.452 – 5.746 | 0.172 | 0.073 – 0.271 | 1.308 | 1.000 – 1.616 | 0.964 | × |
| terrestrial | 48 | 5.986 | 5.573 – 6.400 | 0.258 | 0.060 – 0.456 | 1.219 | 0.883 – 1.554 | 0.947 | × |
| semifossorial | 7 | – | – | – | – | – | – | – | n.c. |
| semiaquatic | 11 | – | – | – | – | – | – | – | n.c. |
| aquatic | 8 | 5.306 | 4.878 – 5.734 | 0.188 | -0.253 – 0.629 | 1.257 | -0.590 – 3.105 | 0.847 | × |

| **SR46 – $d_{su}$** | **n** | ***ln A*** | **95% $CI_{ln A}$** | ***C*** | **95% $CI_C$** | ***D*** | **95% $CI_D$** | **R** | ***D* ≠ 1** |
|---|---|---|---|---|---|---|---|---|---|
| **whole sample** | 136 | 3.585 | 3.328 – 3.843 | 0.442 | 0.260 – 0.624 | 0.901 | 0.731 – 1.071 | 0.948 | × |
| **fissipeds** | 129 | 3.466 | 3.189 – 3.743 | 0.492 | 0.286 – 0.697 | 0.852 | 0.679 – 1.025 | 0.938 | × |
| **Family** | | | | | | | | | |
| Canidae | 16 | 2.289 | 1.984 – 2.594 | 0.378 | 0.071 – 0.685 | 1.167 | 0.529 – 1.804 | 0.918 | × |
| Mustelidae | 32 | 2.429 | 2.191 – 2.667 | 0.407 | 0.185 – 0.629 | 0.880 | 0.593 – 1.167 | 0.949 | × |
| Procyonidae | 7 | 2.100 | 1.546 – 2.655 | 0.247 | -0.720 – 1.214 | 1.498 | -3.882 – 6.877 | 0.784 | × |
| Ursidae | *7* | ***3.249*** | ***2.832 – 3.667*** | ***0.214*** | ***-0.284 – 0.712*** | ***0.852*** | ***-1.712 – 3.416*** | ***0.744*** | n.s. |
| Felidae | 26 | 3.165 | 2.952 – 3.379 | 0.424 | 0.214 – 0.633 | 0.880 | 0.604 – 1.157 | 0.964 | × |
| Herpestidae | 12 | 1.752 | 1.261 – 2.243 | 0.497 | -0.077 – 1.071 | 0.568 | -0.267 – 1.403 | 0.867 | × |
| Eupleridae | 5 | 2.057 | 1.554 – 2.560 | 0.561 | -0.239 – 1.362 | 0.661 | -0.767 – 2.090 | 0.985 | × |
| Viverridae | 14 | – | – | – | – | – | – | – | n.c. |
| **Locomotor type** | | | | | | | | | |
| arboreal | 7 | 2.568 | 1.808 – 3.328 | 0.742 | -0.206 – 1.690 | 0.372 | -0.469 – 1.212 | 0.971 | × |
| semiarboreal | 10 | 2.243 | 2.008 – 2.479 | 0.384 | 0.114 – 0.655 | 0.763 | 0.267 – 1.258 | 0.961 | × |
| scansorial | 45 | 3.229 | 3.016 – 3.443 | 0.304 | 0.135 – 0.474 | 1.089 | 0.802 – 1.377 | 0.961 | × |
| terrestrial | 48 | 3.342 | 2.960 – 3.723 | 0.591 | 0.301 – 0.881 | 0.743 | 0.552 – 0.935 | 0.939 | ✓ (*D*<1) |
| semifossorial | 7 | 2.220 | 1.857 – 2.583 | 0.539 | -0.052 – 1.130 | 0.679 | -0.156 – 1.514 | 0.982 | × |
| semiaquatic | 11 | 2.333 | 2.067 – 2.599 | 0.323 | 0.058 – 0.589 | 1.010 | 0.458 – 1.563 | 0.955 | × |
| aquatic | 8 | 3.351 | 2.823 – 3.879 | 0.185 | -0.329 – 0.699 | 1.596 | -0.693 – 3.885 | 0.873 | × |

| SR47 – $d_{tu}$ | n | *ln A* | 95% $CI_{ln A}$ | *C* | 95% $CI_C$ | *D* | 95% $CI_D$ | R | $D \neq 1$ |
|---|---|---|---|---|---|---|---|---|---|
| **whole sample** | 136 | 3.316 | 3.089 – 3.543 | 0.426 | 0.269 – 0.584 | 0.992 | 0.768 – 1.076 | 0.958 | × |
| **fissipeds** | 129 | 3.370 | 3.108 – 3.632 | 0.585 | 0.382 – 0.788 | 0.798 | 0.657 – 0.939 | 0.953 | ✓ (*D*<1) |
| **Family** | | | | | | | | | |
| Canidae | 16 | 2.392 | 2.067 – 2.717 | 0.443 | 0.093 – 0.793 | 0.861 | 0.291 – 1.432 | 0.906 | × |
| Mustelidae | 32 | 1.887 | 1.639 – 2.134 | 0.238 | 0.037 – 0.439 | 1.141 | 0.667 – 1.615 | 0.915 | × |
| Procyonidae | 7 | 1.677 | 1.041 – 2.314 | 0.436 | -0.262 – 1.134 | 0.432 | -0.535 – 1.400 | 0.903 | × |
| Ursidae | 7 | 3.083 | 2.751 – 3.415 | 0.133 | -0.221 – 0.486 | 2.313 | -1.158 – 5.784 | 0.870 | × |
| Felidae | 26 | 2.974 | 2.654 – 3.293 | 0.545 | 0.223 – 0.867 | 0.815 | 0.491 – 1.139 | 0.945 | × |
| Herpestidae | 12 | 1.618 | 1.269 – 1.968 | 0.422 | -0.002 – 0.846 | 0.741 | -0.126 – 1.607 | 0.884 | × |
| Eupleridae | 5 | 1.687 | 1.516 – 1.858 | 0.358 | 0.124 – 0.592 | 1.010 | 0.310 – 1.710 | 0.997 | × |
| Viverridae | 14 | 1.941 | 1.714 – 2.169 | 0.398 | 0.132 – 0.664 | 0.738 | 0.225 – 1.250 | 0.924 | × |
| **Locomotor type** | | | | | | | | | |
| arboreal | 7 | 1.772 | 1.455 – 2.090 | 0.222 | -0.100 – 0.543 | 1.432 | -0.149 – 3.012 | 0.926 | × |
| semiarboreal | 10 | 1.968 | 1.724 – 2.212 | 0.455 | 0.171 – 0.739 | 0.676 | 0.254 – 1.098 | 0.965 | × |
| scansorial | 45 | 3.157 | 2.877 – 3.437 | 0.512 | 0.258 – 0.766 | 0.866 | 0.627 – 1.106 | 0.962 | × |
| terrestrial | 48 | 3.313 | 2.994 – 3.631 | 0.472 | 0.262 – 0.682 | 0.901 | 0.719 – 1.083 | 0.965 | × |
| semifossorial | 7 | 1.689 | 1.154 – 2.225 | 0.462 | -0.458 – 1.383 | 0.763 | -0.848 – 2.375 | 0.946 | × |
| semiaquatic | 11 | 1.814 | 1.421 – 2.207 | 0.140 | -0.204 – 0.484 | 1.436 | -0.345 – 3.217 | 0.833 | × |
| aquatic | 8 | 3.036 | 2.853 – 3.219 | 0.235 | 0.058 – 0.413 | 1.622 | 0.999 – 2.244 | 0.989 | × |

| SR48 – O | n | *ln A* | 95% $CI_{ln A}$ | *C* | 95% $CI_C$ | *D* | 95% $CI_D$ | R | $D \neq 1$ |
|---|---|---|---|---|---|---|---|---|---|
| **whole sample** | 136 | 4.300 | 4.176 – 4.424 | 0.224 | 0.159 – 0.290 | 1.209 | 1.079 – 1.339 | 0.978 | ✓ (*D*>1) |
| **fissipeds** | 129 | 4.199 | 4.064 – 4.334 | 0.256 | 0.180 – 0.331 | 1.163 | 1.029 – 1.297 | 0.975 | ✓ (*D*>1) |
| **Family** | | | | | | | | | |
| Canidae | 16 | 3.667 | 3.465 – 3.868 | 0.394 | 0.185 – 0.603 | 1.048 | 0.642 – 1.454 | 0.959 | × |
| Mustelidae | 32 | 3.305 | 3.122 – 3.488 | 0.324 | 0.168 – 0.480 | 1.059 | 0.793 – 1.324 | 0.967 | × |
| Procyonidae | 7 | 2.837 | 2.325 – 3.348 | 0.294 | -0.482 – 1.071 | 1.155 | -2.079 – 4.389 | 0.831 | × |
| Ursidae | 7 | 4.395 | 2.809 – 5.982 | 0.608 | -1.027 – 2.242 | 0.225 | -0.664 – 1.113 | 0.927 | × |
| Felidae | 26 | 4.192 | 4.069 – 4.315 | 0.385 | 0.268 – 0.501 | 0.969 | 0.795 – 1.143 | 0.987 | × |
| Herpestidae | 12 | 2.711 | 2.556 – 2.866 | 0.325 | 0.134 – 0.515 | 0.984 | 0.409 – 1.558 | 0.956 | × |
| Eupleridae | 5 | 3.001 | 2.629 – 3.373 | 0.311 | -0.145 – 0.767 | 1.279 | -0.287 – 2.846 | 0.987 | × |
| Viverridae | 14 | 3.117 | 2.951 – 3.283 | 0.314 | 0.138 – 0.489 | 1.141 | 0.670 – 1.612 | 0.956 | × |
| **Locomotor type** | | | | | | | | | |
| arboreal | 7 | 3.207 | 2.815 – 3.600 | 0.379 | -0.063 – 0.822 | 1.099 | -0.184 – 2.381 | 0.934 | × |
| semiarboreal | 10 | 3.109 | 2.942 – 3.438 | 0.395 | 0.121 – 0.669 | 0.990 | 0.465 – 1.515 | 0.972 | × |
| scansorial | 45 | 4.058 | 3.927 – 4.188 | 0.217 | 0.125 – 0.308 | 1.257 | 1.032 – 1.482 | 0.980 | ✓ (*D*>1) |
| terrestrial | 48 | 4.304 | 4.050 – 4.557 | 0.297 | 0.161 – 0.434 | 1.107 | 0.911 – 1.304 | 0.976 | × |
| semifossorial | 7 | 3.184 | 2.952 – 3.416 | 0.403 | -0.013 – 0.820 | 0.979 | 0.063 – 1.894 | 0.985 | × |
| semiaquatic | 11 | 3.257 | 2.953 – 3.562 | 0.256 | -0.029 – 0.541 | 1.232 | 0.449 – 2.014 | 0.944 | × |
| aquatic | 8 | 4.232 | 3.830 – 4.634 | 0.135 | -0.223 – 0.493 | 1.982 | -0.279 – 4.244 | 0.927 | × |

| SR49 – θ | n | *ln A* | 95% CI$_{ln A}$ | *C* | 95% CI$_C$ | *D* | 95% CI$_D$ | R | *D* ≠ 1 |
|---|---|---|---|---|---|---|---|---|---|
| **whole sample** | *136* | *-2.520* | *-3.115 – -1.925* | *-0.009* | *-0.279 – 0.260* | *1.344* | *-11.773 – 14.461* | *0.045* | n.s. |
| **fissipeds** | *129* | *-2.716* | *-3.947 – -1.484* | *-0.105* | *-1.167 – 0.957* | *0.642* | *-3.144 – 4.428* | *0.100* | n.s. |
| **Family** | | | | | | | | | |
| Canidae | 16 | – | – | – | – | – | – | – | n.c. |
| Mustelidae | 32 | – | – | – | – | – | – | – | n.c. |
| Procyonidae | *7* | *-2.082* | *-2.977 – -1.187* | *0.374* | *-1.110 – 1.858* | *1.365* | *-3.892 – 6.622* | *0.753* | n.s. |
| Ursidae | 7 | – | – | – | – | – | – | – | n.c. |
| Felidae | 26 | – | – | – | – | – | – | – | n.c. |
| Herpestidae | 12 | -1.920 | -2.686 – -1.154 | 0.277 | -0.599 – 1.153 | 1.546 | -1.792 – 4.884 | 0.608 | × |
| Eupleridae | 5 | – | – | – | – | – | – | – | n.c. |
| Viverridae | 14 | – | – | – | – | – | – | – | n.c. |
| **Locomotor type** | | | | | | | | | |
| arboreal | 7 | – | – | – | – | – | – | – | n.c. |
| semiarboreal | 10 | – | – | – | – | – | – | – | n.c. |
| scansorial | 45 | – | – | – | – | – | – | – | n.c. |
| terrestrial | *48* | *-3.009* | *-4.259 – -1.759* | *-0.036* | *-0.496 – 0.423* | *1.455* | *-4.256 – 7.166* | *0.207* | n.s. |
| semifossorial | 7 | – | – | – | – | – | – | – | n.c. |
| semiaquatic | 11 | – | – | – | – | – | – | – | n.c. |
| aquatic | 8 | – | – | – | – | – | – | – | n.c. |

| SR50 – α | n | *ln A* | 95% CI$_{ln A}$ | *C* | 95% CI$_C$ | *D* | 95% CI$_D$ | R | *D* ≠ 1 |
|---|---|---|---|---|---|---|---|---|---|
| **whole sample** | 136 | -0.365 | -0.877 – 0.146 | 0.192 | -0.144 – 0.528 | 0.982 | 0.239 – 1.724 | 0.586 | × |
| **fissipeds** | 129 | -0.412 | -1.009 – 0.184 | 0.243 | -0.194 – 0.681 | 0.867 | 0.116 – 1.617 | 0.535 | × |
| **Family** | | | | | | | | | |
| Canidae | 16 | -0.518 | -0.856 – -0.180 | 0.092 | -0.213 – 0.397 | 1.480 | -1.207 – 4.166 | 0.539 | × |
| Mustelidae | 32 | – | – | – | – | – | – | – | n.c. |
| Procyonidae | 7 | -1.193 | -2.295 – -0.092 | 0.630 | -0.741 – 2.000 | 0.712 | -1.264 – 2.687 | 0.817 | × |
| Ursidae | 7 | – | – | – | – | – | – | – | n.c. |
| Felidae | 26 | -0.719 | -1.016 – -0.423 | 0.120 | -0.087 – 0.326 | 1.471 | 0.391 – 2.551 | 0.832 | × |
| Herpestidae | *12* | *-1.087* | *-2.235 – 0.062* | *0.406* | *-0.929 – 1.740* | *0.548* | *-1.764 – 2.859* | *0.526* | n.s. |
| Eupleridae | 5 | – | – | – | – | – | – | – | n.c. |
| Viverridae | 14 | -1.463 | -1.941 – -0.985 | 0.127 | -0.258 – 0.512 | 1.792 | -0.812 – 4.396 | 0.624 | × |
| **Locomotor type** | | | | | | | | | |
| arboreal | 7 | – | – | – | – | – | – | – | n.c. |
| semiarboreal | 10 | – | – | – | – | – | – | – | n.c. |
| scansorial | 45 | -0.636 | -0.910 – -0.362 | 0.137 | -0.056 – 0.330 | 1.251 | 0.502 – 1.999 | 0.826 | × |
| terrestrial | 48 | -0.271 | -1.096 – 0.554 | 0.143 | -0.300 – 0.586 | 1.110 | -0.213 – 2.434 | 0.556 | × |
| semifossorial | 7 | – | – | – | – | – | – | – | n.c. |
| semiaquatic | 11 | – | – | – | – | – | – | – | n.c. |
| aquatic | 8 | – | – | – | – | – | – | – | n.c. |

| SR51 – UR | n | *ln A* | 95% $CI_{ln A}$ | *C* | 95% $CI_C$ | *D* | 95% $CI_D$ | R | *D* ≠ 1 |
|---|---|---|---|---|---|---|---|---|---|
| **whole sample** | 136 | – | – | – | – | – | – | – | n.c. |
| **fissipeds** | 129 | -2.659 | -2.749 – -2.570 | $-5.03 \cdot 10^{-6}$ | $-8.56 \cdot 10^{-5} – 7.55 \cdot 10^{-5}$ | 5.440 | -2.626 – 13.506 | 0.179 | × |
| **Family** | | | | | | | | | |
| Canidae | *16* | *-3.109* | *-3.384 – -2.833* | *0.012* | *-0.123 – 0.146* | *2.720* | *-6.867 – 12.306* | *0.297* | n.s. |
| Mustelidae | *32* | *-1.900* | *-4.729 – 0.929* | *0.409* | *-2.494 – 3.312* | *0.161* | *-1.039 – 1.362* | *0.349* | n.s. |
| Procyonidae | 7 | – | – | – | – | – | – | – | n.c. |
| Ursidae | 7 | – | – | – | – | – | – | – | n.c. |
| Felidae | 26 | -2.400 | -2.966 – -1.835 | 0.209 | -0.414 – 0.832 | 0.469 | -0.797 – 1.736 | 0.475 | × |
| Herpestidae | 12 | – | – | – | – | – | – | – | n.c. |
| Eupleridae | 5 | – | – | – | – | – | – | – | n.c. |
| Viverridae | 14 | – | – | – | – | – | – | – | n.c. |
| **Locomotor type** | | | | | | | | | |
| arboreal | 7 | – | – | – | – | – | – | – | n.c. |
| semiarboreal | *10* | *-2.497* | *-2.756 – -2.239* | *0.027* | *-0.244 – 0.299* | *1.193* | *-6.714 – 9.100* | *0.315* | n.s. |
| scansorial | 45 | -2.336 | -2.914 – -1.758 | 0.182 | -0.415 – 0.779 | 0.463 | -0.733 – 1.658 | 0.374 | × |
| terrestrial | 48 | -3.039 | -3.337 – -2.740 | -0.004 | -0.028 – 0.020 | 2.605 | -0.232 – 5.441 | 0.551 | × |
| semifossorial | 7 | – | – | – | – | – | – | – | n.c. |
| semiaquatic | *11* | *-1.965* | *-2.772 – -1.157* | *0.287* | *-0.606 – 1.181* | *0.412* | *-1.062 – 1.886* | *0.576* | *n.s.* |
| aquatic | 8 | – | – | – | – | – | – | – | n.c. |

| SR52 – IFA | n | *ln A* | 95% $CI_{ln A}$ | *C* | 95% $CI_C$ | *D* | 95% $CI_D$ | R | *D* ≠ 1 |
|---|---|---|---|---|---|---|---|---|---|
| **whole sample** | 136 | -0.422 | -4.259 – 3.416 | 0.992 | -2.882 – 4.866 | 0.114 | -0.315 – 0.543 | 0.341 | ✓ (*D*=0) |
| **fissipeds** | *129* | *-1.358* | *-10.852 – 8.137* | *0.235* | *-9.312 – 9.782* | *0.068* | *-2.713 – 2.850* | *0.055* | n.s. |
| **Family** | | | | | | | | | |
| Canidae | *16* | *-1.753* | *-1.917 – -1.589* | *0.001* | *-0.018 – 0.019* | *4.920* | *-23.901 – 33.741* | *0.237* | n.s. |
| Mustelidae | 32 | -1.243 | -1.575 – -0.910 | 0.104 | -0.216 – 0.425 | 0.795 | -0.762 – 2.352 | 0.459 | × |
| Procyonidae | 7 | – | – | – | – | – | – | – | n.c. |
| Ursidae | 7 | – | – | – | – | – | – | – | n.c. |
| Felidae | 26 | -1.398 | -1.620 – -1.176 | 0.142 | -0.086 – 0.370 | 0.765 | -0.088 – 1.618 | 0.730 | × |
| Herpestidae | 12 | – | – | – | – | – | – | – | n.c. |
| Eupleridae | 5 | -1.458 | -1.856 – -1.059 | 0.162 | -0.421 – 0.745 | 0.881 | -2.828 – 4.589 | 0.920 | × |
| Viverridae | 14 | -1.531 | -1.628 – -1.435 | 0.023 | -0.025 – 0.071 | 2.685 | 0.930 – 4.440 | 0.867 | × |
| **Locomotor type** | | | | | | | | | |
| arboreal | 7 | – | – | – | – | – | – | – | n.c. |
| semiarboreal | 10 | -1.557 | -1.705 – -1.409 | 0.036 | -0.060 – 0.132 | 2.199 | -0.092 – 4.491 | 0.891 | × |
| scansorial | 45 | -1.552 | -1.718 – -1.386 | 0.036 | -0.093 – 0.166 | 1.116 | -0.750 – 2.982 | 0.480 | × |
| terrestrial | 48 | -1.763 | -2.028 – -1.498 | -0.010 | -0.084 – 0.064 | 1.683 | -1.743 – 5.109 | 0.374 | × |
| semifossorial | 7 | – | – | – | – | – | – | – | n.c. |
| semiaquatic | 11 | -1.047 | -1.482 – -0.613 | 0.217 | -0.251 – 0.685 | 0.655 | -0.628 – 1.938 | 0.693 | × |
| aquatic | 8 | -1.027 | -1.156 – -0.899 | 0.009 | -0.045 – 0.062 | 3.696 | -1.901 – 9.294 | 0.911 | × |

| SR53 – $L_m$ | n | *ln A* | 95% $CI_{ln A}$ | *C* | 95% $CI_C$ | *D* | 95% $CI_D$ | R | *D* ≠ 1 |
|---|---|---|---|---|---|---|---|---|---|
| **whole sample** | 136 | 4.449 | 4.254 – 4.644 | 0.080 | 0.017 – 0.143 | 1.620 | 1.248 – 1.993 | 0.887 | ✓ (*D*>1) |
| **fissipeds** | 129 | 4.566 | 4.323 – 4.810 | 0.157 | 0.045 – 0.268 | 1.356 | 1.023 – 1.690 | 0.890 | ✓ (*D*>1) |
| **Family** | | | | | | | | | |
| Canidae | 16 | 4.589 | 4.290 – 4.888 | 0.454 | 0.135 – 0.772 | 0.930 | 0.409 – 1.451 | 0.926 | × |
| Mustelidae | 32 | 3.477 | 3.236 – 3.718 | 0.127 | -0.038 – 0.292 | 1.395 | 0.636 – 2.153 | 0.857 | × |
| Procyonidae | 7 | 3.529 | 2.596 – 4.462 | 0.513 | -0.461 – 1.487 | 0.311 | -0.513 – 1.134 | 0.912 | × |
| Ursidae | 7 | 4.505 | 4.156 – 4.855 | 0.409 | -0.003 – 0.821 | 0.707 | -0.315 – 1.730 | 0.931 | × |
| Felidae | 26 | 4.635 | 4.478 – 4.792 | 0.239 | 0.103 – 0.375 | 1.135 | 0.796 – 1.473 | 0.966 | × |
| Herpestidae | 12 | 3.492 | 3.220 – 3.764 | 0.512 | 0.188 – 0.836 | 0.642 | 0.142 – 1.141 | 0.951 | × |
| Eupleridae | 5 | 3.315 | 3.038 – 3.591 | 0.106 | -0.169 – 0.382 | 1.774 | -0.979 – 4.527 | 0.976 | × |
| Viverridae | 14 | 3.593 | 3.365 – 3.821 | 0.340 | 0.082 – 0.597 | 0.927 | 0.309 – 1.545 | 0.911 | × |
| **Locomotor type** | | | | | | | | | |
| arboreal | 7 | – | – | – | – | – | – | – | n.c. |
| semiarboreal | 10 | 3.699 | 3.142 – 4.256 | 0.430 | -0.221 – 1.081 | 0.620 | -0.373 – 1.613 | 0.830 | × |
| scansorial | 45 | 4.384 | 4.199 – 4.569 | 0.098 | 0.002 – 0.195 | 1.579 | 1.035 – 2.123 | 0.918 | ✓ (*D*>1) |
| terrestrial | 48 | 4.942 | 4.531 – 5.353 | 0.200 | 0.024 – 0.375 | 1.325 | 0.935 – 1.715 | 0.942 | × |
| semifossorial | 7 | – | – | – | – | – | – | – | n.c. |
| semiaquatic | 11 | 3.716 | 3.269 – 4.164 | 0.369 | -0.115 – 0.852 | 0.639 | -0.134 – 1.412 | 0.844 | × |
| aquatic | 8 | 4.643 | 4.250 – 5.036 | 0.523 | 0.100 – 0.945 | 0.916 | 0.319 – 1.514 | 0.960 | × |

| SR54 – $d_{sm}$ | n | *ln A* | 95% $CI_{ln A}$ | *C* | 95% $CI_C$ | *D* | 95% $CI_D$ | R | *D* ≠ 1 |
|---|---|---|---|---|---|---|---|---|---|
| **whole sample** | 136 | 2.692 | 2.524 – 2.860 | 0.283 | 0.180 – 0.385 | 1.060 | 0.903 – 1.217 | 0.963 | × |
| **fissipeds** | 129 | 2.575 | 2.397 – 2.753 | 0.296 | 0.185 – 0.407 | 1.050 | 0.884 – 1.216 | 0.957 | × |
| **Family** | | | | | | | | | |
| Canidae | 16 | 1.987 | 1.781 – 2.194 | 0.291 | 0.099 – 0.483 | 1.396 | 0.861 – 1.931 | 0.951 | × |
| Mustelidae | 32 | 1.717 | 1.482 – 1.952 | 0.339 | 0.125 – 0.553 | 0.933 | 0.595 – 1.270 | 0.937 | × |
| Procyonidae | 7 | 1.248 | 0.595 – 1.901 | 0.325 | -0.498 – 1.149 | 0.742 | -1.622 – 3.105 | 0.776 | × |
| Ursidae | 7 | 2.385 | 2.200 – 2.571 | 0.197 | -0.024 – 0.419 | 0.887 | -0.372 – 2.146 | 0.918 | × |
| Felidae | 26 | 2.454 | 2.348 – 2.560 | 0.320 | 0.223 – 0.418 | 1.021 | 0.844 – 1.198 | 0.988 | × |
| Herpestidae | 12 | 1.223 | 0.979 – 1.446 | 0.322 | 0.026 – 0.618 | 0.754 | -0.046 – 1.554 | 0.901 | × |
| Eupleridae | 5 | 1.196 | 0.616 – 1.776 | 0.248 | -0.379 – 0.875 | 1.586 | -1.106 – 4.278 | 0.973 | × |
| Viverridae | 14 | 1.453 | 1.204 – 1.703 | 0.345 | 0.058 – 0.631 | 0.848 | 0.183 – 1.514 | 0.892 | × |
| **Locomotor type** | | | | | | | | | |
| arboreal | 7 | 1.348 | 0.997 – 1.699 | 0.162 | -0.138 – 0.463 | 1.939 | -0.073 – 3.952 | 0.928 | × |
| semiarboreal | 10 | 1.700 | 1.326 – 2.075 | 0.573 | 0.133 – 1.013 | 0.589 | 0.096 – 1.081 | 0.946 | × |
| scansorial | 45 | 2.370 | 2.254 – 2.487 | 0.174 | 0.099 – 0.250 | 1.358 | 1.125 – 1.591 | 0.980 | ✓ (*D*>1) |
| terrestrial | 48 | 2.587 | 2.251 – 2.923 | 0.303 | 0.108 – 0.498 | 1.034 | 0.763 – 1.305 | 0.948 | × |
| semifossorial | 7 | – | – | – | – | – | – | – | n.c. |
| semiaquatic | 11 | 1.797 | 1.445 – 2.149 | 0.343 | -0.011 – 0.696 | 0.982 | 0.293 – 1.670 | 0.929 | × |
| aquatic | 8 | 2.806 | 2.672 – 2.940 | 0.454 | 0.313 – 0.596 | 1.054 | 0.817 – 1.291 | 0.995 | × |

| **SR55 – $d_{tm}$** | **n** | ***ln A*** | **95% CI$_{ln A}$** | ***C*** | **95% CI$_C$** | ***D*** | **95% CI$_D$** | **R** | ***D* ≠ 1** |
|---|---|---|---|---|---|---|---|---|---|
| **whole sample** | 136 | 2.709 | 2.553 – 2.865 | 0.251 | 0.161 – 0.342 | 1.115 | 0.957 – 1.273 | 0.965 | × |
| **fissipeds** | 129 | 2.670 | 2.499 – 2.841 | 0.304 | 0.197 – 0.412 | 1.048 | 0.892 – 1.203 | 0.962 | × |
| **Family** | | | | | | | | | |
| Canidae | 16 | 2.142 | 1.839 – 2.444 | 0.438 | 0.115 – 0.760 | 0.929 | 0.383 – 1.475 | 0.920 | × |
| Mustelidae | 32 | 1.825 | 1.626 – 2.024 | 0.347 | 0.169 – 0.526 | 0.966 | 0.689 – 1.243 | 0.959 | × |
| Procyonidae | *7* | *1.204* | *0.719 – 1.689* | *0.217* | *-0.382 – 0.817* | *0.699* | *-1.773 – 3.171* | *0.744* | n.s. |
| Ursidae | 7 | 2.469 | 2.297 – 2.641 | 0.215 | 0.009 – 0.422 | 1.047 | -0.083 – 2.178 | 0.944 | × |
| Felidae | 26 | 2.580 | 2.393 – 2.766 | 0.372 | 0.195 – 0.548 | 0.961 | 0.689 – 1.233 | 0.970 | × |
| Herpestidae | 12 | 1.315 | 1.046 – 1.584 | 0.375 | 0.055 – 0.695 | 0.632 | -0.033 – 1.297 | 0.917 | × |
| Eupleridae | 5 | 1.342 | 0.713 – 1.972 | 0.324 | -0.481 – 1.130 | 1.171 | -1.486 – 3.828 | 0.962 | × |
| Viverridae | 14 | 1.505 | 1.309 – 1.702 | 0.363 | 0.136 – 0.590 | 0.821 | 0.325 – 1.317 | 0.933 | × |
| **Locomotor type** | | | | | | | | | |
| arboreal | 7 | 1.465 | 1.139 – 1.791 | 0.278 | -0.077 – 0.633 | 1.206 | -0.193 – 2.606 | 0.929 | × |
| semiarboreal | 10 | 1.709 | 1.398 – 2.019 | 0.495 | 0.138 – 0.852 | 0.743 | 0.239 – 1.247 | 0.958 | × |
| scansorial | 45 | 2.491 | 2.334 – 2.649 | 0.234 | 0.118 – 0.349 | 1.197 | 0.936 – 1.459 | 0.971 | × |
| terrestrial | 48 | 2.695 | 2.407 – 2.983 | 0.283 | 0.124 – 0.441 | 1.087 | 0.848 – 1.326 | 0.963 | × |
| semifossorial | 7 | 1.782 | 0.735 – 2.829 | 0.757 | -0.585 – 2.098 | 0.424 | -0.521 – 1.369 | 0.958 | × |
| semiaquatic | 11 | 1.937 | 1.427 – 2.447 | 0.592 | 0.032 – 1.153 | 0.538 | 0.018 – 1.059 | 0.907 | × |
| aquatic | 8 | 2.837 | 2.513 – 3.160 | 0.523 | 0.167 – 0.878 | 0.717 | 0.241 – 1.192 | 0.962 | × |

| **SR56 – MR** | **n** | ***ln A*** | **95% CI$_{ln A}$** | ***C*** | **95% CI$_C$** | ***D*** | **95% CI$_D$** | **R** | ***D* ≠ 1** |
|---|---|---|---|---|---|---|---|---|---|
| **whole sample** | 136 | – | – | – | – | – | – | – | n.c. |
| **fissipeds** | *129* | *-2.218* | *-2.285 – -2.152* | *$-1.25 \cdot 10^{-6}$* | *$-2.47 \cdot 10^{-5} – 2.22 \cdot 10^{-5}$* | *5.970* | *-3.414 – 15.354* | *0.164* | n.s. |
| **Family** | | | | | | | | | |
| Canidae | 16 | – | – | – | – | – | – | – | n.c. |
| Mustelidae | 32 | -1.700 | -2.840 – -0.560 | 0.283 | -0.919 – 1.484 | 0.287 | -0.923 – 1.498 | 0.370 | × |
| Procyonidae | 7 | – | – | – | – | – | – | – | n.c. |
| Ursidae | 7 | – | – | – | – | – | – | – | n.c. |
| Felidae | *26* | *-2.197* | *-2.495 – -1.899* | *0.064* | *-0.255 – 0.383* | *0.645* | *-1.841 – 3.131* | *0.310* | n.s. |
| Herpestidae | 12 | – | – | – | – | – | – | – | n.c. |
| Eupleridae | *5* | *-2.104* | *-2.858 – -1.350* | *0.169* | *-0.779 – 1.118* | *1.225* | *-4.711 – 7.160* | *0.853* | n.s. |
| Viverridae | 14 | – | – | – | – | – | – | – | n.c. |
| **Locomotor type** | | | | | | | | | |
| arboreal | 7 | – | – | – | – | – | – | – | n.c. |
| semiarboreal | *10* | *-2.003* | *-2.841 – -1.164* | *0.137* | *-0.852 – 1.126* | *0.523* | *-3.848 – 4.894* | *0.295* | n.s. |
| scansorial | 45 | -1.965 | -2.372 – -1.559 | 0.138 | -0.272 – 0.547 | 0.582 | -0.639 – 1.804 | 0.436 | × |
| terrestrial | 48 | -2.507 | -2.740 – -2.274 | -0.006 | -0.036 – 0.023 | 2.279 | -0.054 – 4.613 | 0.595 | × |
| semifossorial | 7 | – | – | – | – | – | – | – | n.c. |
| semiaquatic | 11 | -1.860 | -2.037 – -1.682 | 0.050 | -0.092 – 0.192 | 1.668 | -0.452 – 3.788 | 0.833 | × |
| aquatic | 8 | – | – | – | – | – | – | – | n.c. |

| SR57 – %prox | n | *ln A* | 95% CI$_{ln A}$ | *C* | 95% CI$_C$ | *D* | 95% CI$_D$ | R | *D* ≠ 1 |
|---|---|---|---|---|---|---|---|---|---|
| **whole sample** | 137 | 3.702 | 3.217 – 4.187 | 0.271 | -0.218 – 0.760 | 0.229 | -0.129 – 0.587 | 0.469 | ✓ (*D*=0) |
| **fissipeds** | 130 | 3.355 | 3.241 – 3.469 | 0.014 | -0.075 – 0.104 | 0.785 | -1.706 – 3.275 | 0.173 | × |
| **Family** | | | | | | | | | |
| Canidae | *17* | *3.307* | *3.232 – 3.381* | *0.006* | *-0.032 – 0.044* | *2.613* | *-2.580 – 7.805* | *0.459* | n.s. |
| Mustelidae | 32 | 3.724 | 2.566 – 4.882 | 0.345 | -0.840 – 1.531 | 0.152 | -0.397 – 0.701 | 0.630 | ✓ (*D*=0) |
| Procyonidae | 7 | – | – | – | – | – | – | – | n.c. |
| Ursidae | 7 | – | – | – | – | – | – | – | n.c. |
| Felidae | 26 | 3.412 | 3.224 – 3.601 | 0.082 | -0.125 – 0.290 | 0.374 | -0.553 – 1.301 | 0.563 | × |
| Herpestidae | 12 | – | – | – | – | – | – | – | n.c. |
| Eupleridae | 5 | – | – | – | – | – | – | – | n.c. |
| Viverridae | 14 | – | – | – | – | – | – | – | n.c. |
| **Locomotor type** | | | | | | | | | |
| arboreal | 7 | – | – | – | – | – | – | – | n.c. |
| semiarboreal | 10 | – | – | – | – | – | – | – | n.c. |
| scansorial | 45 | 3.366 | 3.252 – 3.480 | 0.041 | -0.075 – 0.158 | 0.521 | -0.576 – 1.618 | 0.438 | × |
| terrestrial | 49 | – | – | – | – | – | – | – | n.c. |
| semifossorial | 7 | – | – | – | – | – | – | – | n.c. |
| semiaquatic | 11 | – | – | – | – | – | – | – | n.c. |
| aquatic | 8 | – | – | – | – | – | – | – | n.c. |

| SR58 – %mid | n | *ln A* | 95% CI$_{ln A}$ | *C* | 95% CI$_C$ | *D* | 95% CI$_D$ | R | *D* ≠ 1 |
|---|---|---|---|---|---|---|---|---|---|
| **whole sample** | 137 | 3.212 | 2.813 – 3.611 | -0.330 | -0.734 – 0.073 | 0.205 | -0.017 – 0.427 | 0.632 | ✓ (*D*=0) |
| **fissipeds** | 130 | 3.631 | 3.604 – 3.658 | -0.001 | -0.004 – 0.002 | 2.449 | 0.669 – 4.229 | 0.462 | × |
| **Family** | | | | | | | | | |
| Canidae | 17 | 3.571 | 3.521 – 3.620 | -0.034 | -0.088 – 0.019 | 0.836 | -0.281 – 1.953 | 0.714 | × |
| Mustelidae | 32 | 3.616 | 3.530 – 3.703 | -0.068 | -0.159 – 0.024 | 0.450 | -0.076 – 0.976 | 0.727 | ✓ (*D*=0) |
| Procyonidae | 7 | – | – | – | – | – | – | – | n.c. |
| Ursidae | 7 | – | – | – | – | – | – | – | n.c. |
| Felidae | *26* | *3.633* | *3.576 – 3.690* | *-0.007* | *-0.068 – 0.053* | *0.695* | *-3.588 – 4.979* | *0.195* | n.s. |
| Herpestidae | 12 | – | – | – | – | – | – | – | n.c. |
| Eupleridae | 5 | – | – | – | – | – | – | – | n.c. |
| Viverridae | 14 | – | – | – | – | – | – | – | n.c. |
| **Locomotor type** | | | | | | | | | |
| arboreal | 7 | – | – | – | – | – | – | – | n.c. |
| semiarboreal | 10 | – | – | – | – | – | – | – | n.c. |
| scansorial | 45 | – | – | – | – | – | – | – | n.c. |
| terrestrial | 49 | 3.564 | 3.509 – 3.618 | -0.004 | -0.015 – 0.007 | 1.953 | 0.664 – 3.241 | 0.767 | × |
| semifossorial | 7 | – | – | – | – | – | – | – | n.c. |
| semiaquatic | 11 | – | – | – | – | – | – | – | n.c. |
| aquatic | 8 | – | – | – | – | – | – | – | n.c. |

| SR59 – %$_{dist}$ | n | *ln A* | 95% CI$_{ln A}$ | *C* | 95% CI$_C$ | *D* | 95% CI$_D$ | R | $D \neq 1$ |
|---|---|---|---|---|---|---|---|---|---|
| **whole sample** | 137 | 3.516 | 3.497 – 3.535 | $3.27 \cdot 10^{-7}$ | $-2.79 \cdot 10^{-6} – 3.44 \cdot 10^{-6}$ | 6.190 | 1.549 – 10.832 | 0.318 | ✓ (*D*>1) |
| **fissipeds** | 130 | 3.519 | 3.498 – 3.540 | $2.68 \cdot 10^{-6}$ | $-1.93 \cdot 10^{-5} – 2.46 \cdot 10^{-5}$ | 5.354 | 1.226 – 9.482 | 0.329 | ✓ (*D*>1) |
| **Family** | | | | | | | | | |
| Canidae | *17* | *3.599* | *3.301 – 3.897* | *0.012* | *-0.308 – 0.332* | *0.372* | *-11.387 – 12.132* | *0.071* | n.s. |
| Mustelidae | *32* | *3.425* | *3.400 – 3.450* | $4.38 \cdot 10^{-8}$ | $-1.98 \cdot 10^{-6} – 2.07 \cdot 10^{-6}$ | *8.423* | *-19.191 – 36.036* | *0.235* | n.s. |
| Procyonidae | 7 | – | – | – | – | – | – | – | n.c. |
| Ursidae | 7 | – | – | – | – | – | – | – | n.c. |
| Felidae | *26* | *3.452* | *3.115 – 3.789* | *-0.070* | *-0.433 – 0.293* | *0.278* | *-1.231 – 1.787* | *0.363* | n.s. |
| Herpestidae | 12 | – | – | – | – | – | – | – | n.c. |
| Eupleridae | 5 | – | – | – | – | – | – | – | n.c. |
| Viverridae | 14 | – | – | – | – | – | – | – | n.c. |
| **Locomotor type** | | | | | | | | | |
| arboreal | 7 | – | – | – | – | – | – | – | n.c. |
| semiarboreal | 10 | – | – | – | – | – | – | – | n.c. |
| scansorial | *45* | *3.481* | *3.356 – 3.607* | *-0.029* | *-0.157 – 0.100* | *0.485* | *-1.202 – 2.171* | *0.285* | n.s. |
| terrestrial | 49 | 3.598 | 3.533 – 3.664 | 0.002 | -0.005 – 0.009 | 2.400 | 0.574 – 4.226 | 0.696 | × |
| semifossorial | 7 | – | – | – | – | – | – | – | n.c. |
| semiaquatic | 11 | – | – | – | – | – | – | – | n.c. |
| aquatic | 8 | 3.687 | 3.034 – 4.339 | 0.208 | -0.479 – 0.896 | 0.249 | -0.821 – 1.318 | 0.832 | × |
